# Interpretable and scalable spatial gene set activity analysis with GESSO uncovers functional tissue architecture

**DOI:** 10.64898/2026.07.02.736099

**Authors:** Andrew Yang, Chichun Tan, Ying Ma

## Abstract

Recent advances in spatially resolved transcriptomics (SRT) enabled measurement of sets of pathway genes activity within tissues. However, existing gene set activity scoring methods overlook spatial dependencies among tissue locations, restricting their ability to capture region-specific pathway activities associated with disease pathology or cellular communication. Moreover, these methods lack significance-level inference for activity scores, provide limited interpretability of gene-level contribution to a pathway, and scale poorly to advanced large-size SRT datasets. To address these limitations, we present GESSO (Gene sEt activity Score analysis with Spatial lOcation), a spatially informed gene set scoring method adaptable to diverse SRT platforms. GESSO models gene set activity levels through a graph-regularized matrix decomposition algorithm, jointly inferring spatially coherent gene set activity scores (GASs) and interpretable metagene weights that capture gene-level contributions. It further implements a permutation-based local significance test and a stratified low-resolution approximation that scales to high-resolution SRT datasets such as Visium HD, Stereo-seq, and Xenium Prime. Across 13 datasets from five SRT platforms, GESSO outperformed all existing methods in accuracy, calibration, interpretability, and scalability. Applications revealed novel biological programs, including spatially confined EMT activation within tumor-stroma interfaces, developmental signaling gradients across embryonic tissues, and coordinated B-cell, T-cell, and signaling pathways within germinal centers of human lymph node tissue, revealing the spatial organization of immune function at subregional resolution.

## Introduction

Gene set activity analysis (also referred to as pathway activity analysis) provides a principled framework for quantifying how predefined groups of genes act as coordinated functional units across biological systems. By aggregating expression at the pathway level, these approaches enable systematic characterization of cellular functions underlying disease progression, tissue differentiation, and developmental trajectories^1–3^. Gene set activity scoring methods have been widely applied to bulk RNA sequencing (RNA-seq) and single-cell RNA-seq (scRNA-seq) data, revealing key mechanisms of tumor pathogenesis^4–6^, immune response^7–9^ and organismal development^10,11^. However, these analyses lack spatial context: bulk RNA-seq averages signals across mixed cell populations, and scRNA-seq dissociates cells, erasing their native spatial organization. This assumption overlooks the fact that molecular activities are structured by tissue architecture. For example, the tumor microenvironment^12,13^, cellular niche organization^14,15^, and cell-cell proximity^16,17^ modulate pathway activation, while spatial gradients and regional boundaries^18,19^ often define functional transitions. Collectively, these studies^20,21^ highlight that spatial organization shapes the activation and coordination of molecular pathways, underscoring the need to integrate spatial context into pathway activity inference.

Spatially resolved transcriptomics (SRT) now enables simultaneous measurement of gene expression and spatial coordinates, providing unprecedented opportunities to dissect molecular processes within their spatial context. This has led to discoveries in cell type localization^22–24^ and spatially variable gene identification^25,26^ and the delineation of tissue domains^27–29^. Despite these advances, pathway-level analysis in SRT data remains underdeveloped. Most existing gene set scoring methods, including single-sample GSEA (ssGSEA)^6^, GSVA^30^, and AUCell^31^, were originally developed for bulk ^32,33^ or scRNA-seq data^34–36^, where each sample or cell is treated independently. Consequently, they cannot capture spatial correlations among neighboring locations and fail to recover region-specific pathway activity signals which are associated to complex biological processes within a tissue. Recent work has begun exploring pathway analysis in SRT, most notably GSDensity^37^, which co-embeds genes and cells via multiple correspondence analysis and estimates pathway activity through network propagation on a cell-gene graph. While this method implicitly reflects spatial organization encoded within cell-cell relationships, it relies on expression covariance rather than explicitly modeling spatial dependencies derived from spatial coordinates. Furthermore, GSDensity lacks rigorous calibration to ensure false-positive control under the null and becomes computationally infeasible for high-resolution datasets. More broadly, current approaches rely on heuristic enrichment scores without formal statistical testing or calibration under the null, providing no principled ways to infer significance-level of detected pathway activity signals locally. They also scale poorly to the increasingly large high-resolution SRT datasets now encompassing hundreds of thousands of spatial coordinates (for example, Visium HD, Stereo-seq, and Xenium Prime). These limitations motivate the development of a unified framework that directly integrates spatial structure with interpretable pathway inference and well-calibrated statistical testing.

To address these limitations, we developed GESSO (Gene sEt activity Score analysis with Spatial lOcation), a spatially informed computational method for quantifying pathway activity in SRT data. GESSO models pathway activity as a spatially coherent latent process through a graph Laplacian-regularized rank-one decomposition, yielding both spatially resolved gene set activity scores (GASs) and metagene weights that quantify gene-level contributions to each pathway. This formulation anchors inference in known gene sets while enforcing spatial smoothness to capture coherent biological structure. Beyond estimation, GESSO introduces a permutation-based local activity-enrichment test that provides location-specific p-values and ensures robust false-positive control at both global and local spatial scales. To enable scalable analyses for high-resolution SRT data, GESSO incorporates a stratified low-resolution approximation that preserves concordance with full-resolution inference while substantially reducing runtime and memory demands, thereby extending its applicability to massive, high-resolution datasets including Visium HD and Xenium Prime.

Through extensive simulations and analyses of 13 spatial transcriptomics datasets spanning five platforms, GESSO outperformed all existing methods in accuracy, interpretability, calibration, and scalability. It accurately recovered region-specific pathway activities across diverse conditions, generated biologically coherent activity maps aligned with known tissue structures, and revealed new biological insights across systems. In human tumors, GESSO delineated oncogenic and immune signaling across distinct tumor microenvironmental niches; in mouse embryos, it captured organ-specific developmental programs and their temporal dynamics; and in human lymphoid tissue, it uncovered spatial coordination between B-cell, T-cell, and intercellular transport pathways within germinal centers. By unifying spatial modeling, pathway-level interpretability, and rigorous inference in a scalable framework, GESSO establishes a new paradigm for spatially resolved functional genomics.

## Results

### Method overview

GESSO (Gene sEt activity Score analysis with Spatial lOcation) is described in Methods, with technical details provided in Supplementary Note 1 and its method schematic illustrated in Figure 1. Briefly, GESSO is a spatially informed framework for quantifying gene set activity in SRT data. Given a gene-by-location expression matrix from SRT studies and a collection of predefined gene sets, GESSO returns for each gene set a GAS at every spatial location and a corresponding metagene vector that summarizes the relative contribution of individual genes (Figure 1, panels 1-2). GESSO builds on the principle that spatially coherent molecular programs underlie tissue organization and therefore incorporates spatial information through a graph Laplacian-regularized rank-one decomposition. This design enables GESSO to capture localized biological functions that align with tissue structure while reducing noise from randomly expressed genes. To assess statistical significance, GESSO implements a local activity-enrichment test, comparing the observed GAS at each location against a null distribution derived from spatially permuted data. To support increasingly large SRT datasets containing hundreds of thousands of spatial locations (for example, Stereo-seq, Visium HD, and Xenium Prime), GESSO includes an accelerated low-resolution mode that achieves high concordance with the standard computation while substantially reducing runtime and memory cost. The resulting outputs include spatial activity maps, metagene weight profiles, and significance maps, which collectively enable analyses such as functional-region delineation, cross-tumor comparison, colocalization of gene set activities, activity change in a local spatial neighborhood, and spatiotemporal characterization of tissue development (Figure 1, panels 3-4).

**Figure 1.**
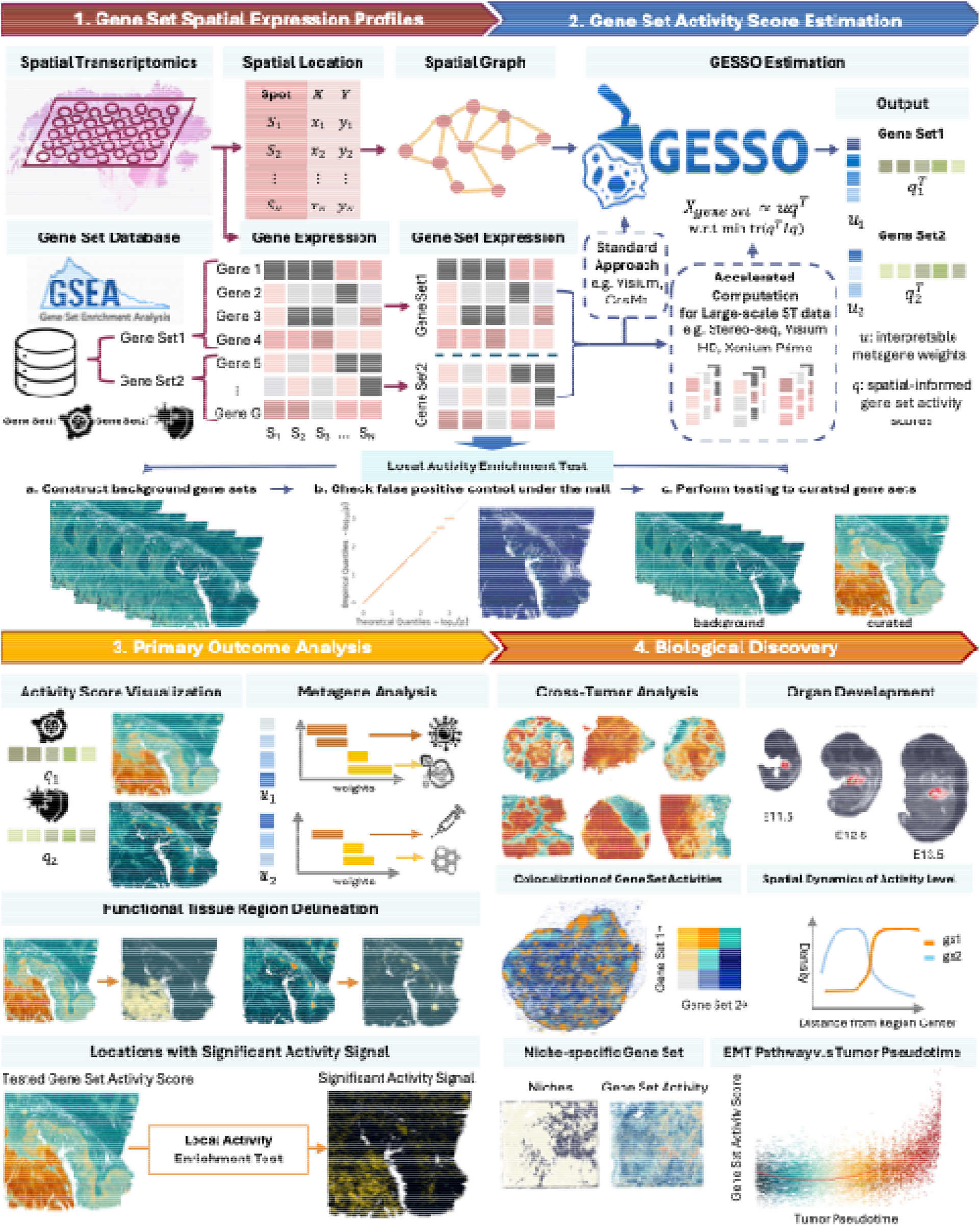
Method overview. An overview of the Gene sEt activity Score analysis with Spatial lOcation (GESSO) method. Taking a spatially resolved gene expression dataset and curated gene set information as inputs (Panel 1), GESSO uses graph-regularized rank-one matrix decomposition to generate a unique GAS for each spatial location. In addition to GASs, GESSO returns a metagene, a vector of importance weights for individual genes within a given pathway (Panel 2). A local activity enrichment testing procedure performs a statistical hypothesis testing to infer the significance of activity signal and return a *p* value at each location. The GASs and the locations with significant signal can be used to delineate functional regional and the metagene weights can be used to infer the most active and inactive genes in the pathways (Panel 3). Further downstream analysis with these three primary outcomes could provide novel biological insights for the input data set (Panel 4).

Throughout the following sections, gene set denotes a simulated or generic group of genes. In contrast, pathway refers to a curated gene set defined by its specific biological context and functional relevance.

### GESSO consistently outperforms competing methods in accuracy and calibration across simulation settings

To systematically evaluate performance, we developed a simulation framework comparing GESSO, AUCell, GSVA, ssGSEA, and GSDensity (Methods). Briefly, using the gene-level mean and standard deviation of expression derived from the human dorsolateral prefrontal cortex (DLPFC) reference data^38^, we simulated expression values for two groups of genes: genes with spatial expression patterns (SGs), which exhibit high expression in the white matter (WM) region, and genes with random expression patterns (RGs), which are randomly expressed across the tissue. We introduced sparsity to match real SRT platforms, ranging from ~60% for 10x Visium to 80-90% for Stereo-seq, Visium HD, Xenium Prime, and CosMx (Supplementary Tables 1-2). Gene sets of size 50, 100, or 200 were generated by mixing SGs and RGs at varying proportions (15%, 30%, 50%, or 70% SGs). Effective scoring methods should yield higher activity in the WM region for gene sets with larger proportion of SGs while controlling false positives for gene sets with RGs. Performance was quantified by the area under the receiver operating characteristic curve (AUC), with Accuracy and F1 as secondary metrics (Methods).

Across all conditions, GESSO consistently achieved the highest AUCs (Figure 2A). Specifically, under moderate sparsity levels (60-70%), GESSO achieved mean AUCs over 0.9 for all gene set settings. When SGs comprised 70% of a gene set, all methods showed improved performance; however, GESSO achieved perfect performance (AUC close to 1). The advantage of GESSO became more pronounced at lower signal levels: with 50% or fewer SG genes, GESSO maintained AUCs close to 1 across all sparsity levels, whereas GSDensity dropped to 0.8 or lower, and AUCell, GSVA, and ssGSEA performed substantially worse (Supplementary Table 3). Importantly, even under high sparsity levels (80-90%), GESSO continued to outperform all the other methods, consistently yielding the highest AUC across gene-set sizes and SGs proportions. Comparable superiority of GESSO was observed when evaluated by accuracy and F1 score (Supplementary Figure S1). To evaluate the contribution of its spatial graph regularization, we performed a sensitivity analysis within GESSO, which confirmed that incorporating spatial structure substantially improves predictive performance (Methods).

**Figure 2.**
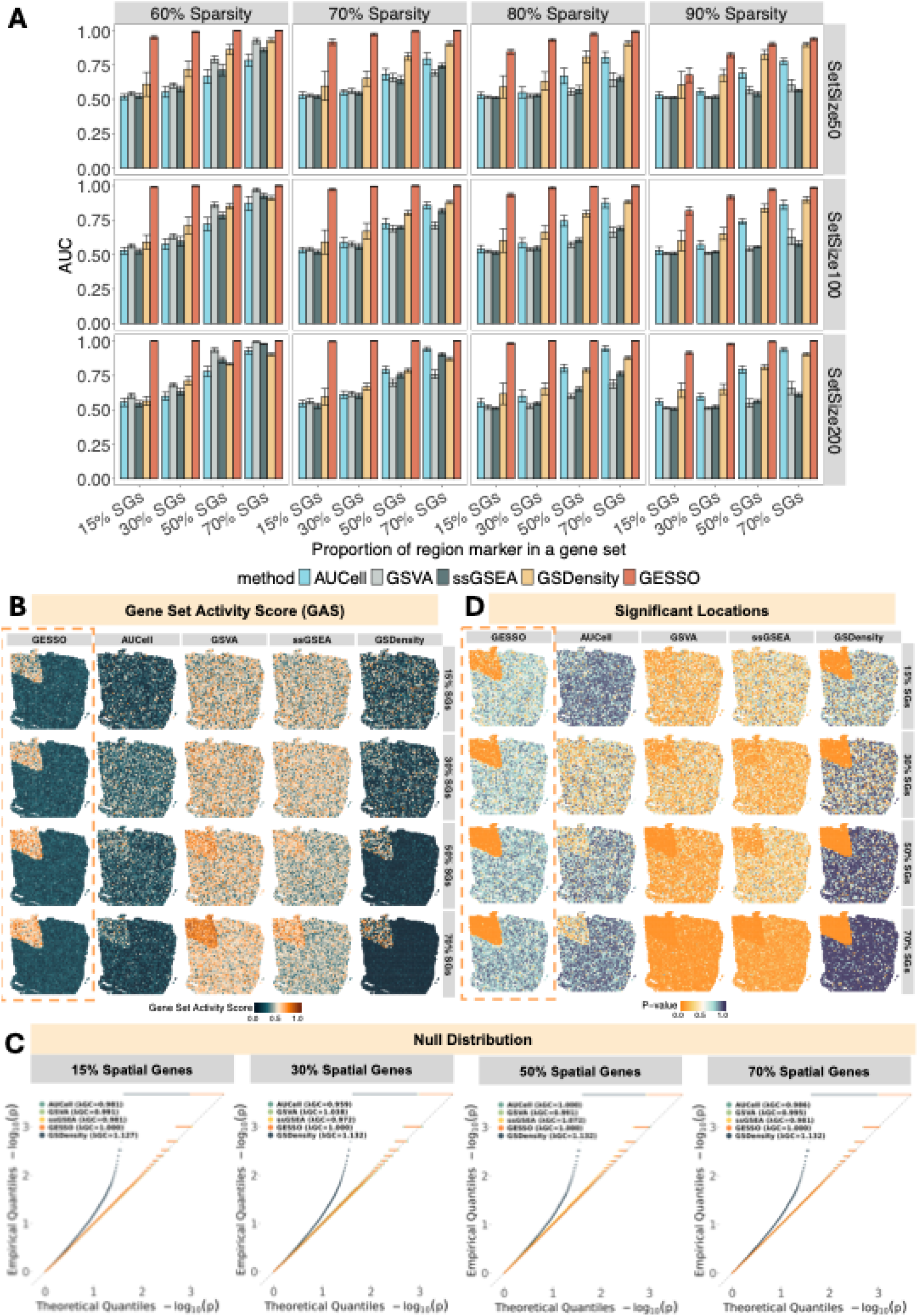
Simulation study results. **A.** Bar plots of area under the receiver operating characteristic curve (AUC) measuring performance of classifying the white matter (WM) region by gene set activity scores (GAS) derived from different methods. Each AUC was obtained from a logistic regression of the binary WM label on GAS for a single gene set. Error bars show the standard error across replicated gene sets. **B**. Spatial visualization of GAS computed by each method. The maps display scores for a representative gene set of 50 genes at 60% data sparsity, across four different proportions of spatial genes. All GASs are processed by min-max scaling for visualization. GESSO results are highlighted by orange. **C**. Quantile-quantile of −log10(*p*-values) from all methods measured by null gene sets of 50 genes at 60% sparsity level. *λ*_*GC*_ is the genomic control factor which closes to 1 indicating a better false positive control. **D**. Spatial maps of locations identified as having statistically significant pathway activity (p < 0.05) by the permutation-based hypothesis test, for the representative gene set of 50 genes at 60% data sparsity. GESSO results are highlighted by orange.

Next, to assess whether high-scoring regions corresponded to the true WM region, we visualized the spatial distribution of the GASs across all methods. We first examined a representative gene set containing 50 genes under different scenarios with varying proportions of SGs at 60% sparsity level (comparable to the sparsity level of the widely applied 10x Visium platform). GESSO produced GAS distributions that were strongly enriched in the WM region, whereas competing methods yielded weaker or more diffuse patterns (Figure 2B). Similar patterns were observed for gene sets with larger sizes (100 and 200 genes, Supplementary Figure S2). When evaluated under increased sparsity (70%, 80% and 90%), GESSO continued to yield spatially coherent activity maps aligned with WM regions, while competing methods produced noisier or less localized patterns (Supplementary Figure S2).

To evaluate whether an estimated GAS is significantly elevated relative to a background distribution, we conducted a permutation-based hypothesis testing procedure that computes a p-value for each spatial location for each gene set (Methods and Supplementary Note 1). We focused on two scenarios with representative sparsity levels (60% and 90%) reflecting empirical characteristics of current SRT platforms and applied the testing procedure to all gene set sizes. We first examined null distributions using control gene sets composed of genes with random expression patterns (RGs). At the global level, null p-values across all locations are expected to follow a Uniform (0,1) distribution. Consistent with this expectation, all methods except GSDensity exhibited well-calibrated null distributions (Figure 2C and Supplementary Figure S3). GSDensity displayed substantial p-value inflation, with genomic inflation factors (*λ*_GC_) much greater than 1, indicating poor false-positive control compared to the other methods (Methods, Figure 2C and Supplementary Figure S3). To further assess calibration at the local level, we computed the Kolmogorov-Smirnov (K-S) statistic at each spatial location to quantify deviations of spot-specific null p-values from the expected uniform distribution (Methods). Mapping these statistics across tissue space provided a spatial diagnostic of calibration quality: uniformly low values indicate robust local control, whereas spatially structured elevations suggest false-positive inflation. GESSO, ssGSEA and GSVA produced uniform distributions (Supplementary Figure S4-S6), while AUCell and GSDensity exhibited the WM region with higher K-S statistics than the remaining area, indicating a local inflation of p-values under the null (Supplementary Figure S7-S8). Together, these analyses demonstrate that GESSO controls false positives effectively, both globally and locally across tissue space. After confirming calibration under null conditions, we applied the testing procedure to representative simulated gene sets for each gene set size (Methods). Specifically, at 60% sparsity level with a gene set size of 50, GESSO identified significant spatial locations (p < 0.05) concentrated in the expected WM regions (Figure 2D and Supplementary Figure S9). Among other methods, only GSDensity partially localized WM regions across scenarios with varying percentage of SGs, while the other methods fail to recover the expected pattern. At higher sparsity (90%), reflecting the extreme dropout of newer platforms, GESSO maintained strong performance, whereas the performance of competing methods further declined (Supplementary Figure S10). For larger gene sets (100 and 200 genes), GESSO and GSDensity again produced significant signals aligned with WM, while the other methods did not (Supplementary Figures S9-S10).

To quantify the spatial concordance between significant GAS locations and the WM region, we used two complementary metrics. First, we computed the AUC treating p-values as a continuous decision statistic for threshold-based classification (ROC analysis) for predicting the WM label (Methods). Second, we calculated the Dice coefficient^39^ (Methods) to measure the overlap between significant spots (p < 0.05) and the WM mask, with higher values indicating stronger concordance. Across all scenarios, GESSO consistently ranked among the top two AUC and Dice coefficients, while AUCell, GSVA, and ssGSEA performed substantially worse (Supplementary Figure S11-16). GSDensity produced comparable concordance to GESSO under the alternative, but as shown in the null experiments (Figure 2C, Supplementary Figures S3 and S8), its p-values were systematically inflated, limiting the reliability of its significance calls. As an illustration, at 60% sparsity with gene set size equals to 50 (Supplementary Figures S11), GESSO reached near-perfect performance (AUC = 0.97-1; Dice = 0.78-0.99) across varying proportions of SG marker gene settings, while AUCell, GSVA, and ssGSEA were markedly lower (AUC = 0.59-0.93; Dice = 0-0.49). GSDensity achieved similar AUC and Dice values under this setting, but its lack of calibration under null scenarios makes these results less trustworthy. Similar trends were observed for larger gene sets and at higher sparsity (Supplementary Figures S12-S16).

### GESSO delineates tumor and immune functional regions across six 10x Visium human cancer datasets

We next evaluated GESSO and the other methods on six 10x Visium human tumor SRT datasets including breast ductal carcinoma (BRCA; 2,518 spatial locations), colorectal cancer (CRC; 2,660 spatial locations), lung squamous cell carcinoma (LUSC; 3,858 spatial locations), ovarian carcinoma (OVCA; 3,455 spatial locations), cervical cancer (CESC; 2,781 spatial locations), and prostate adenocarcinoma (PRAD; 4,371 spatial locations). For each dataset, we calculated GASs for a total of 8,729 qualified pathways (Methods and Supplementary Note 4) curated from the MsigDB database^6,40^ using GESSO and four other methods (AUCell, ssGSEA, GSVA and GSDensity).

First, we examined the BRCA dataset, which includes expert-annotated tissue regions provided by the 10x Genomics Platform (Supplementary Figure S17). Prior research has shown that spatially distinct tissue regions often exhibit divergent pathway activity patterns^41–45^. An effective pathway activity scoring method should therefore generate GASs that accurately characterize diverse functional regions in SRT data. To evaluate biological relevance, we performed a region classification using the 8,729 GASs as predictors of the annotated tissue regions (Methods). Specifically, we trained logistic regression models on GASs from each method to predict region labels (Figure 3A). The predictive accuracy of these models served as a measure of GASs quality, under the rationale that biologically informative GASs should correspond to higher classification performance. Among all methods, GESSO achieved the highest AUC (0.88), outperforming AUCell (0.80), GSVA (0.82), ssGSEA (0.81), and GSDensity (0.81) (Figure 3B). Accuracy and F1-score showed the same trend (Supplementary Figure S18), confirming that GESSO produces more predictive and biologically meaningful measurement of pathway activities. Moreover, a sensitivity analysis of the spatial smoothing strength parameter further confirmed that incorporating spatial information enhances prediction performance (Methods).

**Figure 3.**
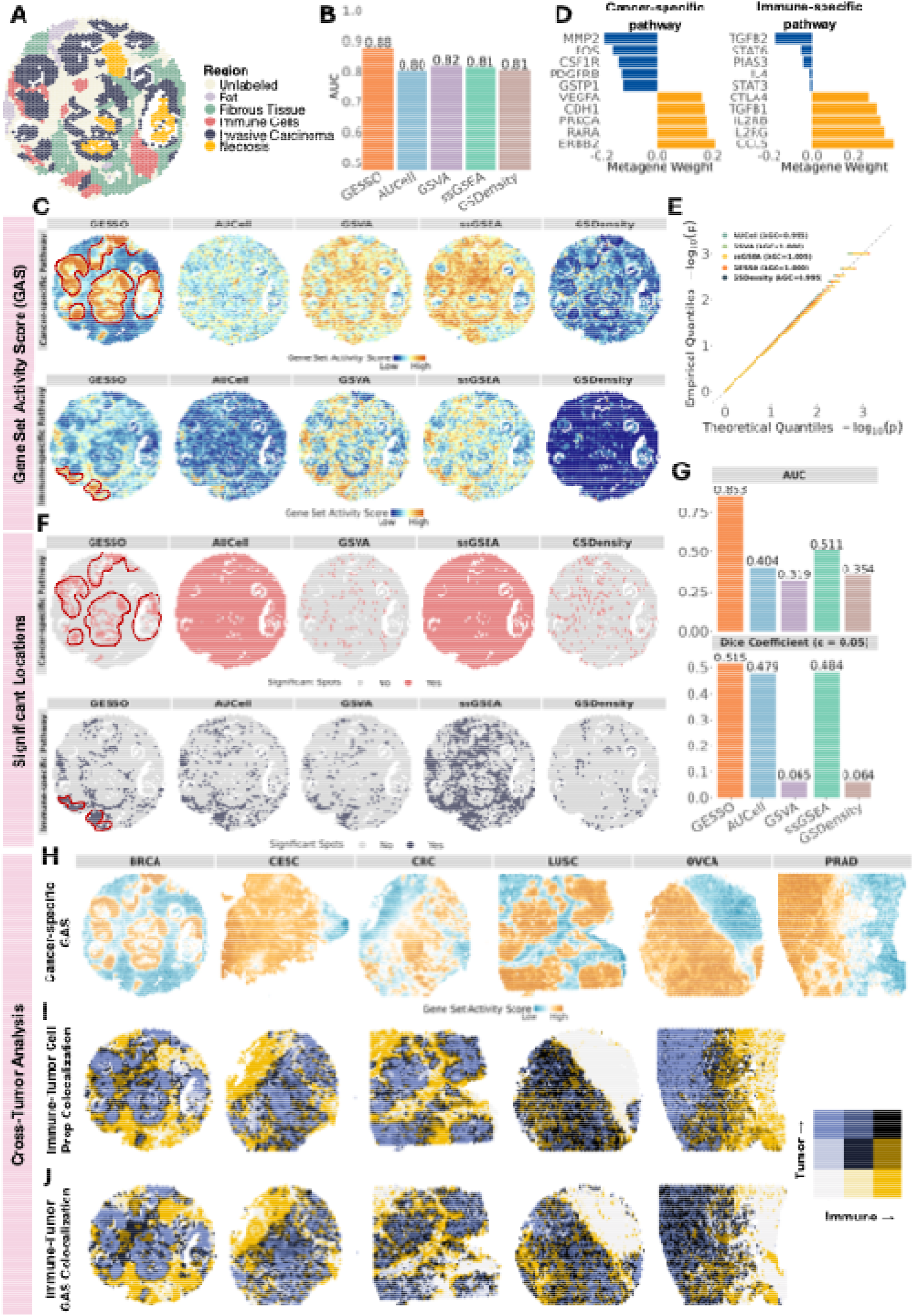
10x Visium human tumor datasets analysis results. **A.** Spatial map of annotated spatial locations for the BRCA dataset. **B**. Bar plots displaying AUC of logistic regression models trained to predict spatial location annotations with GASs generated by GESSO, AUCell, GSVA, ssGSEA, and GSDensity as predictors in BRCA. **C**. Spatial maps of GASs computed by GESSO, AUCell, GSVA, ssGSEA, and GSDensity for a cancer-specific pathway and an immune-specific pathway in BRCA. The regions with high GESSO’s GASs are highlighted. All GASs are processed by min-max scaling for visualization. **D**. Bar plots displaying the top 5 positive and negative metagene weights from GESSO’s cancer-specific and immune-specific GASs. **E**. Q-Q plot comparing p-values computed by all methods for 1,000 randomly generated pathways against the uniform distribution in BRCA. *λ*_*GC*_ is the genomic control factor which closes to 1 indicating a better false positive control **F**. Spatial maps of locations exhibiting significant cancer-specific and immune-specific GAS (p-value < 0.05) based on the local activity enrichment hypothesis testing in BRCA. GASs used in the hypothesis testing framework were computed by GESSO, AUCell, GSVA, ssGSEA, and GSDensity. **G**. Bar plots displaying spatial concordance (measured by AUC and Dice Coefficient) of significant GASs generated by all methods from cancer-specific pathway to the Invasive Carcinoma region. **H**. Spatial maps of GASs generated by GESSO for the cancer-specific pathway across six 10x Visium datasets. **I**. Bivariate spatial maps of immune and tumor cell type proportions driven from cell type deconvolution by CARD. **J**. Bivariate spatial maps of immune-specific and cancer-specific GESSO GASs. Darker color indicates more colocalization patterns.

We next examined whether GESSO accurately detected spatially localized tumor- and immune-specific pathway activities. Mapping pathway activity within tissue context provides insight into how oncogenic signaling and immune responses are organized within the tumor microenvironment^20^. Specifically, we focused on two representative pathways with well-characterized roles in tumor progression and immune regulation: a cancer-specific pathway *KEGG_PATHWAYS_IN_CANCER*, capturing core oncogenic processes, and an immune-specific pathway *WP_INTERACTIONS_BETWEEN_IMMUNE_CELLS_AND_MICRORNAS_IN_TUMOR _MICROENVIRONMENT*, reflecting key immune interactions within tumor microenvironment. Spatial maps revealed that GESSO produced GASs closely aligned with annotated tumor and immune regions (Figure 3A and 3C). For the cancer-specific pathway, GESSO scores peaked in the Invasive Carcinoma region (Figure 3C, the peak region is highlighted), whereas AUCell, GSVA, and GSDensity failed to recover this carcinoma-specific activity, and ssGSEA generated diffuse signals extending into non-carcinoma regions (Figure 3A and 3C). For the immune pathway, GESSO accurately localized immune-cell-enriched regions (Figure 3C, the peak region is highlighted), while GSVA and ssGSEA showed partial concordance but with spillover into adjacent compartments, and AUCell and GSDensity failed to capture immune localization (Figure 3A and 3C). These results underscore that GESSO yields spatially coherent and biologically interpretable pathway maps.

Unlike existing methods that provide only GASs, GESSO also provides interpretable metagenes quantifying gene-specific contributions to each pathway. This allows us to directly examine which genes drive the gene set activity (Methods). For the cancer-specific pathway, we found that *ERBB2, RARA*, and *PRKCA* were top three weighted genes, all well-known oncogenic drivers whose increased expression promotes tumor progression^46–53^ (Figure 3D). The dominance of these genes explains the high cancer-specific pathway activity detected in invasive carcinoma regions. Similarly, for the immune-specific pathway, the top three weighted genes were *CCL5, IL2RG*, and *IL2RB*, all known mediators of immune signaling within the tumor microenvironment^54–57^(Figure 3D). Their elevated expression in immune-enriched regions explains the strong spatial concordance between high immune pathway activity region and the annotated immune cell region. Importantly, GESSO also assigned negative weights to immunosuppression genes (e.g., *TGFB2*^58^,*STAT3*^59^; Figure 3D), suggesting that high pathway activity in immune-rich regions is characterized by a dual signature: the upregulation of immune-activating genes alongside the downregulation of key immunosuppressive genes.

To assess whether GESSO-derived GASs reflect significant activity levels beyond random background variation, we applied permutation-based hypothesis testing that we performed in simulations (Methods). We first assessed how the testing procedure controls for false positives under the null for all the methods. Using control gene sets, GESSO produced well-calibrated null distributions globally (Figure 3E; Supplementary Figure S19), whereas GSDensity showed marked inflation, particularly for the immune-specific pathway (Supplementary Figure S19). Other methods exhibited moderate but acceptable calibration under the null. To further examine calibration locally, we computed K-S statistics at each spatial location, comparing null p-value distributions from control sets against the expected uniform (Methods). GESSO, ssGSEA, and GSVA displayed uniformly low K-S statistics across all spatial locations, indicating reliable false-positive controls (Supplementary Figure S20). In contrast, both AUCell and GSDensity showed relatively higher K-S statistics in some regions, suggesting spatially localized inflation. We then perform the testing procedure to the cancer- and immune-specific pathways GAS and assessed their p-values. GESSO identified significant regions (p < 0.05) concentrated within annotated tumor and immune regions, whereas other methods showed weaker or more diffuse signals (Figure 3F, significant locations are highlighted). Using p-values as continuous decision statistics for predicting either cancer or immune annotated regions, GESSO achieved the highest AUC for both pathways (cancer: 0.85, immune: 0.77), outperforming AUCell (0.40, 0.66), GSVA (0.32, 0.74), ssGSEA (0.51, 0.75), and GSDensity (0.35, 0.63) (Figure 3G, Supplementary Figure S21). In addition to AUC, GESSO also achieved the highest Dice coefficients (cancer: 0.52, immune: 0.31), reflecting superior spatial concordance between significant locations and annotated regions. Other methods showed substantially lower values, including AUCell (0.48, 0.26), GSVA (0.07, 0.27), ssGSEA (0.48, 0.26), and GSDensity (0.06, 0.11) (Figure 3G, Supplementary Figure S21). These results demonstrate that GESSO reliably localizes biologically meaningful pathway activity across diverse tumor regions while maintaining robust global and local false-positive control.

Finally, we extended GESSO to perform a comprehensive pan-cancer analysis across six tumor types: BRCA, CRC, LUSC, OVCA, CESC, and PRAD. GESSO-derived cancer-specific pathway maps revealed spatially distinct regions with significantly increased oncogenic signaling (Figure 3H). We compared these GESSO-derived GASs with tumor and immune cell type proportions estimated by CARD^22^ (Methods). The CESC dataset was excluded from this analysis due to the lack of a matched single-cell RNA-seq reference for deconvolution. Across remaining datasets, spatial maps of cancer-specific pathway GAS showed strong concordance with the spatial distribution of CARD-inferred tumor cell populations (Figure 3I, blue regions). Moreover, GESSO identified regions with colocalized immune and cancer pathway activities (Figure 3J, black regions) corresponding to mixed tumor-immune niches (Figure 3I, black regions). These findings demonstrate that GESSO captures both the spatial heterogeneity of pathway activities and their coordinated tumor-immune interactions across diverse tumor microenvironments^20,60^.

### GESSO captures organ-specific developmental dynamics from Stereo-seq mouse embryo data

We next applied GESSO to the high-resolution Stereo-seq mouse embryo dataset, which profiles mid-gestation embryo stages (E11.5 with 30,124 spatial locations; E12.5 with 51,365 spatial locations; and E13.5 with 77,369 spatial locations) during active tissue differentiation and growth^61^. GASs were computed for curated gene sets relevant to organ-specific region annotations provided by the original study^61^ (Methods and Supplementary Note 4). All methods were compared except GSVA, which failed to complete on the large dataset (i.e., results for the E12.5 sample could not be generated within two days). To assess the biological relevance, we evaluated how well GASs predicted the annotated organ regions^61^, following the same classification framework in 10x Visium BRCA data (Figure 4A). Logistic regression models trained on GESSO-derived GASs achieved the highest predictive accuracy across all stages (E11.5: 0.86, E12.5: 0.87, E13.5: 0.86), outperforming AUCell (0.81, 0.83, 0.77), ssGSEA (0.78, 0.79, 0.71), and GSDensity (0.33, 0.23, 0.17) (Figure 4B). AUC and F1 scores showed consistent trends (Supplementary Figure S22), indicating that GESSO most effectively delineates region-specific pathway activities.

**Figure 4.**
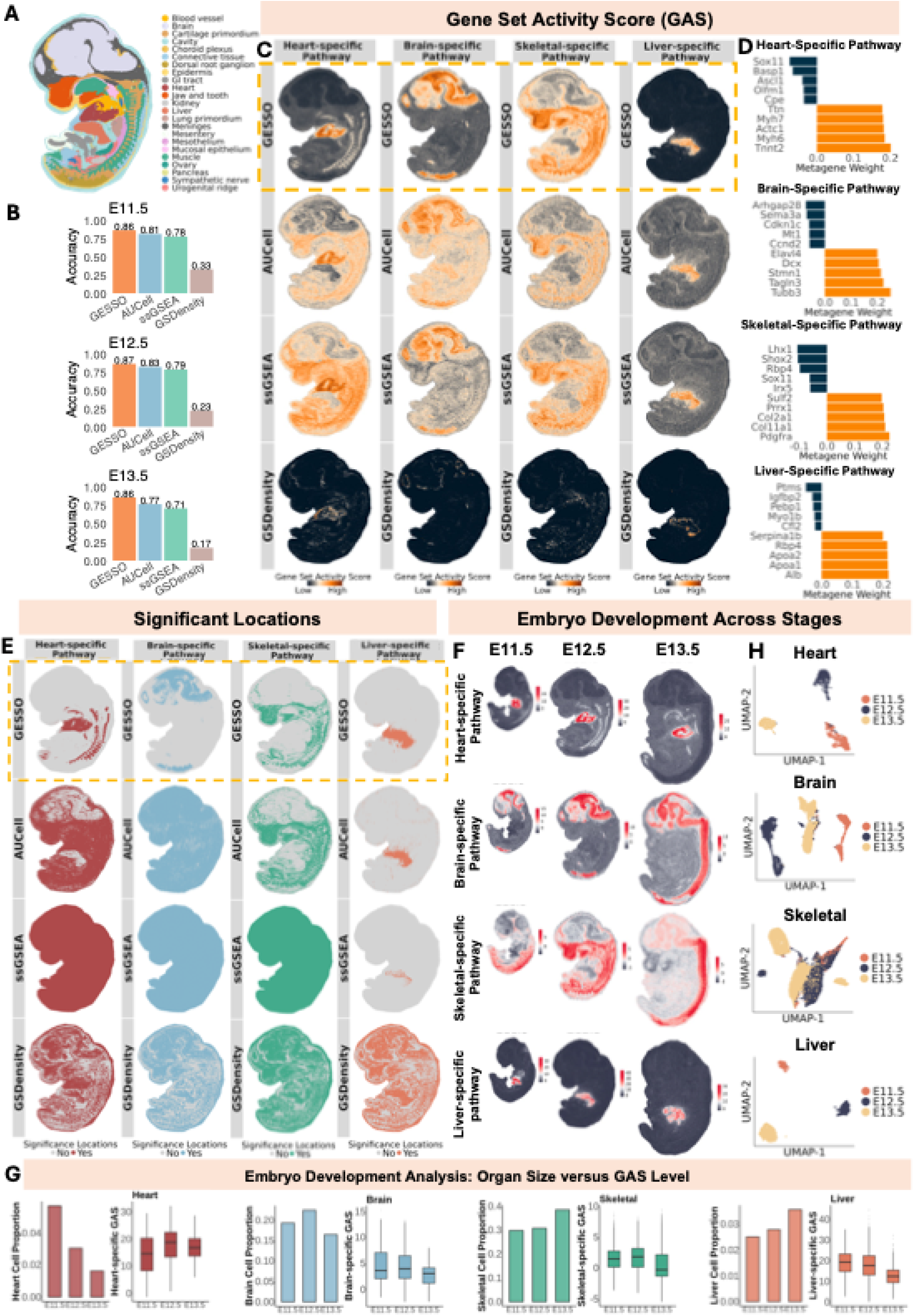
Stereo-seq mouse embryo dataset analysis results. **A.** Spatial map of cell type annotations for the Stereo-seq E12.5 mouse embryo dataset. **B**. Bar plots displaying accuracy of logistic regression models trained to predict cell type annotations with GASs generated by GESSO, AUCell, ssGSEA, and GSDensity as predictors across three stages (E11.5, E12.5, E13.5). **C**. Spatial maps of the heart-, brain-, skeletal-, and liver-specific GASs as computed by GESSO, AUCell, ssGSEA, and GSDensity. All GASs are processed by min-max scaling for visualization. **D**. Bar plots displaying the top 5 positive and negative metagene weights from GESSO’s region-specific GASs. **E**. Spatial maps of significant locations (p-value < 0.05) generated via the proposed local activity enrichment hypothesis testing method for identifying spatial locations with significant elevated pathway activity. **F**. Spatial maps of GASs generated by GESSO for the heart-, brain-, skeletal-, and liver-specific pathways across the three development stages. **G**. For each of four organs/systems (heart, brain, skeletal, liver): proportion of cells labeled as relevant to the organ/system (left column) and box plots of organ/system GAS distributions across development stages (right column). **H**. Two-dimension embedding of the GASs of all organ/system-specific pathways stratified by development stage.

We next examined how well each method could recover anatomically coherent patterns of organ-specific activity. Focusing on the E12.5 embryo stage, where overall prediction accuracy was highest across methods, we focused on four major systems undergoing organogenesis—the brain, heart, liver, and skeletal system—to investigate region-specific pathway activity. Corresponding representative organ pathways include *MANNO_MIDBRAIN_NEUROTYPES_HNPROG* (brain), *GOBP_HEART_DEVELOPMENT* (heart), *GOBP_EMBRYONIC_SKELETAL_SYSTEM_DEVELOPMENT* (skeletal), and *AIZARANI_LIVER_C11_HEPATOCYTES_1* (liver) (Methods). Across all four pathways, GESSO produced spatially distributed GASs that closely matched their expected anatomical regions, yielding sharper and more coherent patterns than the other methods (Figure 4A and 4C). Examination of metagenes revealed top-weighted genes corresponded to known markers of each organ (Figure 4D). For example, the heart-specific pathway’s top three genes were *Tnnt2, Myh6*, and *Actc1*, all key regulators of cardiac development and contraction^62,63^. The top three weighted genes in the brain-specific pathway were *Tubb3, Tagln3*, and *Stmn1*, aligning with their known enrichment in regions undergoing neurogenesis^64–66^. The skeletal-specific pathway highlighted *Pdgfra, Col2a1*, and *Col11a1*, all crucial for proper skeletal system development^67–69^. Finally, in the liver pathway, *Alb, Apoa2* and *Apoa1* were top-weighted, consistent with their known roles in hepatocyte differentiation and organogenesis^70–72^. We next assessed statistical significance for these organ pathways using the same permutation-based testing framework. Null calibration using control gene sets showed that GESSO consistently produced well-calibrated null p-value distributions: QQ plots evidenced global false-positive control (Supplementary Figure S23), while K-S statistics confirmed local false-positive control (Supplementary Figure S24). In contrast, AUCell was comparably stable, while ssGSEA and GSDensity exhibited deflation and inflation, respectively. When applied to real pathways, GESSO identified significant activity localized to the annotated organ regions (Figure 4E), achieving the highest AUCs for predicting organ labels (Supplementary Figure S25, e.g., heart-specific pathway: GESSO 0.97 vs. AUCell 0.79, ssGSEA 0.50, GSDensity 0.78), with Dice coefficients showing similar trends (e.g. GESSO 0.400 vs. AUCell 0.076, ssGSEA 0.059, GSDensity 0.093).

We further extended the analysis across three developmental stages, E11.5, E12.5, and E13.5 to characterize temporal dynamics of organ-specific pathway activity. Spatial maps of the brain-, heart-, skeletal-, and liver-specific pathways showed that GESSO-derived activity consistently aligned with annotated organ regions across all stages (Figure 4F). To assess how pathway activity related to overall organ growth, we compared GESSO-derived activity scores (boxplots) with the relative proportion of each annotated organ (bar plots) across stages (Figure 4G). In all systems, the stagewise ordering of pathway activity differed from that of organ proportion, indicating that changes in anatomical size did not correspond to shifts in pathway activity. For example, the heart region decreased in relative size but showed slightly elevated pathway activity. Brain activity declined slightly while brain proportion first increased and then decreased. Skeletal and liver regions expanded in size, yet both exhibited decreasing pathway activity across stages. These results suggest that GESSO captures molecular developmental dynamics that are distinct from tissue growth. UMAP embeddings^73^ further revealed system-specific temporal dynamics (Figure 4H). Heart-related pathways showed a clear stage-wise progression from E11.5 to E13.5, whereas brain-related pathways displayed a non-linear pattern with E13.5 positioned between E11.5 and E12.5. Skeletal pathways showed substantial overlap across stages, while liver pathways formed three distinct stage-specific clusters rather than a continuous trajectory. Together, these results show that GESSO resolves developmental dynamics beyond morphological growth, revealing both linear and non-linear trajectories of organ maturation.

### GESSO reveals cell-type-specific pathway activity and pathway colocalization in Visium HD human colorectal cancer data

Next, we analyzed paired Visium HD datasets with subcellular resolution derived from colorectal tumor tissue (541,968 spatial locations) and adjacent normal tissue (435,773 spatial locations) of the same patient. Our goal was to evaluate whether spatial variation in GASs corresponded to cell type organization within the tumor microenvironment. Cell type annotations were available only for the tumor tissue in the original study^74^. We focused our analysis on pathways that were significantly enriched in the annotated cell types (Methods and Supplementary Note 4). At this resolution, GSDensity and GSVA failed to converge within 2 days at maximum computational capacity, so comparisons were limited to GESSO, AUCell, and ssGSEA. GESSO achieved the best overall performance, yielding the highest accuracy (0.90 for GESSO, vs. 0.86 for AUCell and 0.88 for ssGSEA), AUC (0.98 vs. 0.97 and 0.98), and F1 score (0.83 vs. 0.78 and 0.82) for predicting annotated cell types in the tumor tissue (Supplementary Figure S26). These results indicate that GESSO delineates cellular organization more accurately than other methods in high-resolution SRT data.

We then examined pathway activities associated with specific cell types. At the intestinal epithelial compartment along the tumor-normal interface (Figure 5A), GESSO identified strong activity for the cell membrane-associated pathway, *GOCC_INTRINSIC_COMPONENT_OF_PLASMA_MEMBRANE*, with high scores precisely localized to the intestinal epithelial region, while the other methods failed to recover this pattern (Figure 5B). This enrichment is consistent with reports linking membrane-associated programs to epithelial differentiation and tumor-cell infiltration in colorectal cancer^75,76^. Within this pathway, the intestinal epithelial marker *PIGR* ranked among top weighted genes (Figure 5C), underscoring its cell-type specificity. In contrast, immune-related genes *ITGA5* and *TGFBR2* showed the largest negative weights, suggesting reduced immune infiltration^77^ and decreased tumor-suppressive signaling^78^. GESSO’s GASs most accurately delineated the intestinal epithelial boundary from the other regions with the highest AUC (Figure 5D, GESSO = 0.88, ssGSEA = 0.73 and AUCell = 0.70). We next evaluated statistical significance using permutation-based testing. Under maximum computational capacity, only GESSO and AUCell completed null distribution construction. GESSO produced well-calibrated global null p-values, whereas AUCell displayed slight deflation (Supplementary Figure S27). K-S statistics further confirmed robust calibration in local false-positive for GESSO (Supplementary Figure S28). When mapping significant locations, GESSO revealed a strongly spatial localized enrichment within the intestinal epithelial region, while AUCell failed to capture this pattern (Figure 5E, Supplementary Figure S29). Quantitatively, GESSO achieved higher spatial concordance with annotated epithelial regions (AUC = 0.94, Dice = 0.74), outperforming AUCell (AUC = 0.70, Dice = 0.13; Figure 5F).

**Figure 5.**
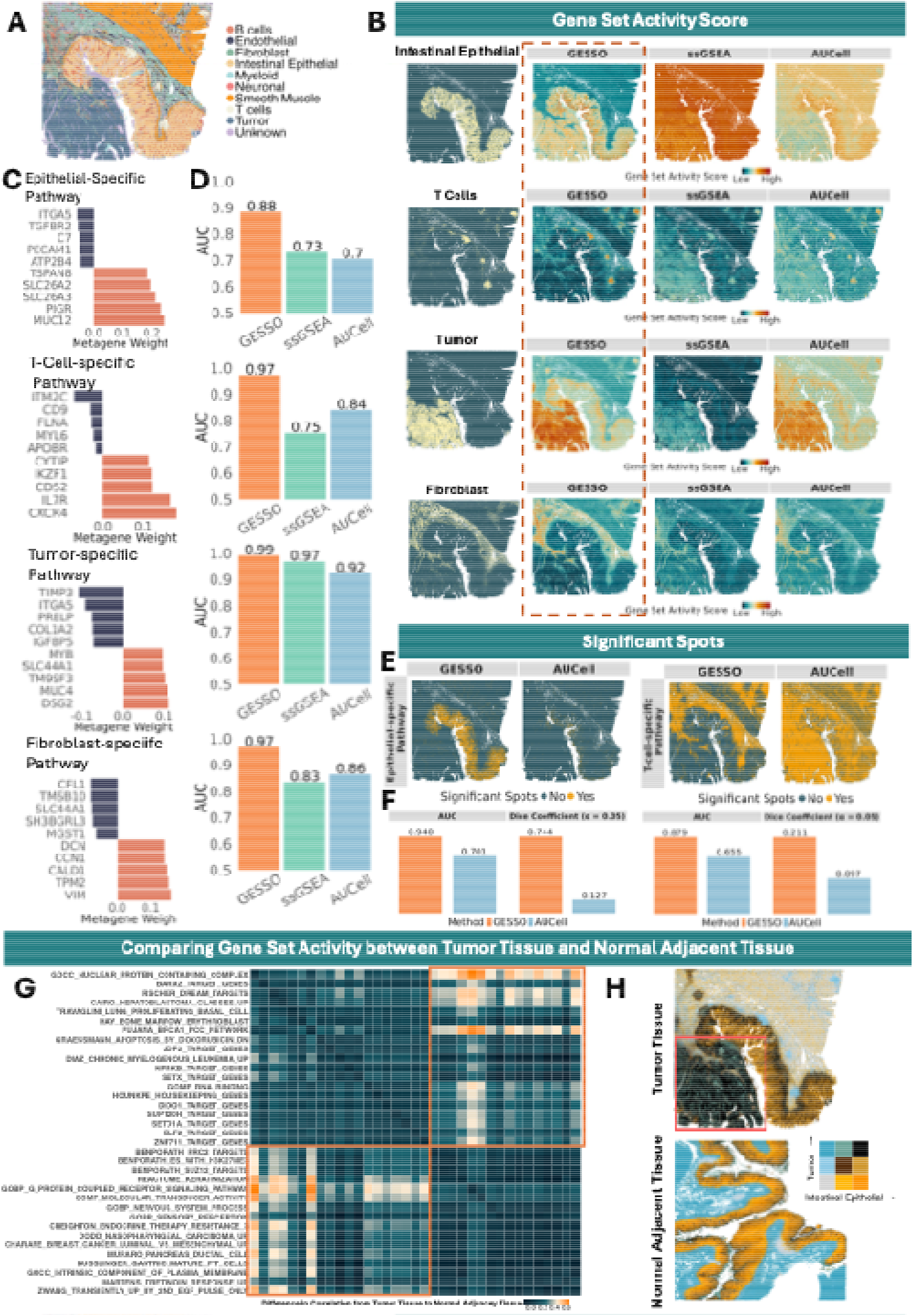
Visium HD human colorectal cancer dataset analysis results. **A.** The spatial map of annotated cell type in Visium HD CRC tumor data. **B**. Spatial map of annotated tissue regions and region-specific GASs measured by GESSO and other methods. The tissue regions, from the top to the bottom, are Intestinal Epithelial, T cells, Tumor and Fibroblast. The GASs, from the left to right, are calculated by GESSO, ssGSEA and AUCell. All GASs are processed by min-max scaling for visualization. GESSO’s results are highlighted by red. **C**. Bar plots displaying the top 5 positive and negative metagene weights from GESSO’s region-specific GASs. **D**. Bar plots displaying AUC evaluated from a univariate logistic regression with region label as a response variable and its region-specific GAS as an independent variable. **E**. Spatial maps of significant locations (p-value < 0.05) generated via the proposed local activity enrichment hypothesis testing method for identifying spatial locations with significant elevated Intestinal-epithelial-specific and T-cell-specific pathway activity. **F**. Bar plots displaying spatial concordance (measured by AUC and Dice Coefficient) of significant GASs generated by GESSO and AUCell from Intestinal-epithelial-specific and T-cell-specific pathway activity to their corresponding annotated region. **G**. Heatmaps presenting a difference of Pearson’s correlation coefficients between GESSO’s GASs from the tumor tissue to the normal adjacent tissue. Lighter orange color indicates a stronger correlation between two pathway activities happening in the tumor tissue compared to the normal adjacency tissue. The orange region highlights the change between a group of tumor-related pathways and a group of epithelial-related pathways. **H**. Bivariate plots showing the co-location patterns between GESSO’s scores of the tumor-specific pathway and the intestinal-epithelial pathway in the tumor sample versus the control sample. Red block highlighted the colocalization of two specific pathways in the tumor region.

Expanding the analysis to additional compartments, GESSO highlighted an immune-specific pathway (*BUSSLINGER_DUODENAL_IMMUNE_CELLS*) and a tumor-specific pathway (*GRAESSMANN_APOPTOSIS_BY_DOXORUBICIN_DN*), whose GAS maps closely aligned with the T-cell and the tumor regions, while other methods failed to do so (Figure 5B). Within the immune-specific pathway, T cell marker genes *CXCR4*^79,80^ and *CD52*^81^ contributed the largest positive weights, whereas tumor suppressor genes *ITM2C*^82^ and *CD9*^83^ exhibited strongest negative weights. In the tumor-specific pathway, genes associated with tumor growth and chemotherapy response (*DSG2, MUC4* and *TM9SF3*)^84^ dominated the metagene weights, while tumor inhibitor genes *ITGA5* and *TIMP3*^85^ showed strongest negative weights (Figure 5C). For both pathways, GESSO again achieved the highest AUCs in predicting their corresponding cell-type regions (Figure 5D, AUCs of GESSO, ssGSEA and AUCell = (0.97, 0.75, 0.84) for T-cell specific pathway, (0.99, 0.97, 0.92) for tumor-specific pathway). Permutation-based testing showed well-calibrated false-positive control globally and locally for GESSO (Supplementary Figures S27 - S28). Significant spatial enrichment was recovered within the expected T-cell compartments, where GESSO outperformed AUCell (AUC = 0.88 vs. 0.66; Dice = 0.21 vs. 0.10; Figure 5F). For the tumor-specific pathway, GESSO achieved strong concordance with annotated tumor regions (AUC = 0.78, Dice = 0.41), comparable to AUCell (AUC = 0.83, Dice = 0.50; Supplementary Figures S29 - S30). GESSO also identified a mesenchymal stromal cell signature (*MURARO_PANCREAS_MESENCHYMAL_STROMAL_CELL*) enriched in fibroblast regions (Figure 5B), which other methods failed to recover. The canonical fibroblast marker gene *VIM*^86^ ranked among the top metagene contributors, consistent with its role in defining fibroblast activity within the mesenchymal stromal-cell program (Figure 5C). Correspondingly, GESSO showed the strongest spatial concordance with annotated fibroblast regions (Figure 5D, AUC = 0.97 vs. 0.83 and 0.86 for AUCell and ssGSEA). With well-controlled false positives both globally and locally (Supplementary Figures S27 - S28), GESSO further recovered significant pathway activity localized to fibroblast-enriched areas (Supplementary Figure S29), achieving higher concordance (AUC = 0.86, Dice = 0.37) than AUCell (AUC = 0.69, Dice = 0.22; Supplementary Figure S30).

Previous studies have shown that pathway activities from distinct biological contexts can colocalize within specific tumor niches^87^. To assess whether such colocalization patterns differ between the tumor and normal adjacent tissue, we computed pairwise Pearson correlations of GESSO-derived GASs separately for each sample. Because cell-type labels were only available for the tumor tissue, pathways were hierarchically clustered based on their correlation structure in the tumor, and this ordering was applied to the normal adjacent tissue for comparison (Supplementary Figure S31). Two major modules were detected: a tumor-associated cluster (Supplementary Figure S31, red block) comprising pathways such as *GRAESSMANN_APOPTOSIS_BY_DOXORUBICIN_DN, PUNJANA_BRCA1_PCC_NETWORK, CARO_HEPATOBLASTOME_CLASSES_UP*, and transcription-factor target sets (*JDP2_TARGET_GENES, ELF2_TARGET_GENES)*; and an epithelial-associated cluster (blue block) containing pathways such as *GOCC_INTRINSIC_COMPONENT_OF_PLASMA_MEMBRANE*. Both modules exhibited stronger intra-cluster correlations in tumor tissue than in normal adjacent tissue, indicating enhanced spatial co-regulation within functional modules in the tumor microenvironment. Correlations between the two modules (Figure 5G, orange block) were also increased in tumor tissue, suggesting emergent cross-talk between tumor- and epithelial-associated programs during tumor progression. To visualize this effect, we generated a bivariate spatial map for two representative pathways, *GRAESSMANN_APOPTOSIS_BY_DOXORUBICIN_DN* and *GOCC_INTRINSIC_COMPONENT_OF_PLASMA_MEMBRANE* (Figure 5H). Their activity levels colocalized (Figure 5H, black region) more strongly within tumor regions compared to the adjacent normal tissue. These findings indicate that tumors exhibit increased spatial colocalization of pathway programs, consistent with coordinated signaling patterns observed in processes such as epithelial-mesenchymal transition^88^.

### GESSO reveals the spatial organization of immune function within germinal centers from Xenium Prime human lymph node data

We next analyzed the Xenium Prime human lymph node dataset (708,983 cells) to quantify pathway activity within germinal centers (GCs) (Methods and Supplementary Note 4). GCs are critical microanatomical sites for B cell maturation, and immune regulation, and are therefore of particular interest in studies of lymphoid tissue biology^89,90^. To investigate the gene set activities defining these structures, we first manually annotated GCs from the hematoxylin and eosin (H&E) tissue image (Methods, Figure 6A, Supplementary Figure S32), which provided reference labels for benchmarking. As in the Visium HD colorectal cancer dataset, only GESSO, AUCell, and ssGSEA successfully computed GASs at this scale, whereas GSDensity and GSVA failed to converge within 2 days. We then benchmarked their performance by predicting GC-labeled cells, following the same evaluation strategy as in previous datasets. GESSO again achieved the strongest performance, yielding the highest AUC (0.91 vs. 0.87 for AUCell and 0.86 for ssGSEA; Figure 6B), as well as the highest accuracy and F1 scores (Supplementary Figure S33).

**Figure 6.**
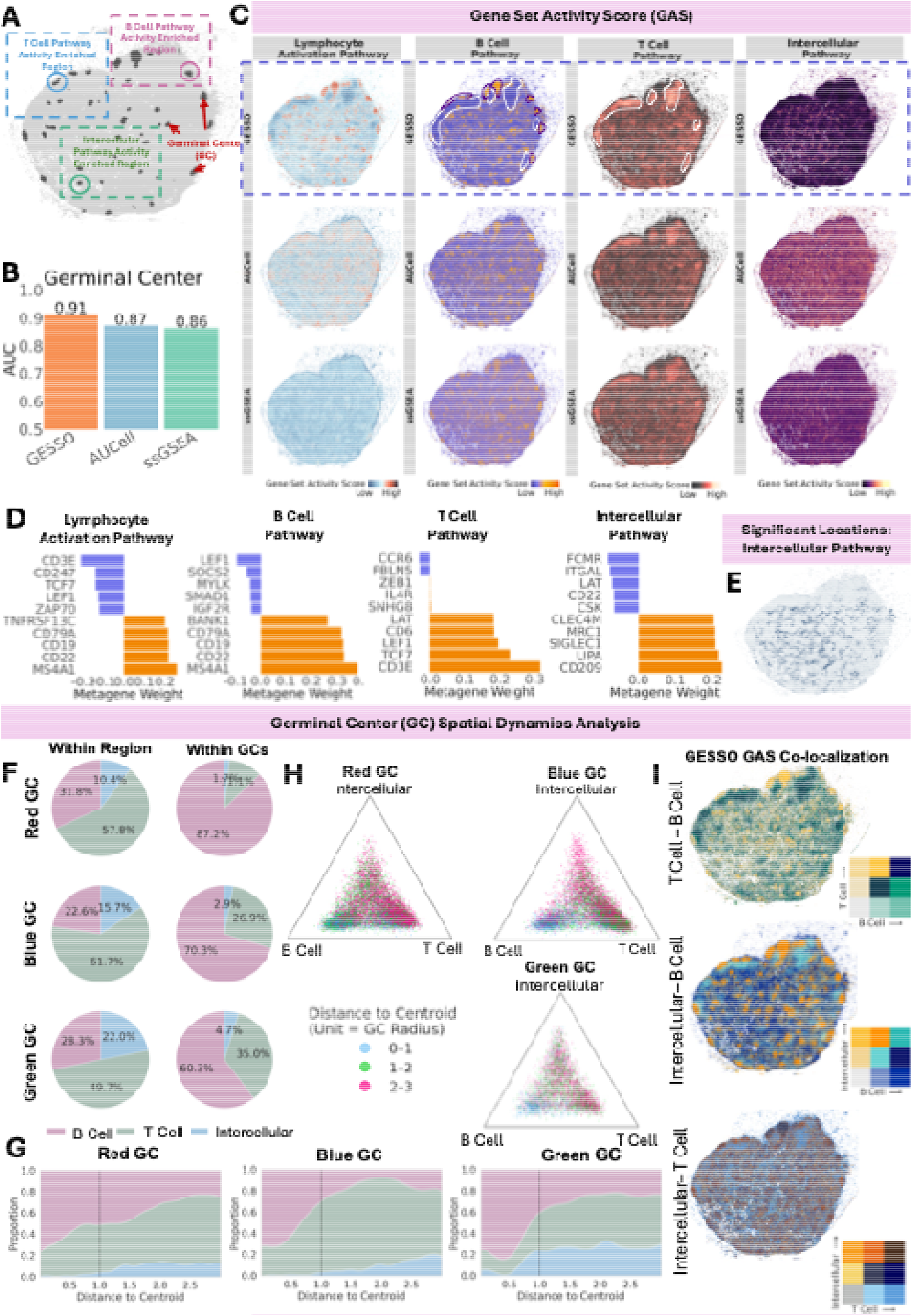
Xenium Prime human lymph node dataset analysis results. **A.** Spatial map of the Xenium Prime human lymph node dataset with GCs (dark gray masks), regions of interest (red, blue, and green boxes), and GCs of interest (red, blue, and green masks). **B**. Bar plots displaying AUC values of logistic regression models trained to predict germinal center cells with GASs generated by GESSO, AUCell, and ssGSEA as predictors. **C**. Spatial maps of the lymphocyte activation, B cell, T cell, and intercellular transport GASs as computed by GESSO, AUCell, and ssGSEA. All GASs are processed by min-max scaling for visualization. In the maps of GESSO, the regions with high B-cell-specific GAS were highlighted by dark purple, and the regions with low B-cell-specific GAS were highlighted by white. **D**. Bar plots displaying the top 5 and the bottom 5 metagene weights from GESSO’s region-specific GASs. **E**. Spatial maps of significant spots (p-value < 0.05) generated via the proposed local activity enrichment hypothesis testing for identifying spatial locations with Intercellular-specific pathway activity. **F**. Proportion of pathways appearing in greatest strength (min-max normalized GAS) across all cells (left column) and GC cells (right column). **G**. Distance-resolved composition plots showing the probability of each pathway—B cell, T cell, or intercellular transport—being dominant as a function of distance from the GC centroid, normalized by GC radius, for the red, blue, and green GCs. **H**. Barycentric plots of GASs for cells in the red, blue, and green GCs. Each point reflects the relative contribution of the three pathways, colored by normalized distance to the GC centroid, illustrating spatial gradients in transcriptional program dominance. **I**. Bivariate plots showing the relationships between germinal activation scores (in terms of GAS) for the three pathways—B cell, T cell, and intercellular transport—displayed for each of the three possible two-pathway combinations.

We then focused on four representative pathways: a lymphocyte-activation pathway (*GOBP_LYMPHOCYTE_ACTIVATION*), a B-cell-specific pathway (*HADDAD_B_LYMPHOCYTE_PROGENITOR*), a T-cell-specific pathway (*HAY_BONE_MARROW_NAIVE_T_CELL*), and an intercellular-transport-specific pathway (*GOBP_VESICLE_MEDIATED_TRANSPORT*). Across all pathways, GESSO produced sharper and more anatomically coherent activity patterns than AUCell and ssGSEA, most notably delineating distinct B and T cell zones and the intervening intercellular interface (Figure 6C). Within GC-labeled regions (Figure 6A), GESSO’s GASs showed strong enrichment for both the lymphocyte-activation and B-cell-specific pathways. These two pathways shared the same top three weighted genes—*MS4A1, CD22*, and *CD19* (Figure 6D)—which are all canonical GC markers essential for B cell receptor signaling, and selection during affinity maturation processes^91–93^. In contrast, the T cell-specific pathway was enriched in regions with lower B-cell pathway activity (Figure 6C, where low and high B-cell-specific GASs are shown in white and dark areas), consistent with a negative correlation between the two pathways (Pearson’s correlation = −0.31). Its top-weighted genes (Figure 6D, *CD3E, TCF7*, and *LEF1*) have been found to mark less-differentiated or memory-prone T-cell populations^94–96^.

This spatial separation delineates the boundary between B cell- and T cell-dominant zones. GESSO revealed that the intercellular transport-specific pathway showed GASs concentrated in the gaps between these compartments (Figure 6C). The top-weighted genes in this pathway were *CD209, LIPA*, and *SIGLEC1*, which mark antigen-presenting cells at the T-B cell interface^97–100^ (Figure 6D). Their spatial co-localization at the interface highlights an active intercellular transport hub coordinating antigen presentation and signaling exchange coordinate B cell activation and T cell help during germinal center reactions. To assess the statistical significance, we applied permutation testing to the lymphocyte-activation, B cell-, T cell-, and intercellular transport-specific pathways. Under maximal computational resources, only GESSO successfully completed the testing procedure. Across all four pathways, GESSO maintained well-calibrated false positive control both globally and locally (Supplementary Figures S34A - S34B). The resulting significance maps for the lymphocyte-activation and B-cell pathways (Supplementary Figure S34C) precisely delineate GC sites (Figure 6A), achieving AUCs of 0.88 and 0.86 for predicting GC labels (Supplementary Figure S34D). For the intercellular transport-specific pathway, GESSO further identified significant GASs at the interface between B and T cell zones (Figure 6E).

Building on GESSO’s ability to reveal distinct spatial patterns of B cell-, T cell-, and intercellular transport-specific pathway activity (Figure 6C), we next examined three lymph-node regions that exhibited varying levels of B cell, T cell, and intercellular transport GASs (Figure 6A, red, blue, and green regions). For each location, we identified the dominant pathway type (B cell, T cell, or intercellular transport-specific pathways) and quantified the proportion of locations dominated by each pathway within each region and within GCs (Methods). This analysis (Figure 6F) confirmed the spatial organization observed in Figure 6C: B cell-specific pathway activity predominated within GCs (87%, 70%, and 60% for the red, blue, and green GCs, respectively), consistent with their role as sites of intense B cell activation, clonal expansion, and affinity-maturation^101–103^. In contrast, T cell-specific activity dominated across the broader lymph node regions (50-62%), reflecting their functions within and surrounding GCs in coordinating immune organization. Intercellular transport-specific activity comprised a smaller fraction (10-22%), but was more prominent in the green region, consistent with its localized enrichment in the spatial maps (Figure 6C). Together, these results show that GESSO effectively delineates spatially organized immune functions within lymphoid tissue.

To further investigate the spatial dynamics of pathway activity within and around GCs, we selected one representative GC from each region (circled in Figure 6A) and quantified the dominant pathway type (B cell, T cell, or intercellular transport-specific pathway) as a function of normalized radial distance from the GC centroid (Methods). This analysis captured gradual spatial transitions in dominant pathway activity from GC cores to peripheral zones (Figure 6G). Across all three GCs, B cell-specific activity peaked at the GC center and declined toward the periphery, typically around one normalized radius. In the blue and green GCs, B cell dominance exhibited a modest rise around 0.5 radii before decreasing, suggesting the presence of a light-zone region enriched for proliferating and selection-phase B cells^101,104^. In the red and blue GCs, this decline was accompanied by gradual increases in T cell and intercellular transport activities, indicating smooth transitions from B cell-rich cores to peripheral immune zones. In contrast, the green GC showed earlier increases in T cell and intercellular transport activity (~0.5 radius), with the latter remaining elevated near the boundary, indicating a sharper peripheral transition. Barycentric coordinate plots further supported these gradients (Methods): cells within one radius of the GC centroid were predominantly B cell-specific GASs, while those beyond two radii exhibited increasing dominance of T cell and intercellular transport GASs (Figure 6H). Finally, spatial bivariate maps of B cell/T cell, B cell/intercellular transport, and T cell/intercellular transport-specific GASs (Figure 6I) visualized these transitions, highlighting localized co-enrichment patterns and transitions in pathway dominance across the GC landscape revealed by GESSO. Collectively, these results demonstrate that GESSO resolves fine-scale pathway organization and radial transitions within germinal centers.

### GESSO resolves distinct cellular niches and EMT programs from CosMx human non-small cell lung cancer data

Finally, we assessed GESSO on CosMx human non-small cell lung cancer data^105^ to identify pathway activities associated with distinct cellular niches that shape the tumor microenvironment (TME) (Methods and Supplementary Note 4). Due to the large number of cells in this dataset (55365 cells), GSVA again failed to converge within 2 days, thus, comparisons were limited to GESSO, AUCell, ssGSEA and GSDensity. Consistent with the results from previous datasets, GESSO achieved the highest predictive performance, with an accuracy of 0.56 (vs. AUCell 0.47 vs. ssGSEA 0.47 vs. GSDensity 0.22), AUC of 0.88 (vs. AUCell 0.82 vs. ssGSEA 0.81 vs. GSDensity 0.6) and F1 score of 0.46 (vs. AUCell 0.35 vs. ssGSEA 0.34 vs. GSDensity 0.03) (Supplementary Figure S35).

Next, we investigated pathways linked to specific TME niches. In both the tumor core and desmoplastic stroma, GESSO more accurately identified niche-specific pathway activities than other methods (Figure 7A). In the tumor core, the GAS for the pathway *TRAVAGLINI_LUNG_ALVEOLAR_EPITHELIAL_TYPE_1_CELL* showed strong enrichment, with top weighted metagenes including *CLDN4*^106^, *EPCAM*^107^ and *KRT19*^108^, all actively involved in tumorigenesis (Figure 7B). By contrast, tumor suppressor genes such as *IGFBP7*^109^ and *TAGLN*^110^ exhibited large negative weights (Figure 7B). In the desmoplastic stroma, characterized by fibroblasts cells^111^, the pathway *TRAVAGLINI_LUNG_ADVENTITIAL_FIBROBLAST_CELL* showed high GASs in this region, and were predominantly influenced by fibroblast markers *COL3A1* and *COL1A*, while the immune response marker *CD74*^112^ showed the largest negative weight (Figure 7B). In contrast, other methods failed to highlight these niche-specific activity in the corresponding areas (Figure 7A). Quantitatively, GESSO achieved the highest AUCs for predicting both niches (AUC = 0.75-0.9, Figure 7C), outperforming ssGSEA (0.61-0.78), AUCell (0.65-0.72) and GSDensity (0.61-0.78). Permutation-based testing showed well-calibrated p-values for GESSO, AUCell, and ssGSEA both globally and locally, whereas GSDensity showed strong inflation (Supplementary Figures S36-S37). Spatially significant GASs from GESSO colocalized most strongly with tumor-core and stromal niches (Supplementary Figure S38), yielding the highest AUC and Dice coefficients compared with AUCell, ssGSEA, and GSDensity (Supplementary Figures S38-S39).

**Figure 7.**
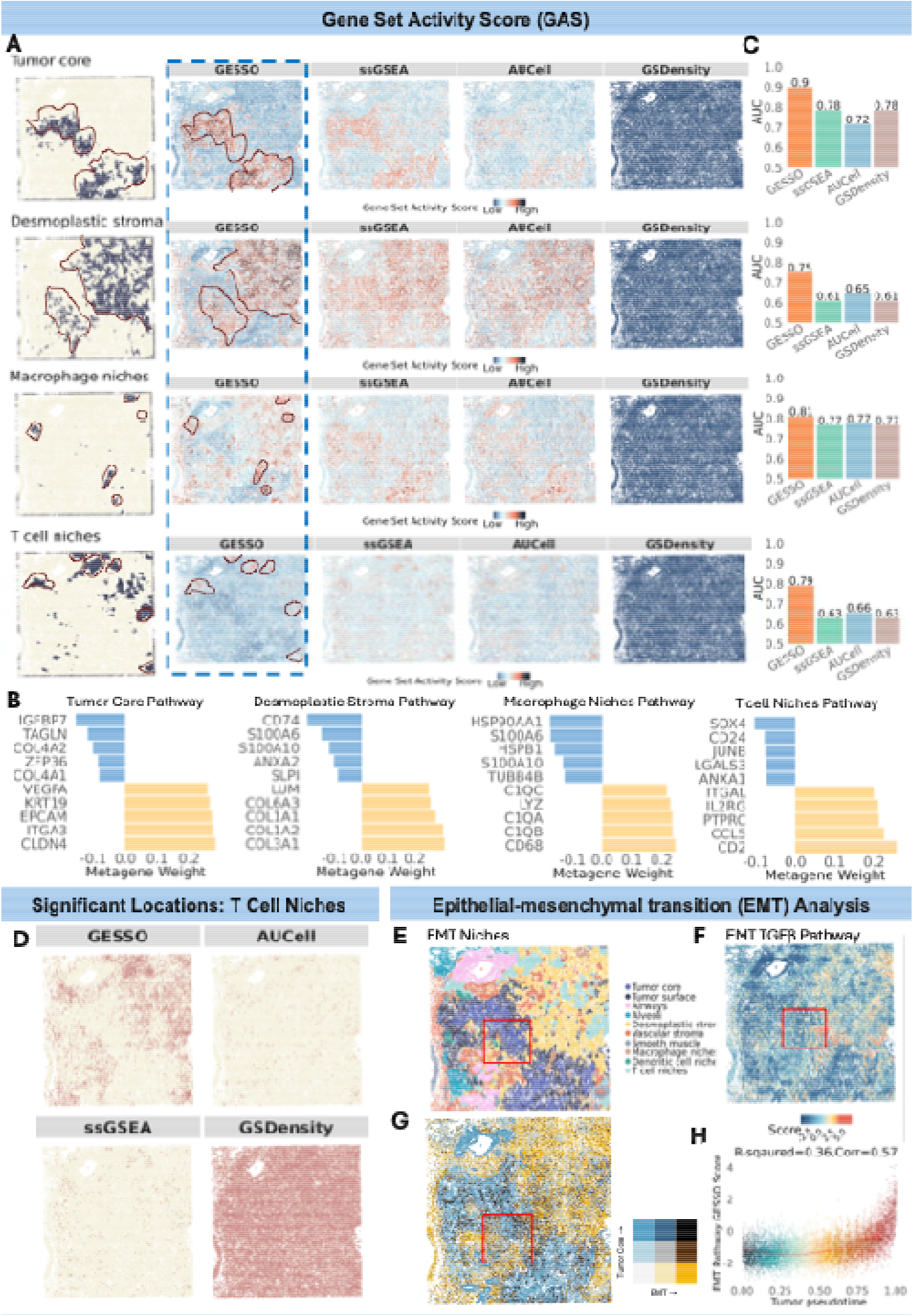
CosMx human non-small cell lung cancer dataset analysis results. **A.** Spatial map of annotated niche regions and niche-specific GASs measured by GESSO and other methods. All GASs are processed by min-max scaling for visualization. The types of niche regions, from the top to the bottom, are Tumor core, Desmoplastic stroma, Macrophage and T cell niches. The GASs, from the left to right, are calculated by GESSO, ssGSEA, AUCell and GSDensity. The locations of niches are highlighted by red. **B**. Bar plots displaying the top 5 positive and negative metagene weights from GESSO’s niche-specific GASs. **C**. Bar plots displaying AUC evaluated from a univariate logistic regression with region label as a response variable and its niche-specific GAS as an independent variable. **D**. Spatial maps of locations exhibiting significant T-cell-niche-specific GAS (p-value < 0.05). **E**. Spatial atlas of identified niches provided by the data. The red block highlighted the EMT niche. **F**. Spatial map of EMT-specific GAS. **G**. Bivariate plots showing the co-location patterns between GESSO’s GASs of the tumor-core-specific pathway and the EMT-specific pathway. **H**. A trajectory plot indicating the relationship between EMT-specific GAS with tumor-pseudotime for tumor cells. *R-squared* is measured by regressing EMT-specific GAS on squared tumor-pseudotime. *Corr* is the Pearson’s correlation between EMT-specific GAS on squared tumor-pseudotime.

Beyond the tumor core and stroma, we also examined pathways associated with immune cell niches. A macrophage-specific pathway (T*RAVAGLINI_LUNG_PROLIFERATING_MACROPHAGE_CELL*) showed high GAS within macrophage niches (Figure 7A), with top metagene weights from macrophage marker genes *CD68*^113^, *C1QB*^114^ and *C1QA*^115^ (Figure 7B). Similarly, a T cell activation pathway (*GOBP_T_CELL_ACTIVATION*) was highly enriched in T cell niches (Figure 7A), driven by *CD2*^116^, *CCL5*^117^ and *IL2RG*^118^, which play key roles in T cell-mediated immune responses (Figure 7B). In both pathways, tumor promoter genes such as *SOX4*^119^ and *HSP90AA1*^120^ showed large negative weights. By contrast, GASs from the other methods produced more diffuse enrichment patterns which failed to locate these small immune niches (Figure 7A). Although both macrophage and T cell niches occupied smaller tissue areas, GESSO more accurately recovered their niche-specific pathway activities, achieving the highest AUC (Figure 7C, for Macrophage niches, GESSO, ssGSEA, AUCell, GSDensity = (0.81, 0.77, 0.77, 0.77); for T Cell niches, (0.79, 0.63, 0.66, 0.63)). Permutation-based tests showed well-calibrated p-values for GESSO, AUCell, and ssGSEA under both the global and local null, whereas GSDensity showed p-value inflation (Supplementary Figures S36-S37). Significant macrophage- and T cell-specific locations identified by GESSO aligned most closely with their respective niches, yielding the highest AUCs (0.72 and 0.78; Figure 7D, Supplementary Figures S38) and Dice coefficients (0.08 and 0.42) among all methods (Supplementary Figures S39).

Epithelial-mesenchymal transition (EMT) is a key process in cancer progression, driving tumor cell migration and invasion^121^. In this CosMx dataset, the original study^105^ identified an EMT niche located between the tumor core and desmoplastic stroma (Figure 7E). To assess EMT activity within the TME, we applied GESSO to an upregulated TGFβ-induced EMT pathway *FOROUTAN_PRODRANK_TGFB_EMT_UP*^122^. GESSO-derived GASs were strongly concentrated in the EMT niche (Figure 7F), with the top metagene weights assigned to the fibroblast cell markers *COL1A1* and *COL5A1*^123,124^, the mesenchymal marker *FN1*^125^, and the metalloproteinase *MMP2*^126^, all central to extracellular matrix remodeling and invasion (Supplementary Figure S40A). Permutation-based testing confirmed well-calibrated p-values for GESSO, AUCell and ssGSEA both globally and locally, whereas GSDensity’s p-values inflation (Supplementary Figure S40B-S40C). The locations with GESSO’s significant EMT-specific GASs aligned most closely with the EMT niche (Supplementary Figure S40D), yielding the highest AUC (Supplementary Figure S39, GESSO, ssGSEA, AUCell, GSDensity = (0.69, 0.64, 0.65, 0.53)) and one of the top Dice coefficients (Supplementary Figure S39, (0.42, 0.44, 0.20, 0.41)). Bivariate spatial maps further showed coordinated enrichment between EMT and both tumor-core (Figure 7G) and desmoplastic-stroma pathways (Supplementary Figure S40E). Finally, we evaluated the relationship between EMT-specific GAS and tumor pseudotime defined by the original study^105^. Specifically, tumor pseudotime reflects the inferred progression of tumor cells from early to late stages. As expected, EMT activity increased along this trajectory: GESSO-derived EMT-specific GASs were positively associated with tumor pseudotime (Figure 7H, R-squared = 0.36, Pearson correlation = 0.57).

## Discussion

We introduced GESSO, an interpretable and scalable method for quantifying gene set activity in SRT data. By leveraging spatially regularized rank-one matrix decomposition with predefined pathways, GESSO produces spatially aware GASs that capture pathway activity across tissue architecture while providing metagene weights that quantify each gene’s contribution. This design enhances interpretability and enables the identification of gene-specific drivers of pathway activity. To provide statistical significance, we developed a local activity enrichment test that evaluates whether observed pathway activity exceeds background variation. Moreover, a stratified low-resolution approximation allows GESSO to scale to datasets containing hundreds of thousands of spatial locations (e.g., Stereo-seq, Visium HD and Xenium Prime) without compromising accuracy. Together, these advances make GESSO a comprehensive tool for sensitive, interpretable, and computationally scalable pathway analysis in SRT data.

Across extensive simulations and 13 datasets spanning multiple SRT platforms and tissue contexts, GESSO consistently outperformed existing gene set activity scoring methods. In simulations, GESSO more accurately recovered region-specific pathway activities and showed superior calibration of both local and global null distributions, yielding statistically robust and spatially coherent significant signals. These advantages carried over to real datasets: in 10x Visium tumors, GESSO more accurately localized cancer- and immune-specific pathways to their associated regions; in high-resolution Visium HD and CoxMx tumors, it resolved complex tumor-microenvironment programs such as epithelial-mesenchymal transition (EMT); in developmental mouse embryo data, it captured temporal trends in growth-related pathway activity; and in human lymph node data, it identified pathway activities reflecting coordinated T-B cell interactions.

Together, these analyses demonstrate that GESSO recovers biologically meaningful pathway patterns that reflect both cellular organization and tissue function.

There are several important future extensions for GESSO. First, although the current method focuses on modeling gene expression, emerging spatial multi-omics technologies suggest opportunities to integrate chromatin accessibility, protein abundance, or perturbation-response layers to better characterize regulatory and therapeutic pathways^127,128^. Second, GESSO is designed for single-slice SRT data but can in principle operate on multiple slices by concatenating data matrices^129,130^. As multi-slice and multi-region spatial atlases become more common^131–133^, extending GESSO to explicitly model inter-slice alignment and shared pathway structure will be increasingly important. Finally, while we used curated gene sets from resources such as MSigDB^6^, the metagene weights inferred by GESSO provide a natural mechanism for refining pathway membership by highlighting context-specific active genes, enabling data-driven pathway adaptation.

## Methods

### GESSO method overview

Here, we present an overview of the GESSO (Gene sEt activity Score analysis with Spatial lOcation) method. Additional technical details, including derivations, are provided in Supplementary Note 1. Briefly, GESSO is a spatially regularized matrix decomposition framework for quantifying GASs in SRT data. Unlike traditional activity scoring approaches that do not account for spatial information explicitly, GESSO integrates gene expression with tissue spatial organization by incorporating a graph Laplacian regularization into the matrix decomposition. This enables biologically coherent inference of spatially varying gene set activity while preserving local spatial structure. Furthermore, GESSO generates metagene weights that quantify the contribution of individual genes to each pathway’s spatial activity, enhancing interpretability and facilitating biological insight into tissue-level functional organization. To ensure statistical rigor, GESSO also supports hypothesis testing to assess the significance of gene set activity scores against randomly distributed null distributions, distinguishing biologically meaningful activity scores from noise. Furthermore, to support large-scale, high-resolution SRT datasets, the algorithm is implemented with a stratified low-resolution approximation framework that substantially reduces computational cost. Together, GESSO provides a principled framework for identifying spatially organized molecular programs and functional heterogeneity within tissues.

Specifically, we denote the SRT gene expression matrix as ***X*** ∈ ℝ ^*G*X*N*^, where *G* is the number of measured genes and *N* is the number of spatial locations. Each column of ***X*** represents the expression profile of a spatial location, and each row corresponds to a gene. The spatial coordinates of these locations are given by ***F*** ∈ ℝ^NX2^, where each row stores the (*x, y*) coordinates of a spatial location. For a predefined gene set *l* consisting of *G*_*l*_ genes, we extract the corresponding submatrix 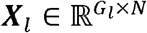. GESSO aims to infer the gene set activity score for the gene set *l* by decomposing ***X***_*l*_ into two components: a metagene vector 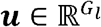, representing the relative contribution of each gene to the overall pathway signal, and a gene set activity score vector ***q*** ∈ ℝ ^N^, which captures the spatial pattern of gene set activity across tissue locations. We formulate the following objective function:

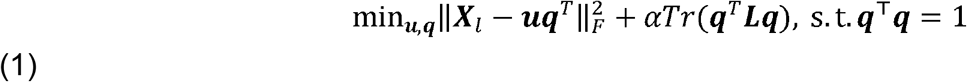

Where || · ||_***F***_ denotes the Frobenius norm, and *Tr* (·) represents the trace operator. ***L*** ∈ ℝ^*N×N*^ is the graph Laplacian matrix, which encodes the spatial relationships among locations. *α* is a regularization parameter that controls the weights of the spatial regularization term. The first term minimizes the reconstruction error, while the second term enforces spatial smoothness between neighboring locations, encouraging spatially coherent pathway activity scores.

To model spatial relationships, GESSO constructs the graph Laplacian matrix ***L*** from the spatial coordinates in ***F***. Specifically, the graph is built using a k-nearest neighbor (k-NN) approach, where each spatial location is connected to its k-nearest neighbors based on Euclidean distance, with k set to 6 for datasets with fewer than 10,000 spatial locations and 20 otherwise (Supplementary Table 4). The resulting adjacency matrix ***A*** has entries *A*_*ij*_ = 1 if locations *i* and *j* are neighbors, and 0 otherwise. The diagonal degree matrix ***D*** is defined as *D*_*ii*_ = Σ_*j*_ *A*_*ij*_ and the graph Laplacian is computed as ***L*** = ***D*** − ***A***, capturing local spatial topology among tissue locations. Minimizing the regularization term *Tr* (***q***^T^ ***Lq***) encourages pathway activity scores to be similar for spatially neighboring locations, reflecting the fact that nearby regions often exhibit similar molecular profiles.

### GESSO optimization procedure

We solve the GESSO’s objective function in (1) to obtain the optimal metagene vector ***u***^*^ and gene set activity score vector ***q***^*^. The details of the derivation of the solution are provided in Supplementary Note 1. Briefly, the optimization first fixes *q* and solves for *u*:

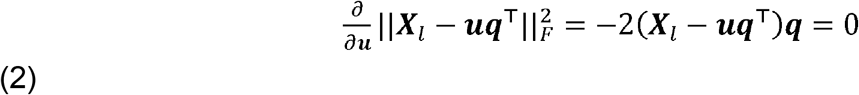

Which yields

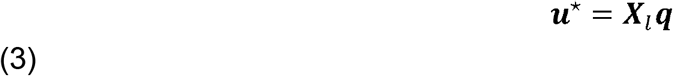

Substituting this expression in (3) into the objective function in (1) yields a reduced optimization problem in ***q***:

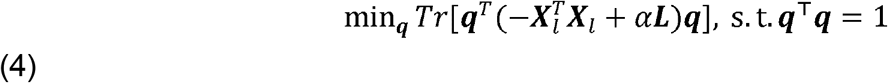

This formulation corresponds to a generalized eigenvalue problem, where ***q***^*^ is the eigenvector associated with the smallest eigenvalue of 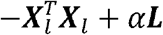 (see Supplementary Note 1).

To make the spatial regularization term more interpretable and to balance the expression and spatial components on comparable scales, we reparametrize the regularization parameter^134^ by defining 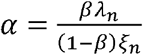, where *λ*_*n*_ denotes the largest eigenvalue of 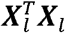 which represents the sample variance of the gene expression data. *ξ*_n_ is the largest eigenvalue of the graph Laplacian ***L***, which represents the magnitude of spatial variation. This formulation provides a normalized trade-off between expression-based and spatial smoothness constraints. Substituting this reparameterization into the objective function in (4) and multiplying by the constant 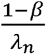 that normalizes the contribution of the two terms while preserving their eigenvectors. This yields:

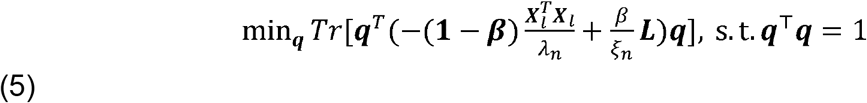

To reformulate this problem in a positive semidefinite form that facilitates interpretation, we add a constant multiple of the identity matrix, (1 − β) ***I***. This addition shifts the eigenvalue scale without changing the eigenvectors, yielding the equivalent normalized formulation:

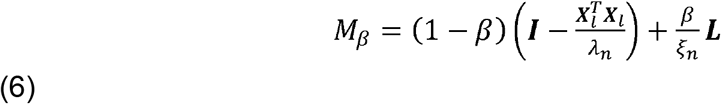

Substituting this matrix form into (5), the problem becomes finding the vector ***q*** minimizes the quadratic form *Tr* (***q***^***T***^ *M*_*β*_ ***q***) Accordingly, the optimal solution ***q*** * is the eigenvector associated with the smallest eigenvalue of *M*_*β*_, and the corresponding metagene is computed as ***u***^*^= ***X***_*l*_ ***q***\*. This transformation makes the regularization term directly interpretable through the trade-off parameter *β*, which is bounded by 0 and 1 and controls the balance between the expression reconstruction and spatial regularization. When *β* = 0 the optimization problem simplifies to standard principal component analysis (PCA), whereas when *β* = 1, the optimization problem becomes that of Laplacian eigenmaps^134^. We set *β* = 0.33 in the process of producing all the results presented in this paper (see Supplementary Table 4).

### Sign correction for interpretability

Matrix decomposition methods, such as singular value decomposition (SVD)^135^ and the graph Laplacian-regularized rank-one approximation described above, inherently suffer from sign ambiguity. That is, in our algorithm, if ***u***^*^ and ***q***^*^ minimize the objective function 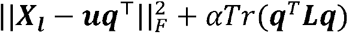, so do − ***u***^*^ and − ***q***^*^. This problem arises because these decomposition methods do not inherently resolve the sign of the resulting components, leading to inconsistencies in GASs in real biological scenarios^136^. To address this challenge, GESSO implemented a sign correction strategy that aligns the direction of the optimized vector ***q***^*^ with a biologically meaningful proxy for pathway activity (Supplementary Note 1). Specifically, we compute the correlation between ***q***^*^ and the overall average expression of genes in the pathway and flip the sign of ***q***^*^ (and ***u***^*^, accordingly) if the correlation is negative. This heuristic ensures sign consistency across runs and aligns GASs with expected biological trends, such as higher activity in regions with elevated overall expression of pathway genes. This novel sign correction approach in GESSO improves the robustness and interpretability of the pathway activity scores. By embedding this strategy into the rank-one approximation framework, GESSO provides more reliable and biologically coherent pathway activity scores, ensuring they align with expected biological trends.

### Scalable gene set activity scoring for large datasets

In contrast to standard matrix decomposition, which allows for reparameterization along either rows or columns, our rank-one approximation is constrained to operate along the rows, corresponding to spatial locations. This constraint is essential for incorporating spatial regularization, which depends on spatial proximity between locations. However, it also introduces substantial memory and computational demands for datasets with tens of thousands of spatial locations (e.g., Visium HD and Xenium Prime). To mitigate this limitation, we implemented a low-resolution approximation by partitioning the dataset into *p* spatially coherent subsets of approximately equal size. To preserve the overall spatial diversity of the tissue within each subset, we use a stratified k-means partitioning strategy. First, spatial locations are clustered using k-means based on their spatial coordinates, setting the number of clusters *k* to the number of desired subsets *p* (see Supplementary Table 4 for the assignment of *p* for each dataset). Then instead of assigning entire clusters to a single subset, we allocate points from each cluster across all subsets, so that every subset contains representative locations from the entire tissue. This ensures that each subset captures the overall spatial heterogeneity of the tissue sample. In practice, we found that stratified partitioning produced more accurate low-resolution approximations of GASs than simple random splitting (Supplementary Note 2 and Supplementary Figure S41). Empirically, we observed that the low-resolution method produced GASs that were nearly indistinguishable from those computed using the full dataset. This result held across multiple 10x Visium human tumor samples. Additional methodological details are provided in Supplementary Note 1, and benchmarking experiments validating the accuracy of the low-resolution GESSO approximation are described in Supplementary Note 2.

### Permutation-based local enrichment hypothesis testing for elevated pathway activity detection

To identify spatial locations with significantly elevated gene expression, we developed a permutation-based local activity enrichment testing procedure. The goal is to test the alternative hypothesis that a given spatial location exhibits significantly higher activity for the tested pathway compared to randomly generated pathways containing the same number of genes. Suppose we have a pathway with *G*_l_ genes. Our first step is to compute its gene set activity score at the spatial location of interest. Then, we randomly generate *n* combinations of *G*_l_ genes from the pool of *G* genes, yielding *n* control gene sets, and we compute their activity scores at the spatial location of interest. The control GASs form an empirical null distribution. The p-value is calculated as the proportion of control GASs that exceed the observed GAS for the actual pathway. This method directly accounts for pathway size and gene-level expression variability, providing a robust test for spatially distributed pathway activity scores. Full details are available in Supplementary Note 1.

### Simulation design

We developed a simulation framework to evaluate the performance of different methods for quantifying pathway activity scores. Following the strategy of the spatially variable gene detection method nnSVG^137^, we generated synthetic SRT data consisting of both spatially distributed genes (SGs) and randomly expressed genes (RGs). Spatial coordinates were obtained from the 10x Visium human dorsolateral prefrontal cortex DLPFC dataset^38^, which contains 3,611 spatial locations, of which 513 correspond to the white matter (WM) region, which forms a biologically distinct compartment with clear spatial boundaries in the DLPFC.

To generate realistic spatial expression patterns, we estimated distributional parameters from the real DLPFC data. Specifically, we first calculated the sample mean (µ) and sample variance (σ^2^) of log-transformed expression values for the white-matter marker gene *MOBP* within and outside the white-matter region: 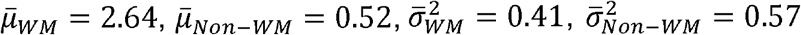. For each spatially distributed gene *g* ∈ *G*_*SG*_ expression values were generated according to

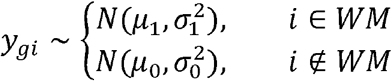

Where 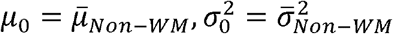, and 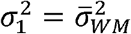. The mean gene expression inside the WM region is defined as

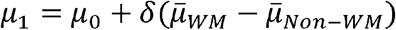

With *δ* = 1 and *δ* = 0.5 representing strong and weak spatial contrasts between WM and non-WM regions, respectively.

For each randomly distributed gene *g* ∈ *G*_*RG*_, expression values were drawn independently across all spatial locations. To mimic realistic random expression patterns, we parameterized the simulation using the housekeeping gene *GAPDH*^138,139^, whose log-transformed expression across the DLPFC yielded a mean of 3.3 and variance of 0.17 (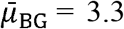 and 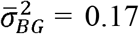). The gene-specific mean *μ*_g_ was then sampled from a uniform distribution between the non-white-matter baseline and the GAPDH mean, 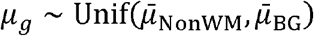 and expression at each spatial location *i* was generated as

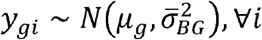

In total, we simulated 5000 genes, comprising 1000 SGs (500 strong, 500 weak SGs) and 4000 RGs. To mimic sparsity observed in real SRT data, we randomly dropped simulated gene expression at rates of 60% (e.g., 10x Visium level), 70%, 80% (e.g., Stereo-seq level), 90% (e.g., CosMx, Visium HD and Xenium Prime levels) expression for each gene’s expression over the whole tissue sample (see Supplementary Table 2). We then constructed gene sets with varying degrees of spatial enrichment following the GSDensity^37^ framework. Each gene set contained a mixture of SGs and RGs, with proportions of SGs ranging from 15%, 30%, 50%, to 75%, and total sizes from 50, 100, to 200 genes. For each setting, 10 replicate gene sets were generated by random sampling.

We benchmarked each method by evaluating its ability to distinguish the WM from non-WM regions. For each scenario, we used method-specific GAS values for each designated gene set in a logistic regression model to classify the binary WM label. Performance was quantified using AUC, F1 score, and overall accuracy, with F1 and accuracy computed at a fitted probability threshold of 0.5. These metrics match those used in our real-data benchmarking, and their calculation is detailed in the *Gene set activity-based prediction analysis* section.

### Compared methods

We compared GESSO with four gene set level activity scoring methods: (1) AUCell (R package version 1.26.0) (2) GSVA (R package version 1.52.3) (3) ssGSEA (an option built in GSVA package) (4) GSDensity (R package 0.1.3). Details for these four competing methods were in Supplementary Note 3. The input for the compared methods is a count or normalized count expression matrix and a list of gene sets. For AUCell and GSDensity, we followed the tutorials on the corresponding GitHub pages and used the recommended default parameter settings in their tutorials. For GSVA and ssGSEA, we applied the built-in algorithms in GSVA R package with the tutorial-suggested defaults. All four methods produced gene-set-by-location activity score matrices as their primary outputs.

### Real data analysis

To demonstrate the performance and biological interpretability of GESSO in real-world settings, we analyzed 13 publicly available spatial transcriptomics datasets spanning 10x Visium, Visium HD, Xenium Prime, CosMx, and Stereo-seq platforms. Datasets covered tumors, developmental tissues, lymphoid organs, and human cortex; full details, including sample sources, preprocessing, and details about the gene sets we analyzed, are provided in Supplementary Note 4 and Supplementary Table 1 and 4. Our comprehensive evaluation included the following types of analyses:

#### Gene set activity-based prediction analysis

GESSO produces gene set activity scores (GASs) that capture the dominant spatial expression pattern of each gene set and can distinguish locations with different functional or cellular compositions. To evaluate the predictive utility of GASs, we compared GESSO with AUCell, GSVA, ssGSEA, and GSDensity across all datasets for predicting biologically relevant labels at each spatial location. We reason that gene set activity methods producing more biologically meaningful scores will yield better predictions of the known labels for each spatial location. Specifically, spatial labels were obtained from the original studies for the Stereo-seq, Visium HD, and CosMx datasets; for the 10x Visium BRCA and Xenium Prime datasets, we manually annotated regions by overlaying histology or protein-stained images with spatial coordinates. For each dataset, spatial locations were split into 80% training and 20% test sets. Logistic regression models were trained using the GASs from each method as features. For multiclass classification tasks (all datasets except Xenium Prime), we used a one-vs-rest logistic regression framework, which fits a separate classifier for each annotated region and enables direct identification of the GASs most predictive of each region. Model performance on the held-out test set was evaluated using three metrics: macro-averaged AUC (one-vs-one formulation), macro-averaged F1 score, and overall accuracy, computed using the *roc_auc_score(), f1_score*(), and *accuracy_score()* functions in scikit-learn.

#### Hypothesis testing workflow

We proposed a permutation-based hypothesis testing framework (Supplementary Note 1) to detect spatial locations with significantly elevated gene set activity. Specifically, for each pathway, we generated 2,000 random gene sets matched for size and computed GASs for both the original and random sets across all methods. The first 1,000 random sets were used to construct an empirical null distribution of GASs, from which one-sided p-values were obtained; the remaining 1,000 served as independent nulls to evaluate p-value calibration. Calibration was assessed globally and locally. Globally, aggregated null p-values across all spatial locations should follow a uniform distribution, indicating proper control of false positives. Locally, null p-values within each individual location should also approximate uniformity, ensuring that the test is not biased toward specific spatial regions. Global calibration was assessed using Q-Q plots and quantified via the genomic inflation factor *λ*_*GC*_, defined as the ratio between the median observed and expected test statistics under the null^141,142^; values near 1 indicate well-calibrated p-values. Local calibration was evaluated using the Kolmogorov-Smirnov statistic, which measures the deviation of per-location null p-values from the uniform (0,1) distribution.

To assess the biological relevance of the p-values produced by GESSO, as well as by GSVA, GSDensity, AUCell, and ssGSEA, we evaluated their correspondence with known spatial annotations. Specifically, we examined whether low p-values occurred in biologically meaningful regions, such as “Invasive Carcinoma” for tumor-specific pathways, immune-rich areas in 10x Visium datasets, or anatomically defined organs (e.g., brain, heart, liver, skeletal tissues) in the Stereo-seq embryo data. For each method, negative p-values were used as ranking scores for ROC analysis, and concordance with annotated regions was quantified using the area under the ROC curve (AUC). We additionally computed the Dice coefficient between significant locations (p < 0.05) and annotated regions, defined as 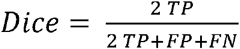, where *TP* denotes the number of true-positive spatial locations, *FP* denotes the number of false-positive spatial locations, and denotes the number of false-negative spatial locations. For these metrics, higher AUC and Dice values indicate better alignment between detected pathway activity and true biological structure.

#### Mapping cancer-immune spatial interactions in 10x Visium human tumor datasets

In the 10x Visium human tumor dataset, we qualitatively assessed whether GESSO-derived cancer and immune GASs could capture tumor-immune colocalization patterns consistent with those obtained from a widely used cell type deconvolution method for spatial transcriptomics, CARD^22^. Specifically, we applied CARD with a single-cell RNA-seq reference datasets for BRCA^143^ from Gene Expression Omnibus (GEO) repository (accession number GSE176078), CRC^144^ from GSE144735, LUSC^145^ from GSE131907, PRAD^146^ from GSE181294, and OVCA^147^ from https://data.mendeley.com/datasets/rc47y6m9mp/1 (Supplementary Table 1), to estimate the immune and tumor cell type proportions. GESSO-derived GASs were then compared with CARD-derived tumor and immune cell type proportions to assess whether GESSO accurately captured the spatial interplay between tumor and immune compartments.

#### Tracking developmental dynamics of organ gene set activity in Stereo-seq mouse embryo dataset

We analyzed organ-specific GASs across embryonic stages in the Stereo-seq dataset to assess how pathway activity relates to organ growth. Relative organ size was estimated as the proportion of spatial locations annotated to each organ. For each stage, we extracted GESSO-derived GASs for brain-, heart-, liver-, and skeletal-related gene sets within their corresponding annotated regions. For example, brain-related gene sets were evaluated within “Brain”-annotated locations, heart-related gene sets within “Heart,” liver-related gene sets within “Liver,” and skeletal-related gene sets within stage-specific skeletal annotations (e.g., “Cartilage primordium,” “Jaw and tooth,” “Connective tissue,” “Meninges,” “Dorsal root ganglion”). We then investigated the relationship between organ size and gene set activity scores. Finally, we embedded all organ-specific GASs into a UMAP^148^ space and stratified the embedding by developmental stage to visualize how organ-level transcriptional programs evolve over time.

#### Germinal center-focused analysis of transcriptional program gradients in the Xenium Prime human lymph node dataset

In the Xenium Prime human lymph node dataset, we defined three rectangular regions of interest (ROIs), hereafter red, blue, and green (Figure 6A). For every ROI, we investigated whether germinal center (GC) cells were enriched for particular transcriptional programs relative to all cells in the same spatial window. For each spatial location, we first obtained GESSO gene set activity scores for a B-cell progenitor signature (*HADDAD_B_LYMPHOCYTE_PROGENITOR*), a naive T-cell signature (*HAY_BONE_MARROW_NAIVE_T_CELL*) and an intercellular-transport signature (*GOBP_VESICLE_MEDIATED_TRANSPORT*). GASs were min–max scaled, and each location was assigned a dominant pathway label corresponding to the highest rescaled score. Within each ROI, we compared dominant pathway labels between GC and non-GC locations. We then investigated how dominant pathway composition varied with distance from the GC center. To do so, we selected one representative GC per ROI and computed the Euclidean distance from each cell in the ROI to the centroid of the nearest GC. Distances were normalized by the maximum GC radius to enable comparisons across GCs of different sizes. At each normalized distance, we estimated the probability that each pathway (B-cell, T-cell, or intercellular transport) was dominant.

These probabilities were smoothed using Gaussian kernel density estimation via *scipy*.*stats*.*gaussian_kde()* with its default parameters to generate continuous profiles of pathway dominance from GC cores to peripheral regions.

To further characterize the relationship between distance from the GC center and pathway dominance, we projected each cell’s relative B-cell, T-cell, and intercellular transport GASs into barycentric (ternary) space. Cells were then organized by their normalized distance to the nearest GC centroid, enabling inspection of how transcriptional program composition transitions across spatial gradients. These ternary analyses provided a complementary perspective on pathway coordination, revealing distance-dependent shifts in program dominance that are not apparent from pathway-wise summaries alone.

### Sensitivity analysis of the spatial smoothing strength parameter

To assess the impact of spatial regularization on GAS estimation, we evaluated GESSO across a range of smoothing strengths (*β* = 0, 0.05, 0.1, 0.2, 0.25, 0.33, and 1.0). As discussed in Supplementary Note 1, *β* = 0 corresponds to no spatial smoothing, while *β* = 1 reduces the optimization to the Laplacian eigenmaps formulation. For each *β*, we computed GASs for the simulated dataset and two representative real datasets—the 10x Visium BRCA sample and the Stereo-seq embryo data—spanning low to moderate spatial resolution. We did not extend this evaluation to the highest-resolution datasets (e.g., Visium HD and CosMx), as recomputing GASs for thousands of pathways across multiple values are computationally prohibitive at that scale. Importantly, our simulation framework was designed to mimic key characteristics of both low- and high-resolution technologies, allowing systematic assessment of smoothing effects across SRT settings. Performance was assessed using the gene set activity-based prediction framework for real data and white-matter association for simulated data. Across both simulated and real datasets, prediction accuracy and AUC increased monotonically as *β* increased up to 0.33 (Supplementary Figures S42-S44), indicating that moderate spatial smoothing substantially improves the biological informativeness of GESSO’s GASs while avoiding over smoothing associated with larger *β* values.

## Data availability

The details of all 13 datasets are provided in Supplementary Table 1. The datasets used in this work can be accessed through the following links: **10x Visium:** (1) human breast ductal carcinoma (BRCA), colorectal cancer (CRC), lung squamous cell carcinoma (LUSC), ovarian carcinoma (OVCA), cervical cancer (CESC), and prostate adenocarcinoma (PRAD) at https://www.10xgenomics.com/datasets; (2) human dorsolateral prefrontal cortex (DLPFC) reference data at https://research.libd.org/spatialLIBD/. **Visium HD:** (1) human colorectal cancer and matched adjacent normal tissue at https://www.ncbi.nlm.nih.gov/geo/query/acc.cgi?acc=GSE249344. **Xenium Prime:** (1) human lymph node dataset at https://www.10xgenomics.com/datasets/preview-data-xenium-prime-gene-expression. **Stereo-seq:** (1) mouse embryo stages (E11.5, E12.5, E13.5) at https://db.cngb.org/search/project/CNP0001515/. **NanoString CosMx:** (1) human non-small cell lung cancer (NSCLC) niches at https://nanostring.com/products/cosmx-spatial-molecular-imager/ffpe-dataset/nsclc-ffpe-dataset/.

## Code availability

The GESSO method is implemented as a Python package and is publicly available for community use and contribution at https://github.com/YMa-lab/GESSO.

## Author contributions statement

Y.M. conceived the idea and supervised the study. A.J.Y., C.T., and Y.M. developed the method and designed the experiments. A.J.Y. developed and implemented the GESSO software. C.T. designed and performed the simulation study. A.J.Y. and C.T. performed real data analysis, benchmarking experiments, and prepared the figures. A.J.Y, C.T., and Y.M. wrote the manuscript.

## Acknowledgements

This work was supported by National Institutes of Health (NIH) grant R35GM160372 to Y.M., and National Science Foundation (NSF) grants DBI-2526948 and IIS-2500960 to Y.M.

