## Supplementary Material for "Interpretable and scalable spatial gene set activity analysis with GESSO uncovers functional tissue architecture"

### Supplementary Figures

**Supplementary Figure S1.** Bar plots of accuracy and F1-score evaluating binary prediction performance for each method per each scenario in the simulation studies. A threshold of fitted probability equal to 0.5 was used to classify the predicted labels.

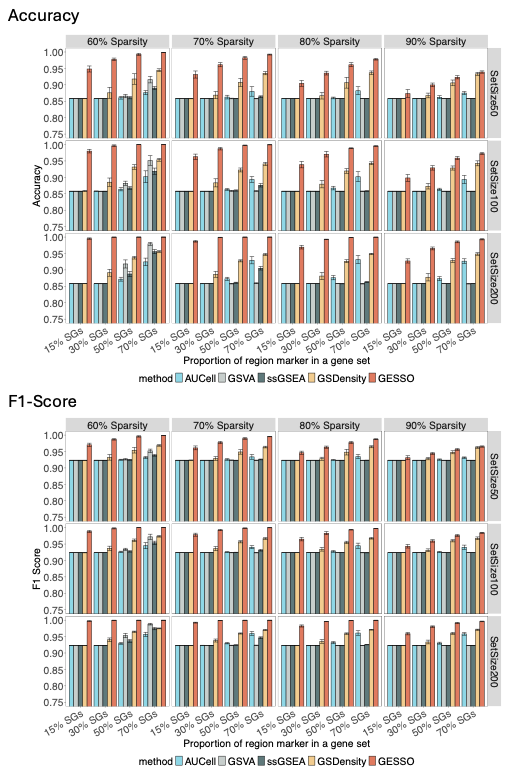

**Supplementary Figure S2.** Spatial maps of GAS calculated by all methods for gene sets with size 50, size 100 and size 200 when there are 60%, 70%, 80% 90% sparsity in the simulated data. GESSO results are highlighted by red color. For each scenario, GAS visualizations are organized by 15%, 30%, 50% and 70% markers from left to right.

**Supplementary Figure S3.** Quantile-quantile plots of –log10(*p*-values) from each method are shown under a null scenario for each sparsity level, each gene set and each marker proportion level in the simulated data. $\lambda_{gc}$ is the genomic control factor.

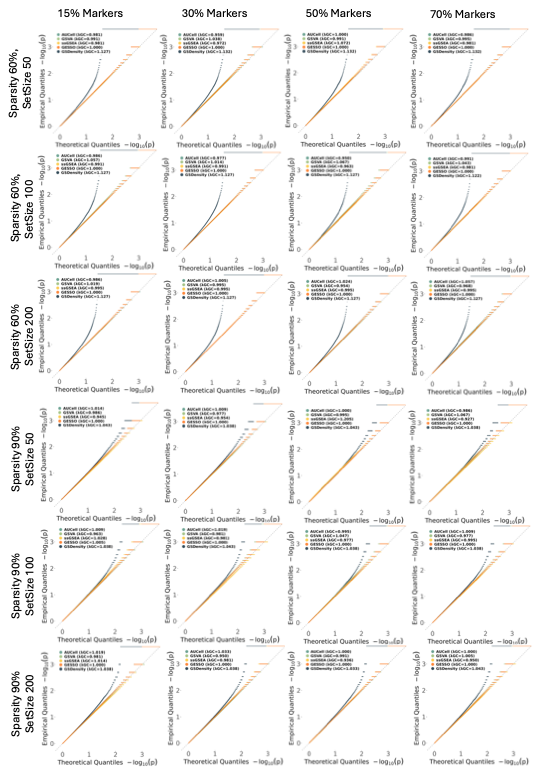

**Supplementary Figure S4.** Spatial map of Kolmogorov-Smirnov (K-S) statistics calculated from the distribution of p-values for tested control gene sets across tissue spots using GESSO in the simulated data. Lower K-S statistics indicate that the observed p-value distribution is closer to the expected uniform distribution under the null. Uniformly low K-S statistics across the tissue suggest proper false positive control, while spatial variation in K-S statistics indicates p-value inflation under the null.

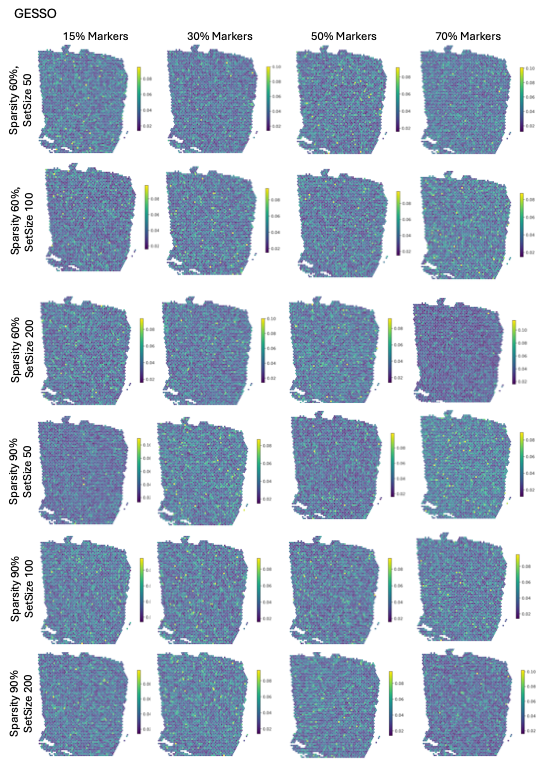

**Supplementary Figure S5.** Spatial map of Kolmogorov-Smirnov (K-S) statistics calculated from the distribution of p-values for tested control gene sets across tissue spots using ssGSEA in the simulated data. Lower K-S statistics indicate that the observed p-value distribution is closer to the expected uniform distribution under the null. Uniformly low K-S statistics across the tissue suggest proper false positive control, while spatial variation in K-S statistics indicates p-value inflation under the null.

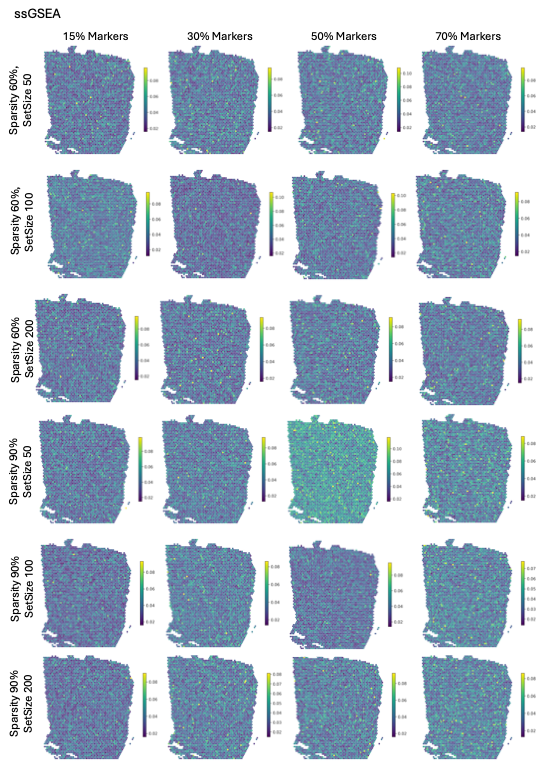

**Supplementary Figure S6.** Spatial map of Kolmogorov-Smirnov (K-S) statistics calculated from the distribution of p-values for tested control gene sets across tissue spots using GSVA in the simulated data. Lower K-S statistics indicate that the observed p-value distribution is closer to the expected uniform distribution under the null. Uniformly low K-S statistics across the tissue suggest proper false positive control, while spatial variation in K-S statistics indicates p-value inflation under the null.

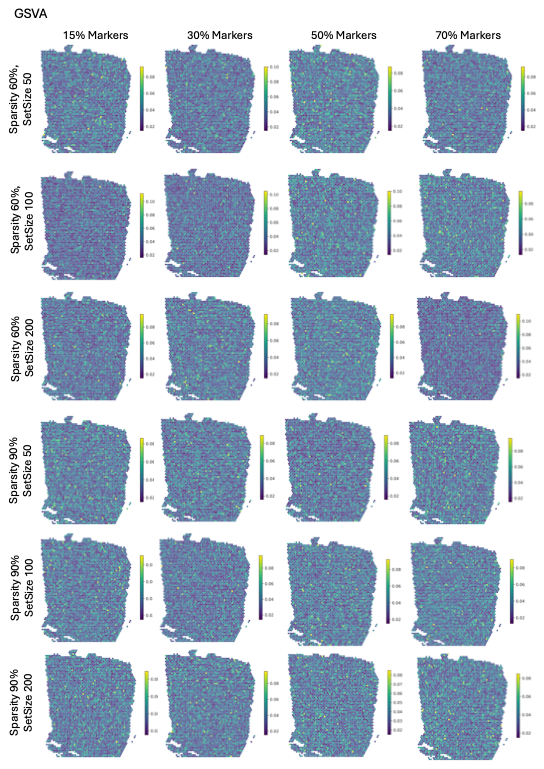

**Supplementary Figure S7.** Spatial map of Kolmogorov-Smirnov (K-S) statistics calculated from the distribution of p-values for tested control gene sets across tissue spots using AUCell in the simulated data. Lower K-S statistics indicate that the observed p-value distribution is closer to the expected uniform distribution under the null. Uniformly low K-S statistics across the tissue suggest proper false positive control, while spatial variation in K-S statistics indicates p-value inflation under the null.

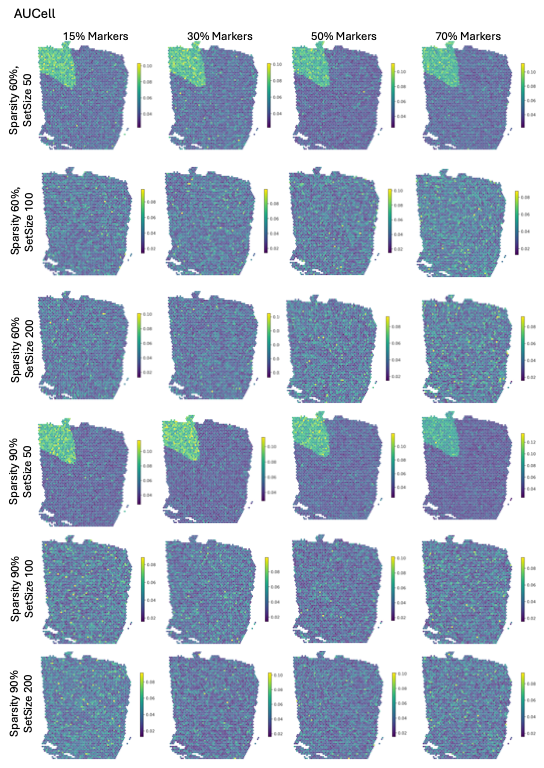

**Supplementary Figure S8.** Spatial map of Kolmogorov-Smirnov (K-S) statistics calculated from the distribution of p-values for tested control gene sets across tissue spots using GSDensity in the simulated data. Lower K-S statistics indicate that the observed p-value distribution is closer to the expected uniform distribution under the null. Uniformly low K-S statistics across the tissue suggest proper false positive control, while spatial variation in K-S statistics indicates p-value inflation under the null.

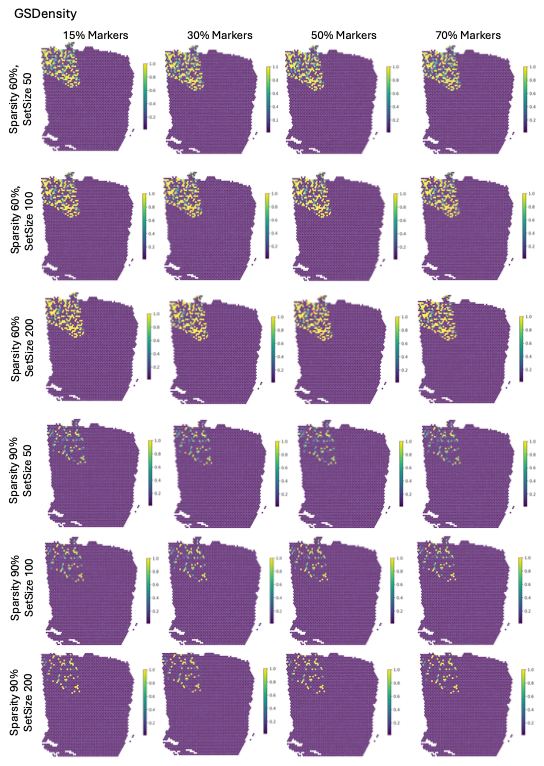

**Supplementary Figure S9.** Spatial map of p-values and significant spots for a representative gene set per gene set size and proportion of markers on the simulated data with 60% sparsity. The left-hand side are the p-values from the local activity-enrichment test, where lighter color indicates smaller p-values; the right-hand size are the maps of significant locations, where yellow indicates the location has p-value < 0.05.

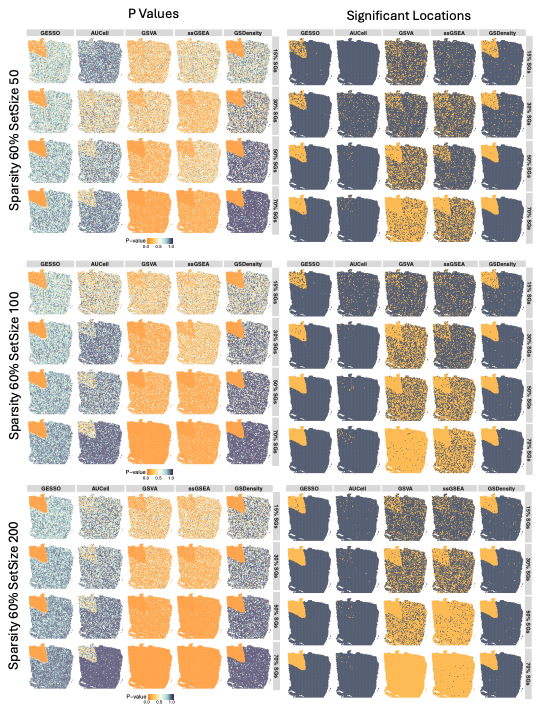

**Supplementary Figure S10.** Spatial map of p-values and significant spots for a representative gene set per gene set size and proportion of markers on the simulated data with 90% sparsity. The left-hand side are the p-values from the local activity-enrichment test, where lighter color indicates smaller p-values; the right-hand size are the maps of significant locations, where yellow indicates the location has p-value < 0.05.

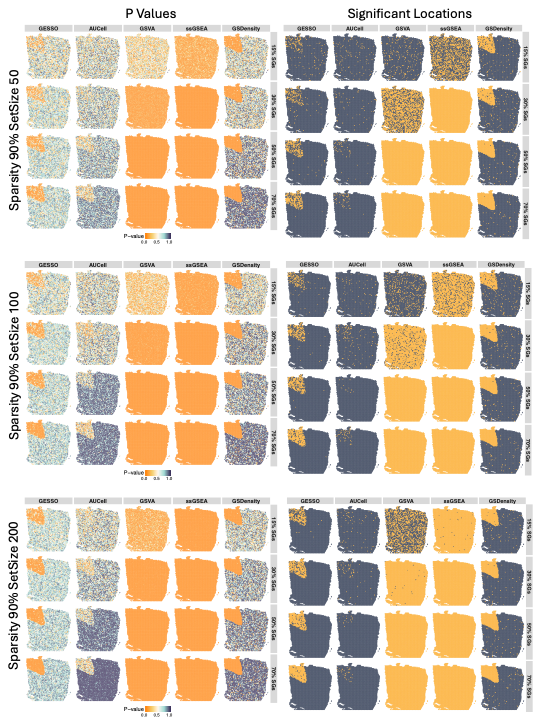

**Supplementary Figure S11.** Spatial concordance of significant GAS from representative gene sets with size 50 with the White Matter (WM) region, measured on the simulated data with 60% sparsity. For each setting of maker proportion, the first row shows the AUC scores obtained by using the p-values as decision scores to predict the WM region; the second row presents the Dice coefficients, calculated between the significant spatial locations (with a significance level of 0.05) and the masks of the WM region.

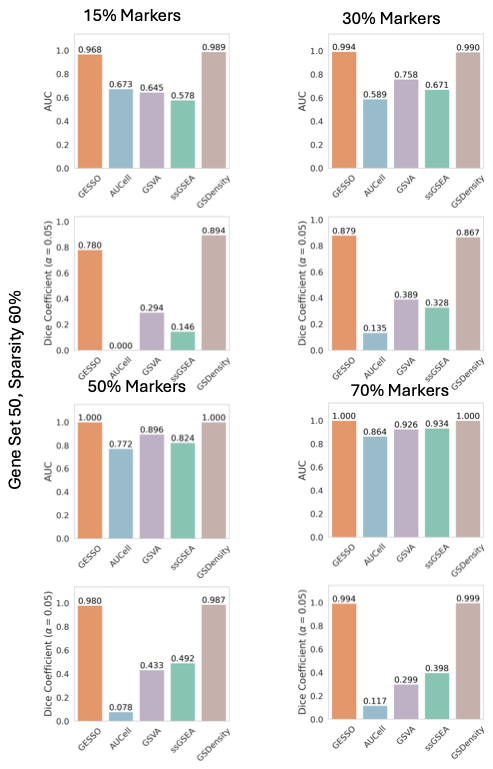

**Supplementary Figure S12.** Spatial concordance of significant GAS from representative gene sets with size 100 with the White Matter (WM) region, measured on the simulated data with 60% sparsity. For each setting of maker proportion, the first row shows the AUC scores obtained by using the p-values as decision scores to predict the WM region; the second row presents the Dice coefficients, calculated between the significant spatial locations (with a significance level of 0.05) and the masks of the WM region.

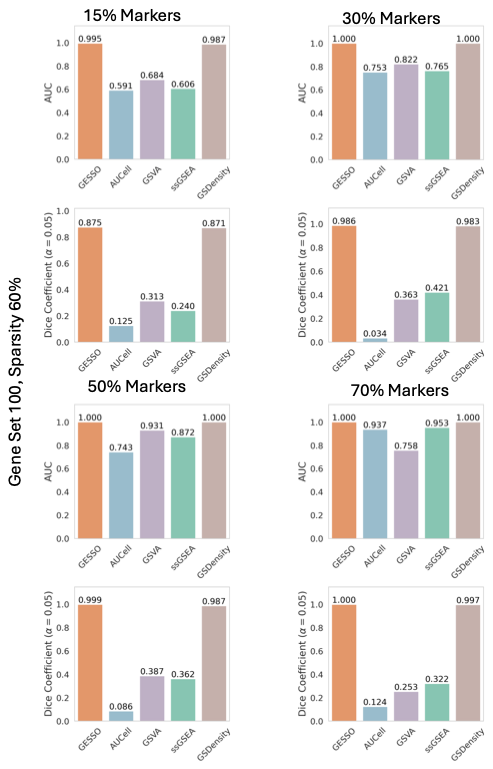

**Supplementary Figure S13.** Spatial concordance of significant GAS from representative gene sets with size 200 with the White Matter (WM) region, measured on the simulated data with 60% sparsity. For each setting of maker proportion, the first row shows the AUC scores obtained by using the p-values as decision scores to predict the WM region; the second row presents the Dice coefficients, calculated between the significant spatial locations (with a significance level of 0.05) and the masks of the WM region.

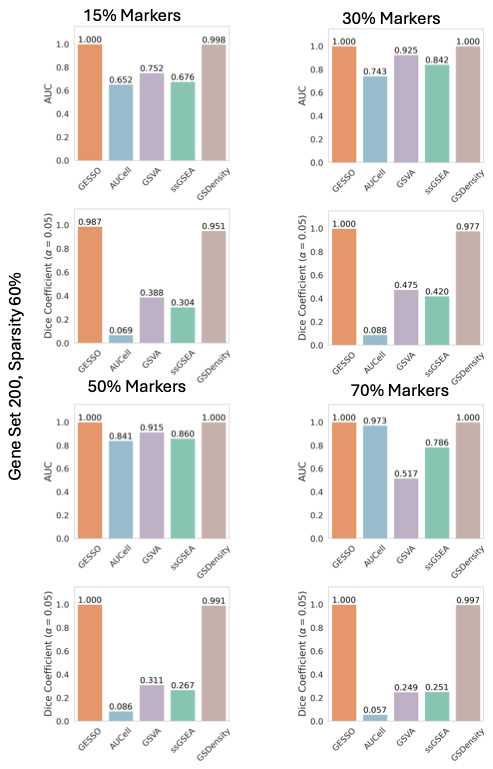

**Supplementary Figure S14.** Spatial concordance of significant GAS from representative gene sets with size 50 with the White Matter (WM) region, measured on the simulated data with 90% sparsity. For each setting of maker proportion, the first row shows the AUC scores obtained by using the p-values as decision scores to predict the WM region; the second row presents the Dice coefficients, calculated between the significant spatial locations (with a significance level of 0.05) and the masks of the WM region.

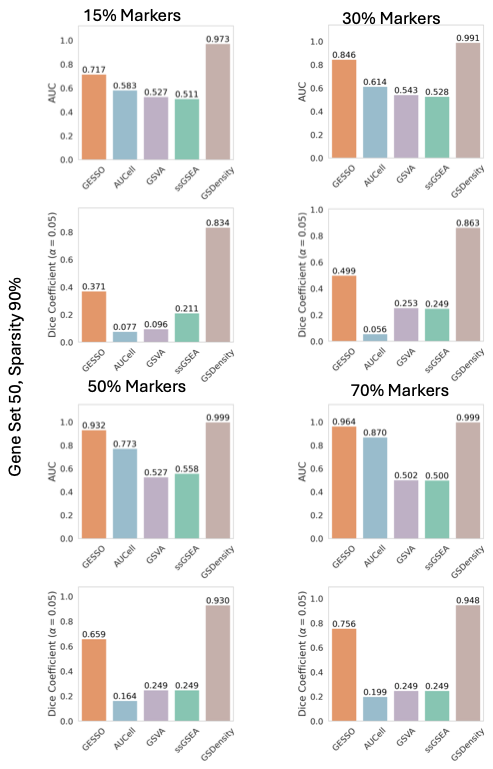

**Supplementary Figure S15.** Spatial concordance of significant GAS from representative gene sets with size 100 with the White Matter (WM) region, measured on the simulated data with 90% sparsity. For each setting of maker proportion, the first row shows the AUC scores obtained by using the p-values as decision scores to predict the WM region; the second row presents the Dice coefficients, calculated between the significant spatial locations (with a significance level of 0.05) and the masks of the WM region.

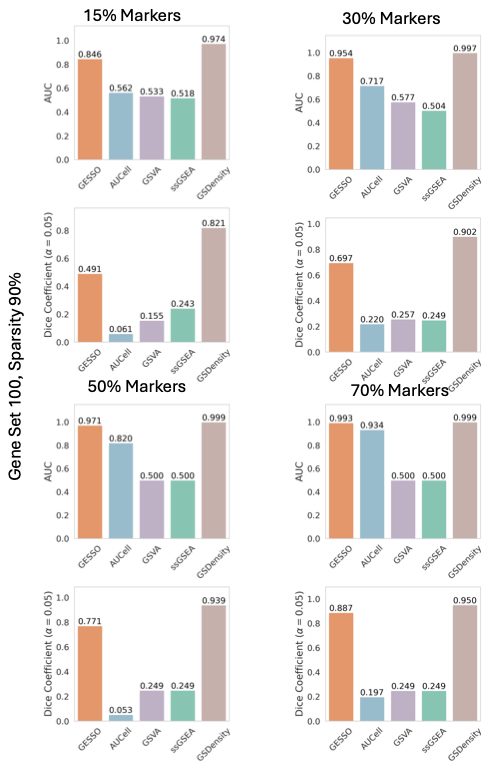

**Supplementary Figure S16.** Spatial concordance of significant GAS from representative gene sets with size 200 with the White Matter (WM) region, measured on the simulated data with 90% sparsity. For each setting of maker proportion, the first row shows the AUC scores obtained by using the p-values as decision scores to predict the WM region; the second row presents the Dice coefficients, calculated between the significant spatial locations (with a significance level of 0.05) and the masks of the WM region.

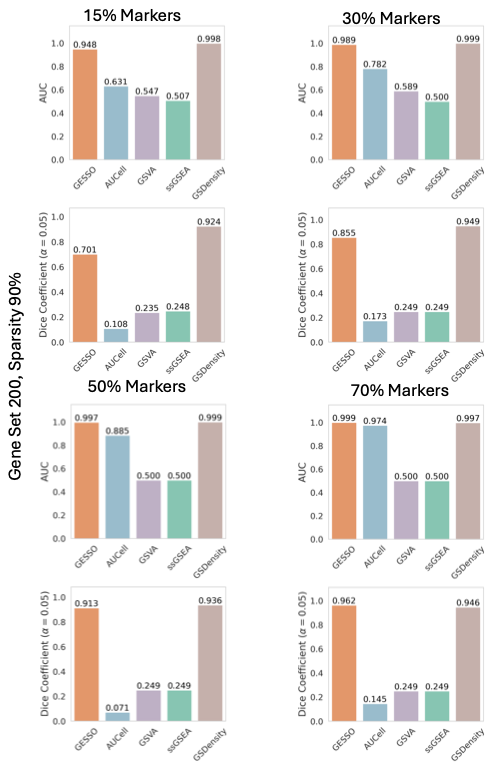

**Supplementary Figure S17.** H&E Image of 10x Visium BRCA data with annotated regions. The annotation map was downloaded from <https://www.10xgenomics.com/datasets/human-breast-cancer-ductal-carcinoma-in-situ-invasive-carcinoma-ffpe-1-standard-1-3-0>.

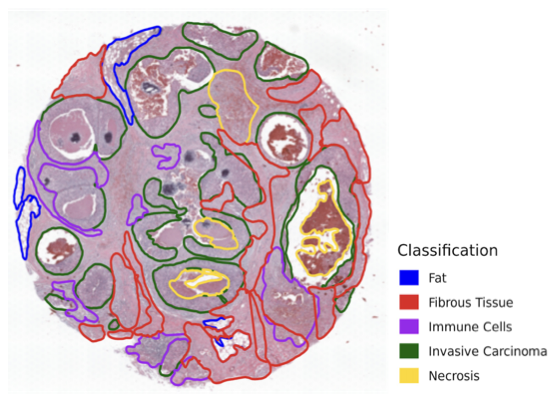

**Supplementary Figure S18.** Accuracy, macro-averaged one-vs-one AUC, and macro-averaged F1 scores of logistic regression classifiers fit to predict 10x Visium BRCA spot labels from the gene set activity scores generated by GESSO, AUCell, GSVA, ssGSEA, and GSDensity.

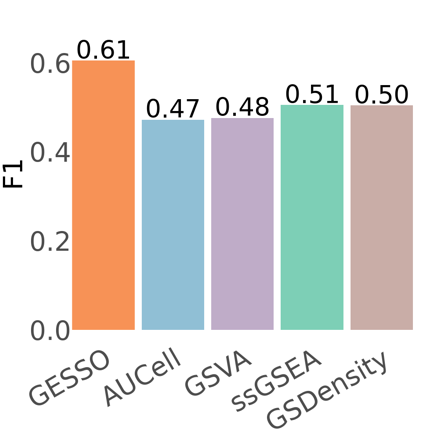

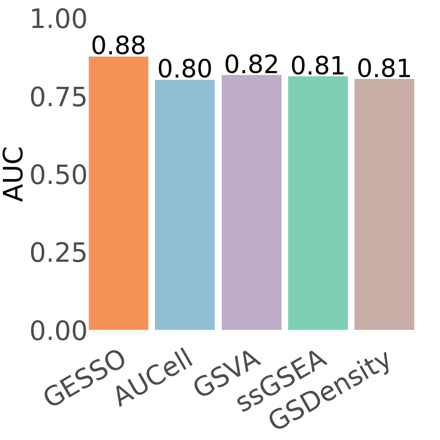

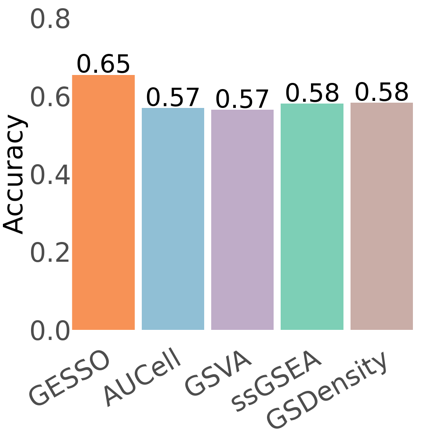

**Supplementary Figure S19.** Quantile-quantile plots of –log10(*p*-values) for each method are shown under a null scenario for cancer-specific and immune-specific pathway in 10x Visium tumor BRCA dataset. $\lambda_{gc}$ is the genomic control factor.

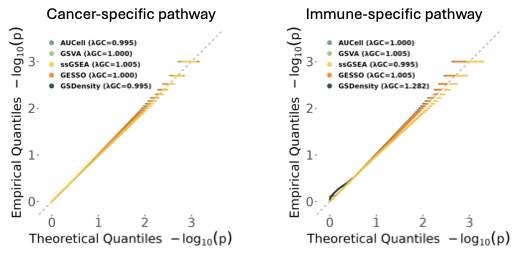

**Supplementary Figure S20.** Spatial maps showing the Kolmogorov-Smirnov statistic for 10x Visium BRCA data computed at each location for 1,000 randomly sampled gene sets, used to evaluate deviations from a uniform distribution. Lower K-S statistics indicate that the observed p-value distribution is closer to the expected uniform distribution under the null. Uniformly low K-S statistics across the tissue suggest proper false positive control, while spatial variation in K-S statistics indicates p-value inflation under the null

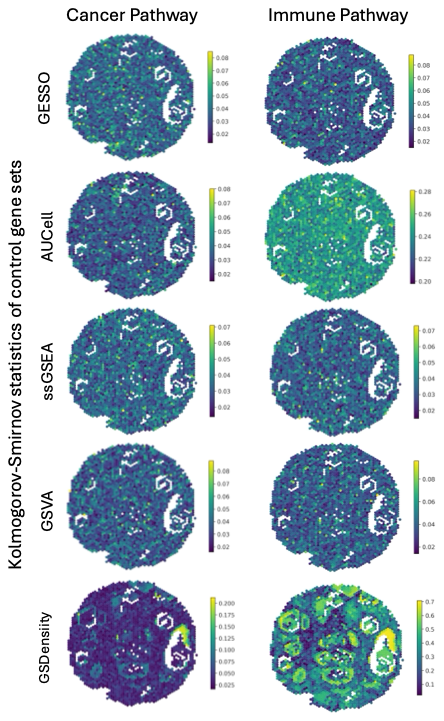

**Supplementary Figure S21.** Spatial concordance of elevated cancer-specific and immune-specific pathway expression with corresponding 10x Visium BRCA cancer- and immune- labeled regions. The left column depicts metrics computed using the p-values derived from the permutation-based hypothesis test based on gene set activity scores generated by GESSO, AUCell, GSVA, ssGSEA, and GSDensity for the cancer pathway (see Methods). The right column represents the same for the immune-specific pathway. The first row shows the AUC scores obtained by using the p-values as decision scores to predict cancer- and immune-labeled regions. The second row presents the Dice coefficients, calculated between the significant spatial locations (with a significance level of 0.05) and the masks of cancer and immune regions.

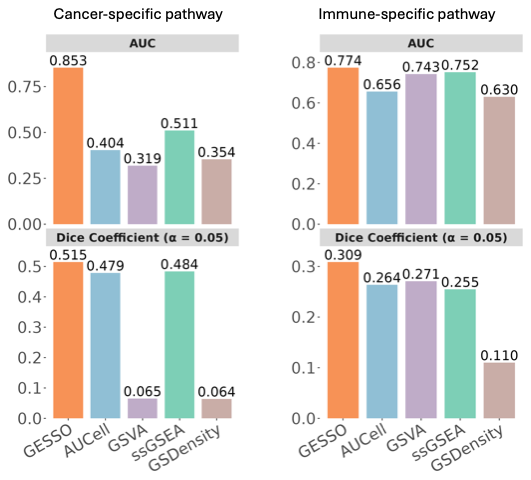

**Supplementary Figure S22.** Accuracy, macro-averaged one-vs-one AUC, and macro-averaged F1 scores of logistic regression classifiers fit to predict E11.5, E12.5, and E13.5 Stereo-seq mouse embryo stage spot labels from the gene set activity scores generated by GESSO, AUCell, ssGSEA, and GSDensity.

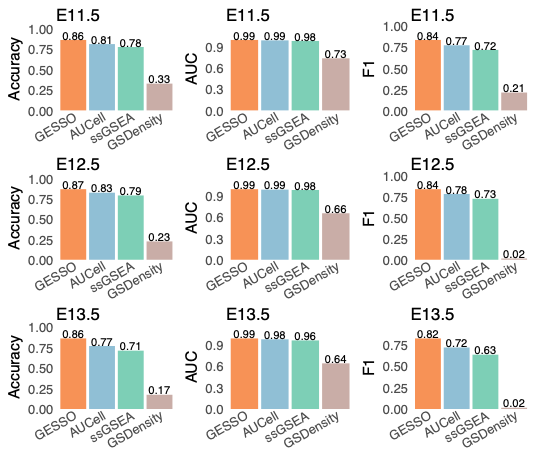

**Supplementary Figure S23.** Quantile-quantile plots of –log10(*p*-values) for each method are shown under a null scenario for the heart, brain, skeletal, and liver-specific pathways of the Stereo-seq E12.5 mouse embryo dataset. $\lambda_{gc}$ is the genomic control factor.

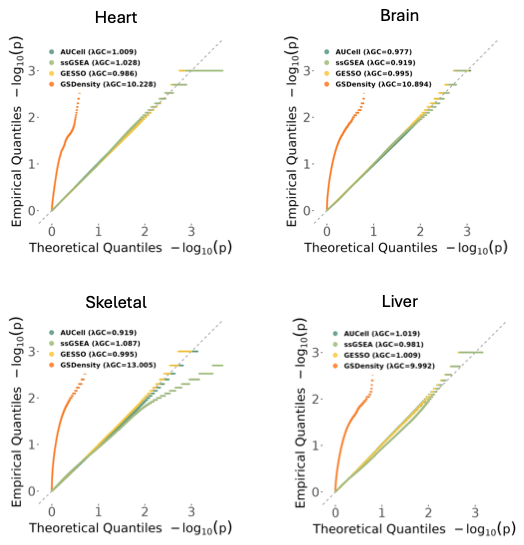

**Supplementary Figure S24.** Spatial maps showing the Kolmogorov-Smirnov statistic computed at each location in Stereo-seq Mouse Embryo E12.5 for 1,000 randomly sampled gene sets, used to evaluate deviations from a uniform distribution. Heart, Brain, Skeletal and Liver represents their specific pathway.

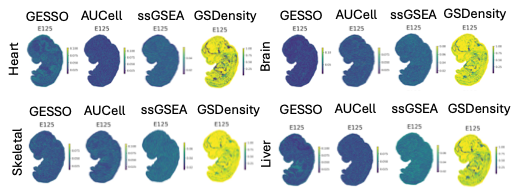

**Supplementary Figure S25.** Spatial concordance of elevated pathway expression with corresponding labels in the Stereo-seq E12.5 mouse embryo dataset. Each column depicts metrics computed using the p-values derived from the permutation-based hypothesis test based on gene set activity scores generated by GESSO, AUCell, GSVA, ssGSEA, and GSDensity for a specific pathway. The first row shows the AUC scores obtained by using the p-values as decision scores. The second row presents the Dice coefficients, calculated between the significant spatial locations (with a significance level of 0.05) and the masks of cancer and immune regions.

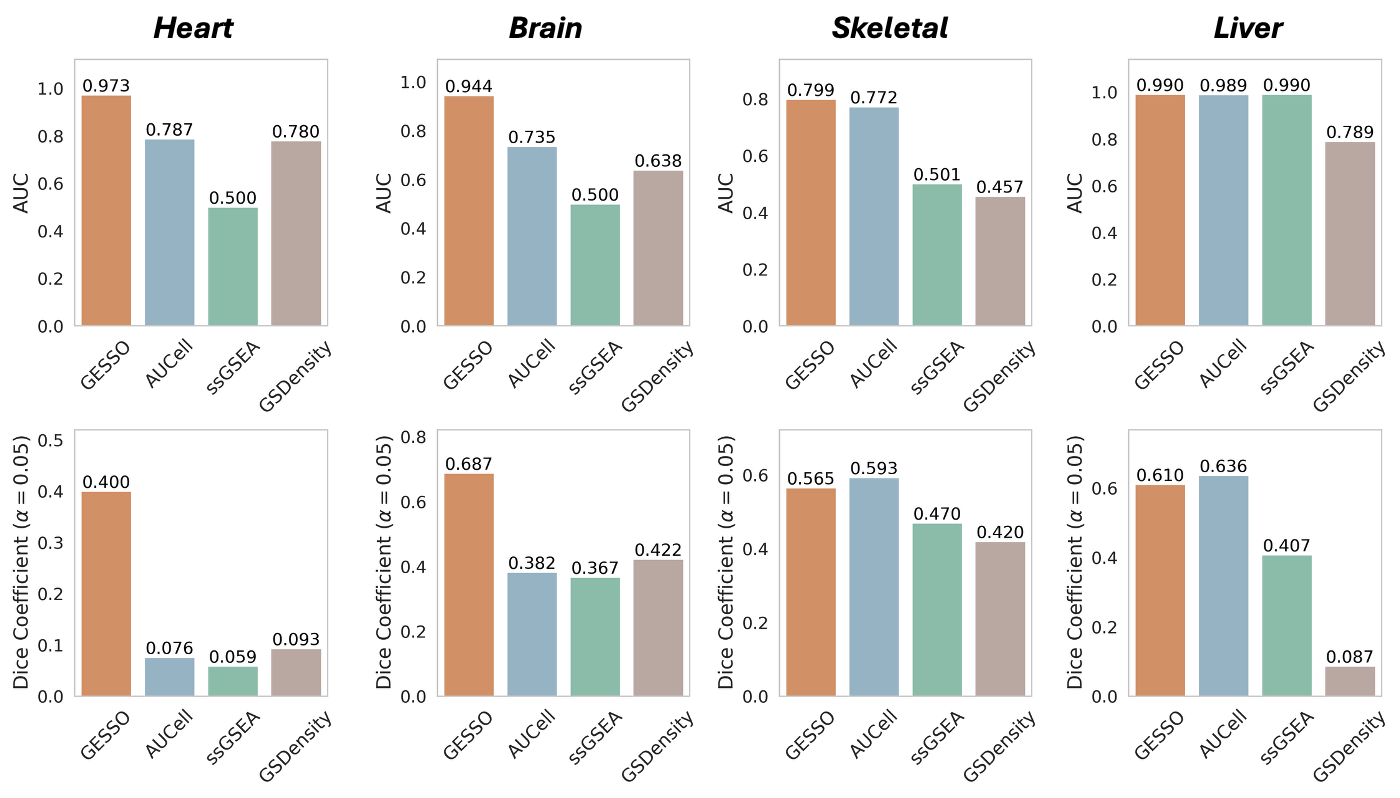

**Supplementary Figure S26.** Bar plots of accuracy, macro-averaged one-vs-one AUC, and macro-averaged F1 scores of logistic regression classifiers fit to predict Visium HD Tumor data cell type label from the GAS generated by GESSO, AUCell, and ssGSEA.

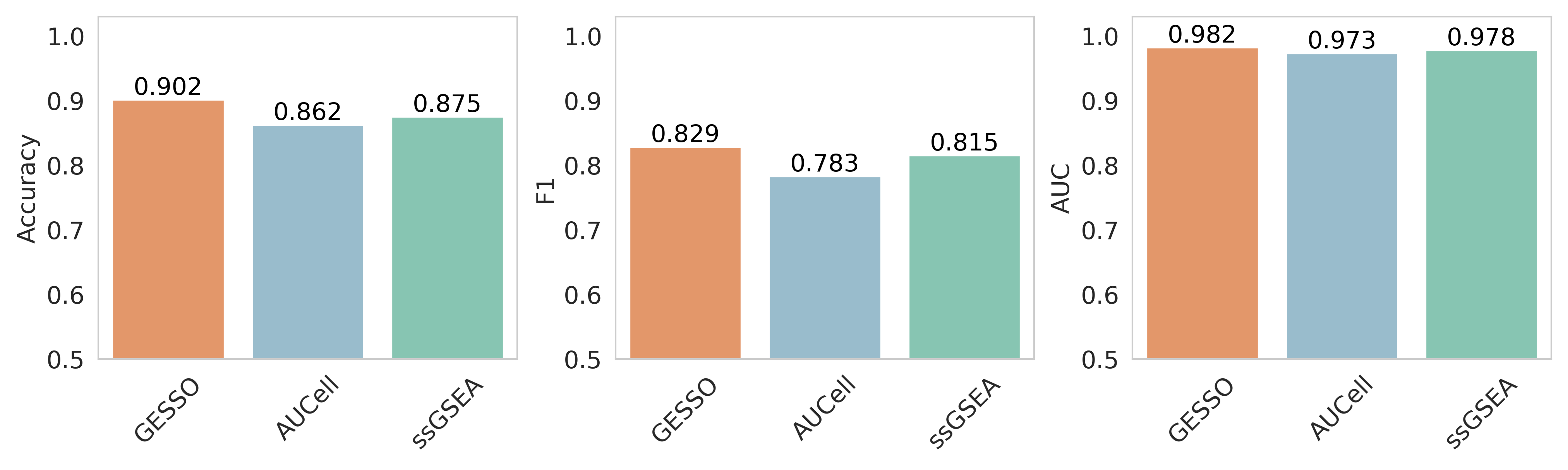

**Supplementary Figure S27.** Quantile-quantile plots of –log10(*p*-values) for each method are shown under a null scenario for the intestinal-epithelial-, T-cell-, Tumor-, and Fibroblast-specific pathways of the Visium HD CRC dataset. $\lambda_{gc}$ is the genomic control factor.

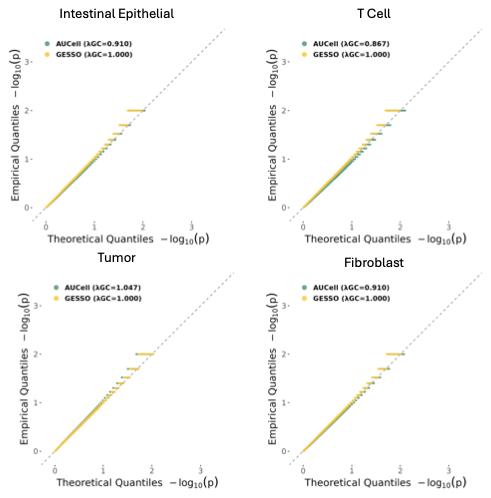

**Supplementary Figure S28.** Spatial maps showing the Kolmogorov-Smirnov statistic computed at each location of Visium HD human CRC data for 1,000 randomly sampled gene sets, used to evaluate deviations from a uniform distribution.

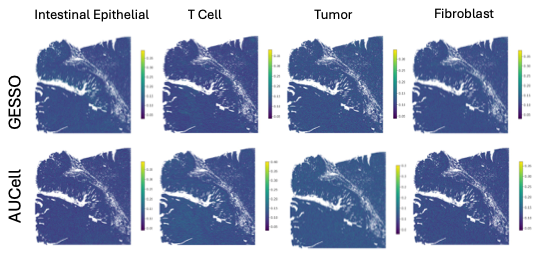

**Supplementary Figure S29.** Spatial map of spots showing significant cell-type-specific GAS. Yellow color indicates a spot is significant under *p*-value < 0.05.

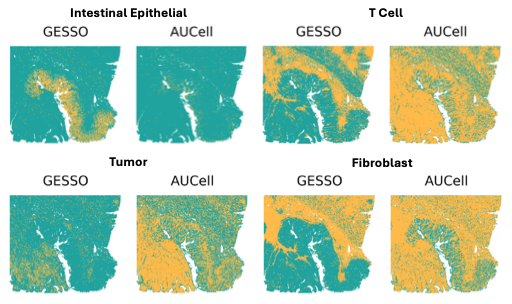

**Supplementary Figure S30.** Spatial concordance of elevated pathway expression with corresponding labels in the Visium HD human CRC dataset. Each column depicts metrics computed using the p-values derived from the permutation-based hypothesis test based on gene set activity scores generated by GESSO AND AUCell for a specific pathway. The first row shows the AUC scores obtained by using the p-values as decision scores. The second row presents the Dice coefficients, calculated between the significant spatial locations (with a significance level of 0.05) and the masks of pathway’s corresponding regions.

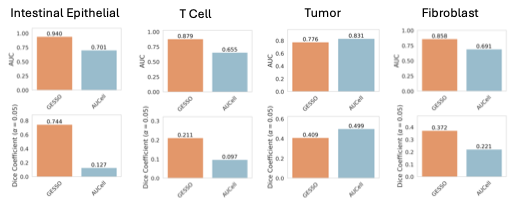

**Supplementary Figure S31.** Heatmaps presenting Pearson’s correlation coefficient between GESSO’s GASs in the tumor sample versus the control sample. The red block highlights the associations between tumor relevant gene set activity scores. The blue block highlights the associations between intestinal-epithelial relevant gene set activity scores. The brown block highlights the associate between activity scores of gene sets from the red and the yellow block.

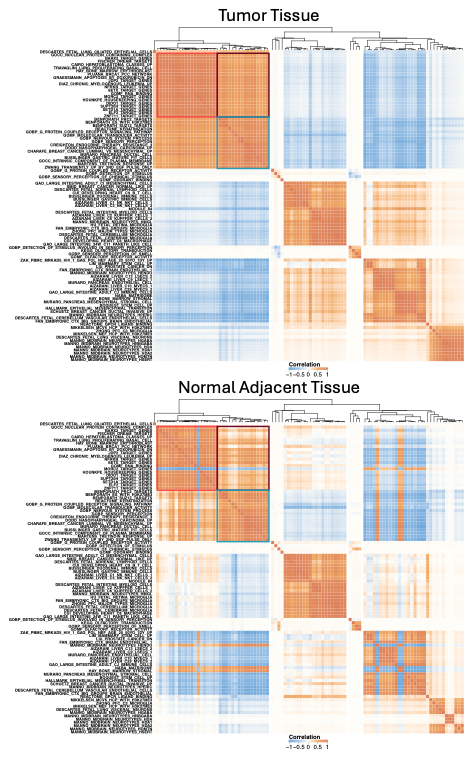

**Supplementary Figure S32.** H&E Image of 10x XeniumPrime Lymph node downloaded from <https://www.10xgenomics.com/datasets/preview-data-xenium-prime-gene-expression>.

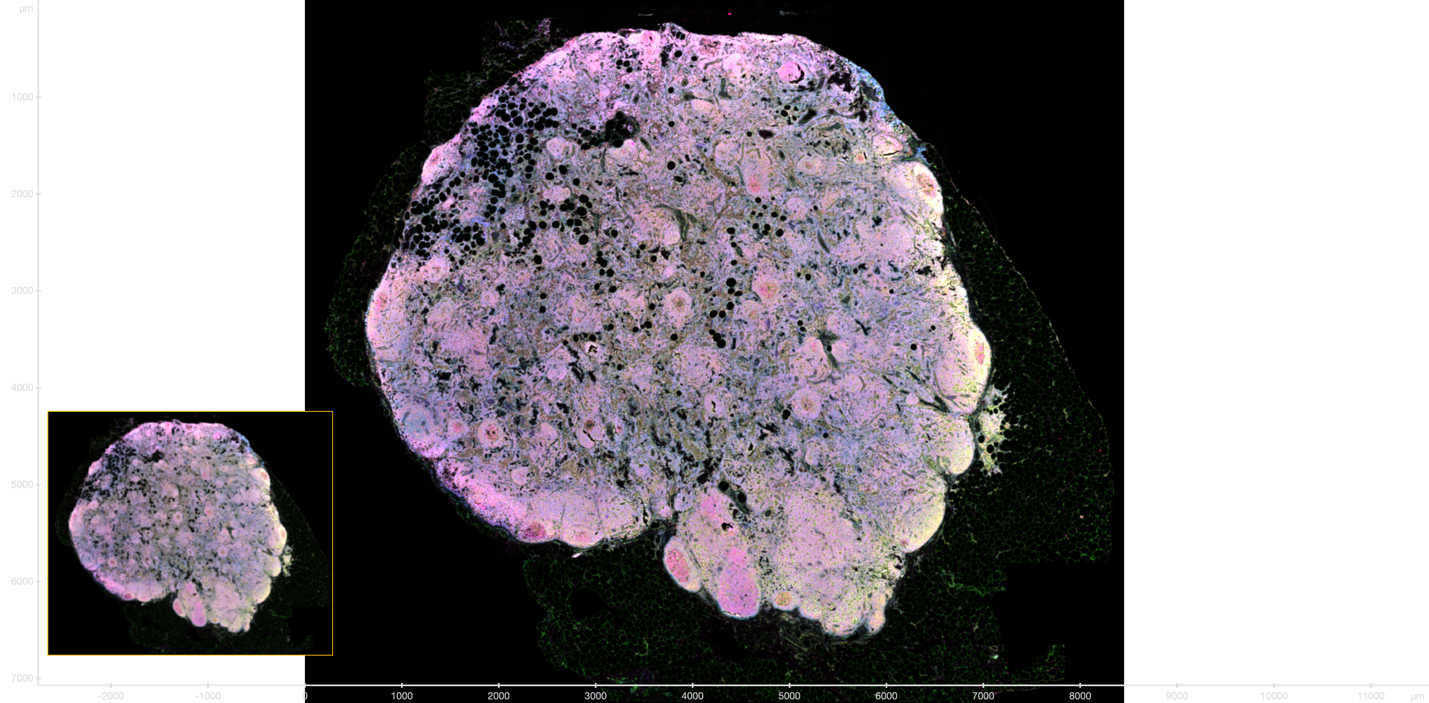

**Supplementary Figure S33.** Accuracy, weighted F1 scores, and AUC of logistic regression classifiers fit to predict the germinal centers in the 10x XeniumPrime human lymph node dataset from the gene set activity scores generated by GESSO, AUCell, and ssGSEA.

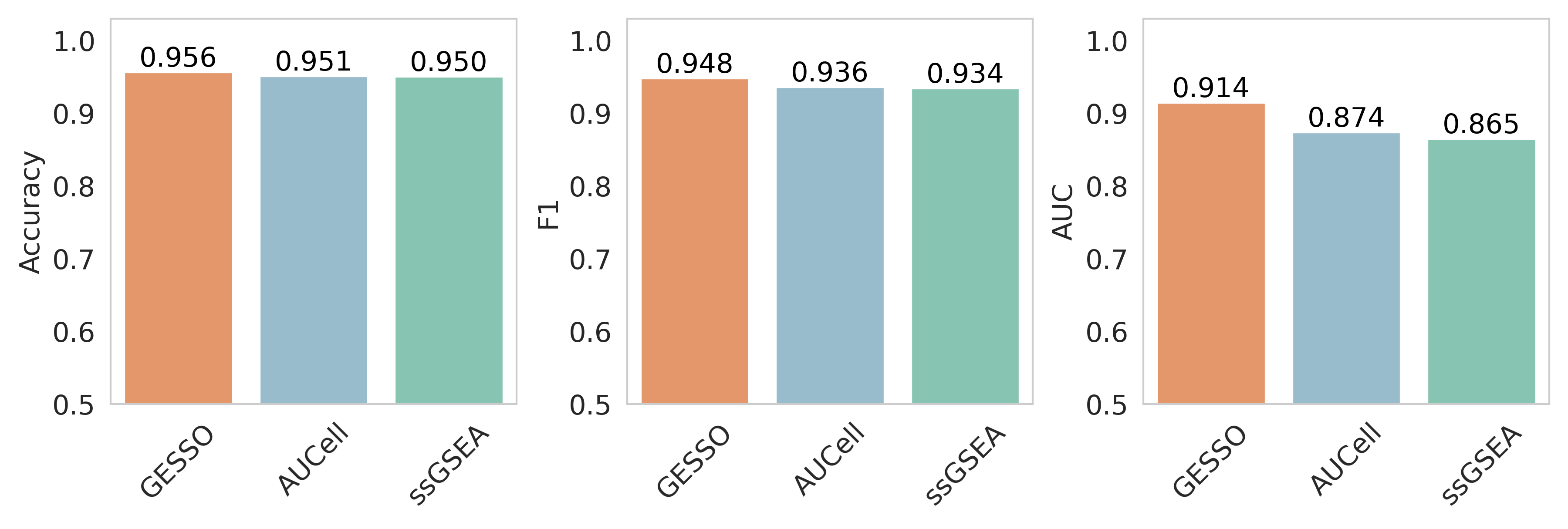

**Supplementary Figure S34.** Results of the permutation-based testing applied to the XeniumPrime human lymph node dataset. **A.** Quanti**l**e-quantile plots of –log10(p-values) for GESSO are shown under a null scenario for the lymphocyte-activation-, B-cell-, T-Cell-, and intercellular-specific pathways. $\lambda_{gc}$ is the genomic control factor. **B.** Spatial maps showing the Kolmogorov-Smirnov statistic computed at each location for 1,000 randomly sampled gene sets, used to evaluate deviations from a uniform distribution. **C.** Spatial map of spots showing significant GAS. Blue color indicates a spot is significant under *p*-value < 0.05. **D.** Spatial concordance of elevated pathway expression with the GC regions. Each column depicts metrics computed using the p-values derived from the permutation-based hypothesis test based on gene set activity scores generated by GESSO for a specific pathway. The first row shows the AUC scores obtained by using the p-values as decision scores. The second row presents the Dice coefficients, calculated between the significant spatial locations (with a significance level of 0.05) and the masks of the GC regions.

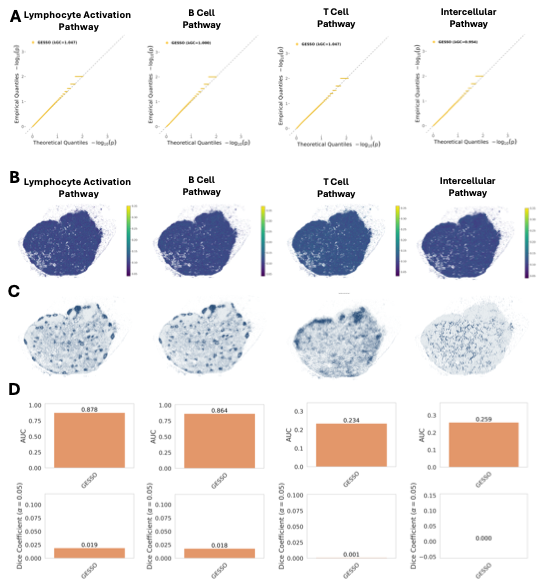

**Supplementary Figure S35.** Bar plots of accuracy, macro-averaged one-vs-one AUC, and macro-averaged F1 scores of logistic regression classifiers fit to predict CosMx NSCLS data niches label from the GAS generated by GESSO, AUCell, ssGSEA and GSDensity.

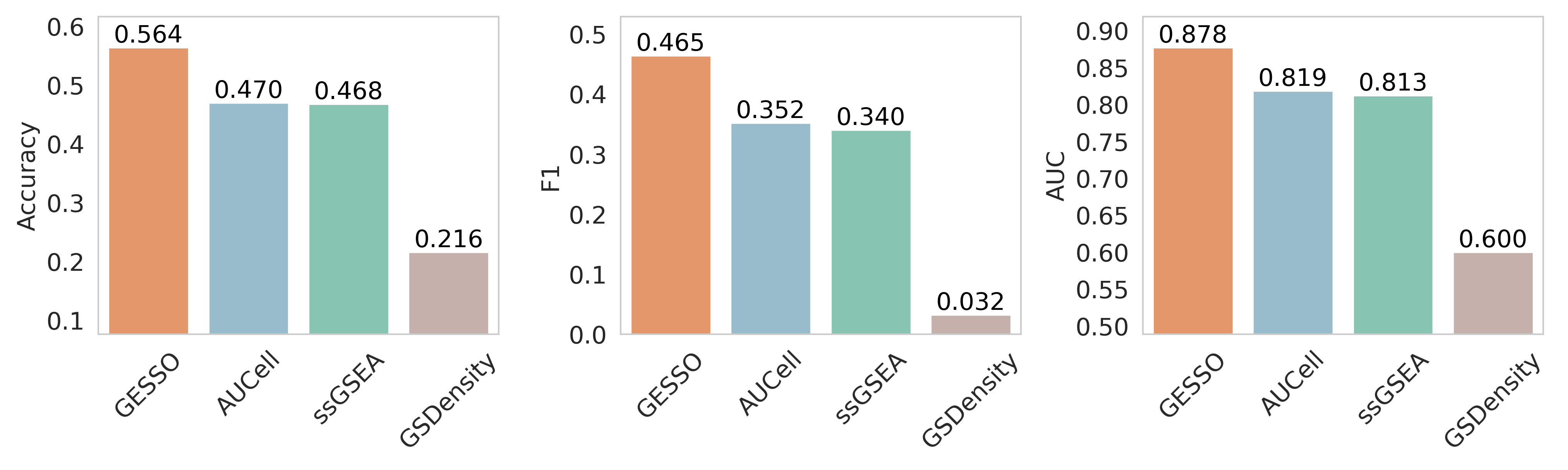

**Supplementary Figure S36.** Quantile-quantile plots of –log10(p-values) are shown under a null scenario for testing the niches-specific pathway of the CosMx human non-small cell lung cancer dataset. $\lambda_{gc}$ is the genomic control factor.

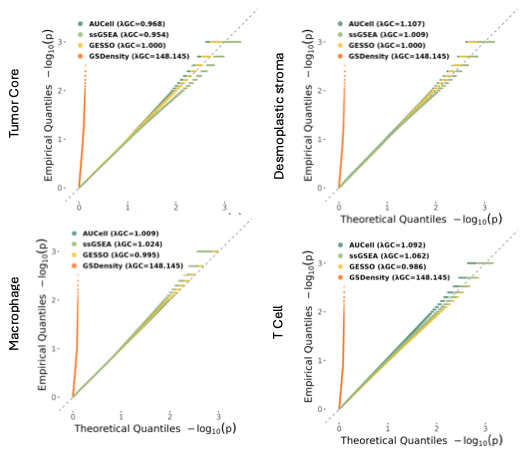

**Supplementary Figure S37.** Spatial maps showing the Kolmogorov-Smirnov statistic computed at each location of CosMx NSCLC for 1,000 randomly sampled gene sets, used to evaluate deviations from a uniform distribution**.**

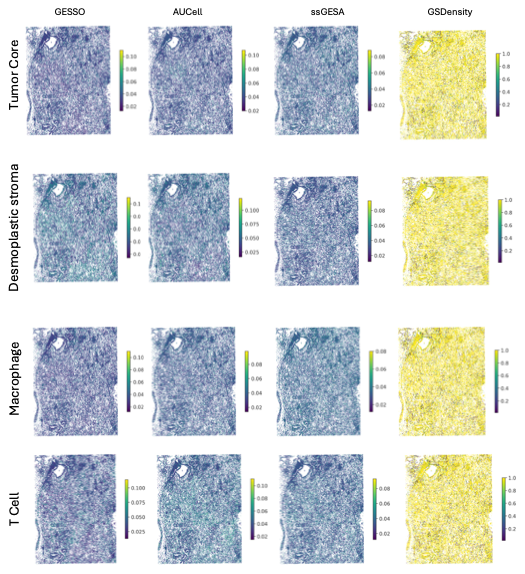

**Supplementary Figure S38.** Spatial map of spots showing significant niche-specific GAS. Red color indicates a spot is significant under p-value < 0.05.

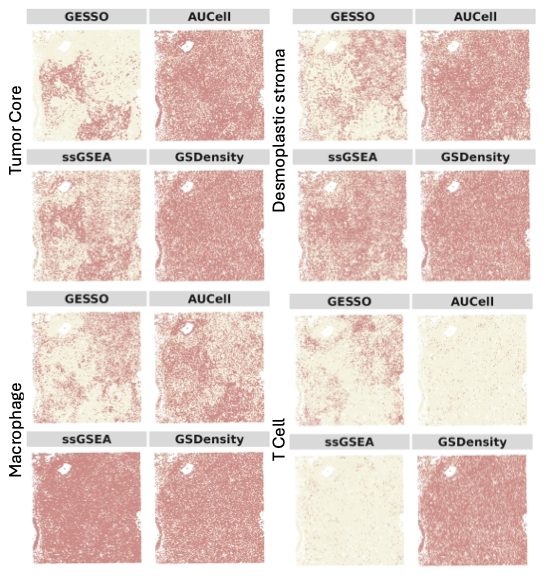

**Supplementary Figure S39.** Spatial concordance of significantly elevated tumor core-, desmoplastic stroma-, T cell, macrophage, and EMT-specific pathway expression with corresponding CosMx ECM tumor core- and desmoplastic stroma-labeled regions. Each subfigure depicts metrics computed using the p-values derived from the permutation-based hypothesis test based on GASs generated by GESSO, AUCell, ssGSEA, and GSDensity. The first row shows the AUC scores obtained by using the p-values as decision scores to predict associated labeled regions. The second row presents the Dice coefficients, calculated between the significant spatial locations (with a significance level of 0.05) and the masks associated labeled regions.

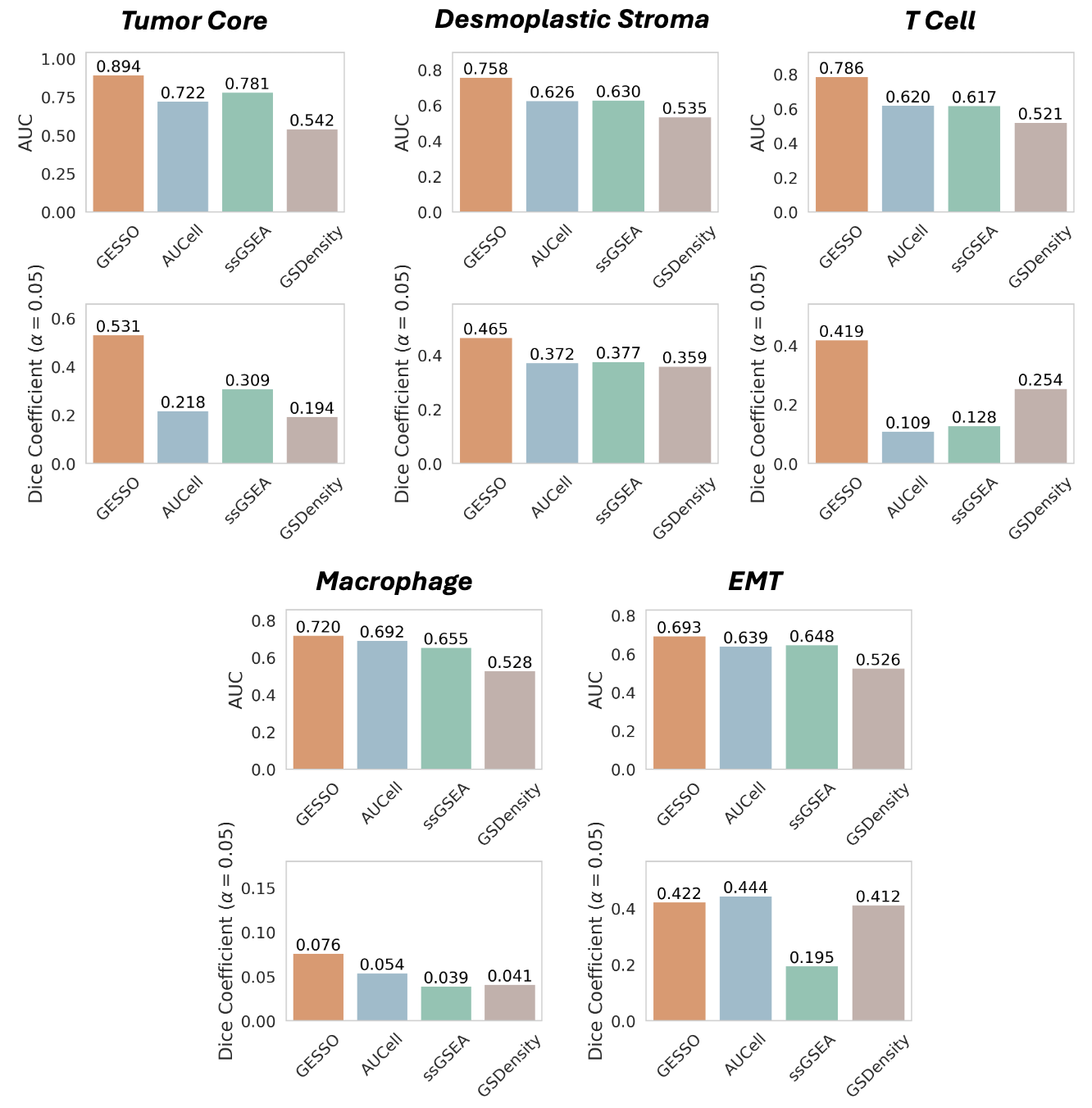

**Supplementary Figure S40.** **A**. Bar plots displaying the top 5 positive and negative metagene weights from GESSO’s EMT-specific GAS. **B**. Quantile-quantile plots of –log10(*p*-values) are shown under a null scenario for testing the EMT-specific pathway. $\lambda_{gc}$ is the genomic control factor. **C.** Spatial maps showing the Kolmogorov-Smirnov statistic computed at each location for 1,000 randomly sampled gene sets, used to evaluate deviations from a uniform distribution. **D**. Spatial map of spots showing significant EMT-specific GAS. Red color indicates a spot is significant under *p*-value < 0.05. **E.** Bivariate plot showing the co-location patterns between GESSO’s GAS of the desmoplastic-stroma-specific and the EMT-specific pathways.

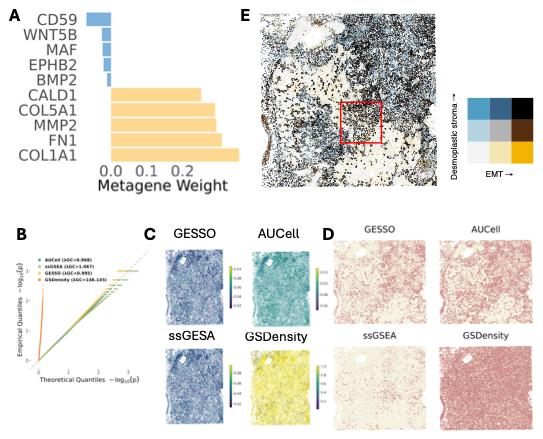

**Supplementary Figure S41.** **A.** Root mean squared error (RMSE) between low-resolution GASs generated using different partitioning strategies (method and number of partitions) and the reference GESSO GASs (see Supplementary Note 2). **B.** Pearson correlation between the same low-resolution GASs and the reference GESSO GASs.

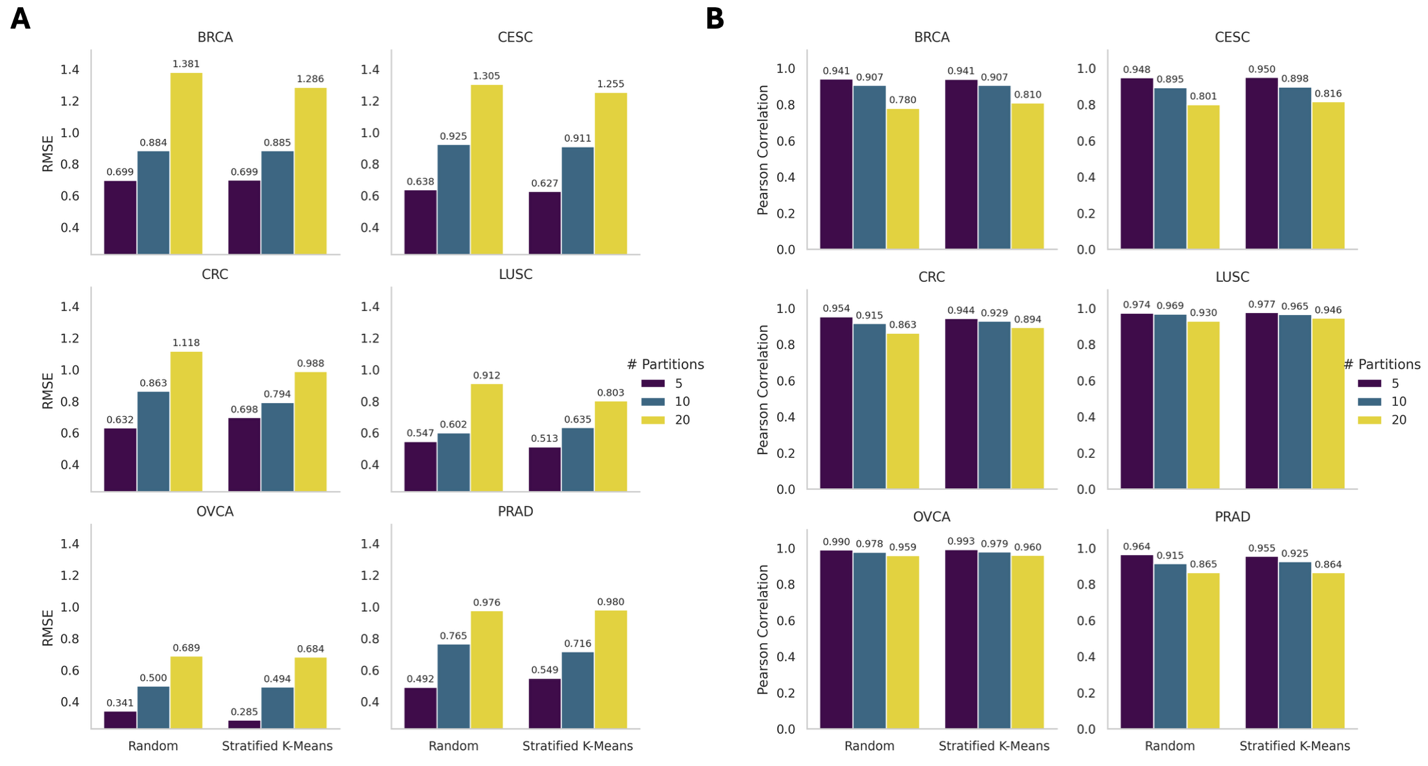

**Supplementary Figure S42. Sensitivity analysis results for real datasets.** **A.** Accuracies (left column) and AUC scores (right column) of linear classifiers fit to predict spot label with 10x Visium BRCA dataset gene set activity scores—computed by GESSO across six unique spatial smoothing strengths ($\beta$; see Supplementary Note 1)—as features. **B.** Same as A but for each of the three development stages of the Stereo-seq mouse embryo dataset.

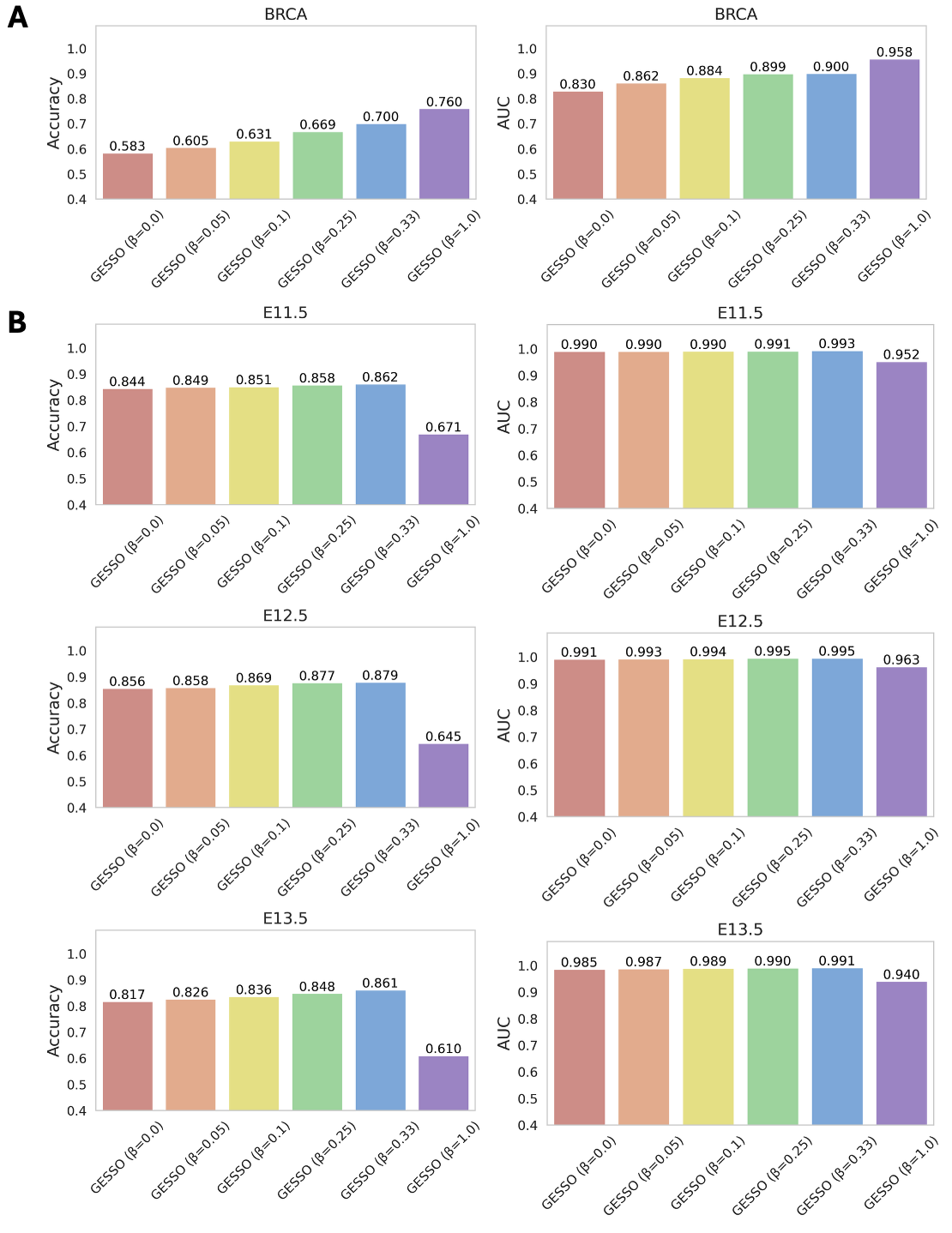

**Supplementary Figure S43. Sensitivity analysis results for simulated datasets.** Depicts the accuracy of GESSO-generated GASs in predicting the White Matter region of the simulation study dataset across a range of the spatial smoothing strength parameter $\beta$.

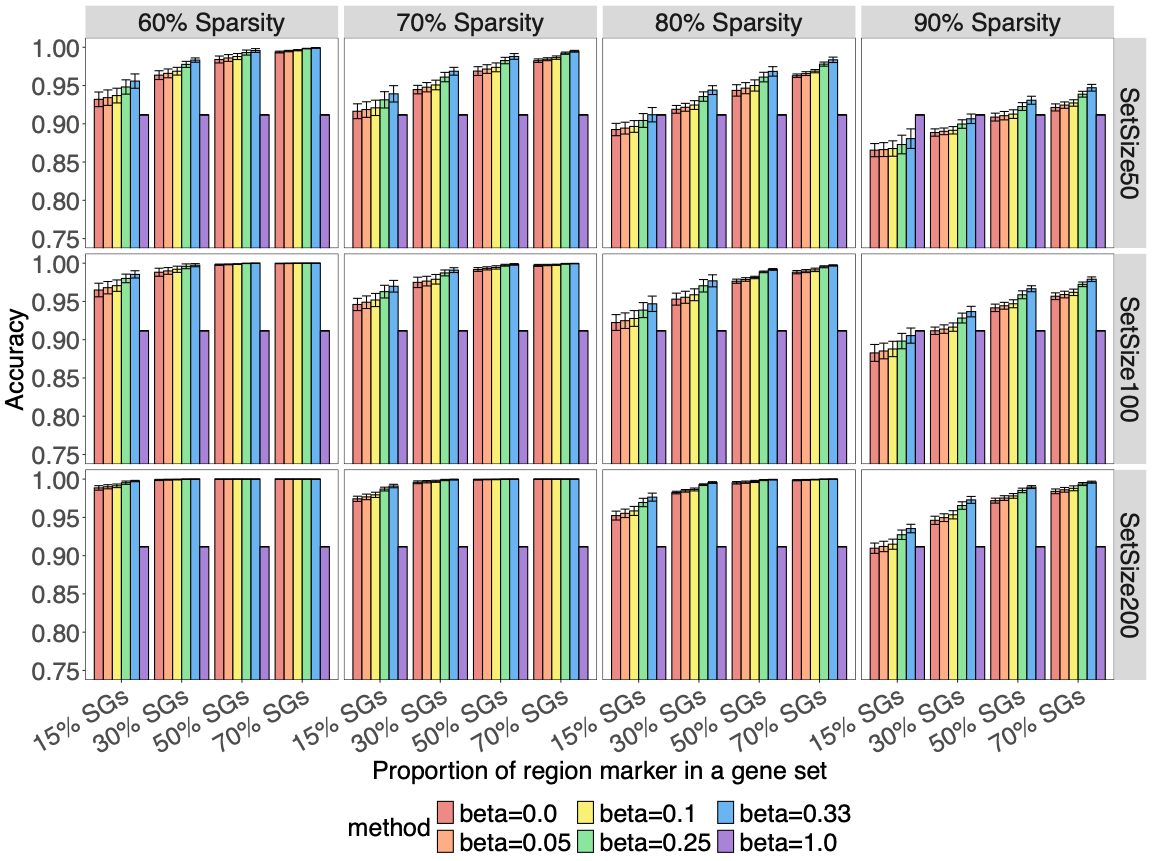

**Supplementary Figure S44.** **Sensitivity analysis results for simulated datasets.** Depicts the AUC of GESSO-generated GASs in predicting the White Matter region of the simulation study dataset across a range of the spatial smoothing strength parameter $\beta$.

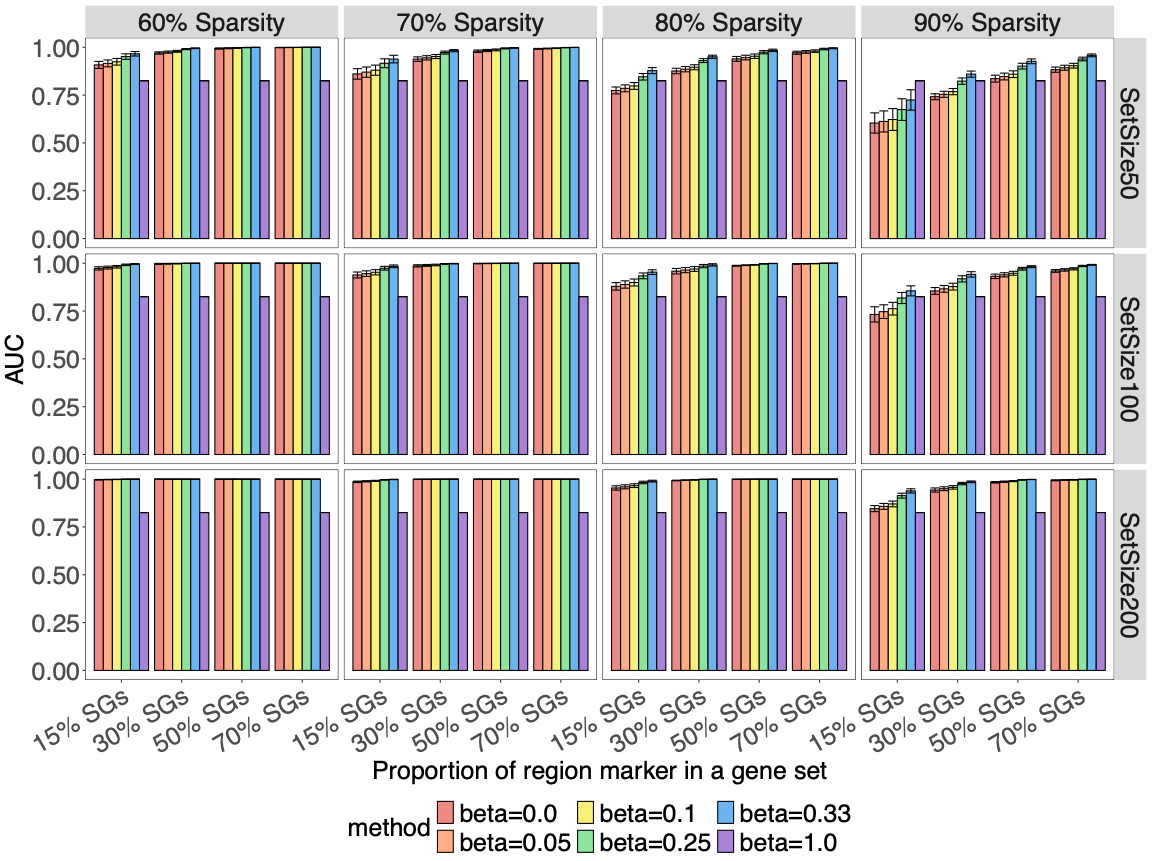

### Supplementary Tables

**Supplementary Table 1. List of all spatial transcriptomics data and scRNA-seq data used in in our analysis.** scRNA-seq data sets were used as the references for cell type decomposition. The table contains data set name (1^st^ column), tissue (2^nd^ column), experimental protocol/platform (3^rd^ column), number of spatial locations/cells (4^th^ column), number of genes (5^th^ column), download link (6^th^ column) and existence of H&E image and spatial labels (7^th^ column). GSEXXX represents the GEO accession number that the corresponding dataset can be downloaded from the NCBI GEO website.

| **Data Set** | **Tissue** | **Platform** | **# Locations** | **# Genes** | **URL to Dataset Source** | **Spot Label Source** |
| --- | --- | --- | --- | --- | --- | --- |
| Human_BRCA | Human Breast Cancer | 10x Visium | 2518 | 17943 | https://www.10xgenomics.com/datasets/human-breast-cancer-ductal-carcinoma-in-situ-invasive-carcinoma-ffpe-1-standard-1-3-0 | manual from annotated histology image |
| Human_CRC | Human Colorectal Cancer | 10x Visium | 2660 | 17943 | https://www.10xgenomics.com/datasets/human-intestine-cancer-1-standard | N/A |
| Human_LUSC | Human Lung Cancer | 10x Visium | 3858 | 18085 | https://www.10xgenomics.com/datasets/human-lung-cancer-ffpe-2-standard | N/A |
| Human_CESC | Human Cervical Cancer | 10x Visium | 3455 | 17943 | https://www.10xgenomics.com/datasets/human-ovarian-cancer-1-standard | N/A |
| Human_OVCA | Human Ovarian Canver | 10x Visium | 2781 | 17943 | https://www.10xgenomics.com/datasets/human-cervical-cancer-1-standard | N/A |
| Human_PRAD | Human Prostate Cancer | 10x Visium | 4371 | 17943 | https://www.10xgenomics.com/datasets/human-prostate-cancer-adenocarcinoma-with-invasive-carcinoma-ffpe-1-standard-1-3-0 | N/A |
| Mouse_Embryo_E115 | Mouse Embryo | Stereo-seq | 30124 | 23761 | https://db.cngb.org/stomics/mosta/download/ | included in original |
| Mouse_Embryo_E125 | Mouse Embryo | Stereo-seq | 51365 | 23761 | https://db.cngb.org/stomics/mosta/download/ | included in original |
| Mouse_Embryo_E135 | Mouse Embryo | Stereo-seq | 77369 | 23761 | https://db.cngb.org/stomics/mosta/download/ | included in original |
| Human_CRC_Tumor | Human Colorectal Cancer | Visium HD | 435773 | 18035 | https://www.10xgenomics.com/platforms/visium/product-family/dataset-human-crc | included in original |
| Human_CRC_Normal | Human normal adjacent tissue | Visium HD | 541968 | 18035 | https://www.10xgenomics.com/platforms/visium/product-family/dataset-human-crc | included in original |
| Human_Lymphnode | Human Lymph node | Xenium Prime | 708983 | 11094 | https://www.10xgenomics.com/datasets/preview-data-xenium-prime-gene-expression | manual from morphology and protein image |
| Human_NSCLS | Human Lung Cancer | CosMx | 57993 | 960 | https://zenodo.org/records/15240431 | included in original |
| Wu et al.  (GSE176078) | Human  Breast Cancer | 10x  Chromium | 100064 | 29733 | https://www.ncbi.nlm.nih.gov/geo/query/acc.cgi?acc=GSE176078 |  |
| Lee et al. (GSE144735) | Human Colorectal Cancer | 10x  Chromium | 27414 | 33694 | https://www.ncbi.nlm.nih.gov/geo/query/acc.cgi?acc=GSE144735 |  |
| Kim et al.  (GSE131907) | Human Lung Cancer | 10x  Chromium | 208506 | 29634 | https://www.ncbi.nlm.nih.gov/geo/query/acc.cgi?acc=GSE131907 |  |
| Zheng et al. | Human Ovarian Cancer | 10x  Chromium | 223363 | 27127 | <https://data.mendeley.com/datasets/rc47y6m9mp/1> |  |
| Hirz et al.  (GSE181294) | Human Prostate Cancer | 10x  Chromium | 156301 | 32629 | https://www.ncbi.nlm.nih.gov/geo/query/acc.cgi?acc=GSE181294 |  |

**Supplementary Table 2. The proportions of zero expressions within the SRT data analyzed in this study.** The 1^st^ column is the data set name, the 2^nd^ column is their platforms, and the 3^rd^ column are the percentage of zeros over the whole expression matrices. Genes with no expression overall all locations are removed.

| **Data Set** | **Platform** | **Sparsity (percentage of zero expressions for all expressed genes)** |
| --- | --- | --- |
| Human_BRCA | 10x Visium | 72.94% |
| Human_CRC | 10x Visium | 61.49% |
| Human_LUSC | 10x Visium | 66.89% |
| Human_CESC | 10x Visium | 65.38% |
| Human_OVCA | 10x Visium | 60.03% |
| Human_PRAD | 10x Visium | 71.99% |
| Mouse_Embryo_E115 | Stereo-seq | 88.22% |
| Mouse_Embryo_E125 | Stereo-seq | 89.79% |
| Mouse_Embryo_E135 | Stereo-seq | 91.28% |
| Human_CRC_Tumor | Visium HD | 98.28% |
| Human_CRC_Normal | Visium HD | 98.79% |
| Human_Lymphnode | Xenium Prime | 97.62% |
| Human_NSCLS | CosMx | 89.52% |

**Supplementary Table 3. Table summarizing the evaluation of prediction performance for all methods in the simulation study.** The first 3 columns describe the scenario for each simulation setting: the 1^st^ column is the number of genes in a set, the 2^nd^ column is the sparsity level of data, the 3^rd^ column is the proportions of markers in the gene set. Each scenario has 10 replicates. The rest columns are the evaluations of prediction, including mean AUC and its standard deviation, mean accuracy and its standard deviation, and mean F1 score and its stand deviation.

| geneset_size | sparsity | mix_prop | method | mean.auc | sd.auc | mean.acc | sd.acc | mean.f1 | sd.f1 |
| --- | --- | --- | --- | --- | --- | --- | --- | --- | --- |
| SetSize50 | 60% Sparsity | 15% Markers | AUCell | 0.51861605 | 0.01927321 | 0.85793409 | 0 | 0.92353555 | 0 |
| SetSize50 | 60% Sparsity | 15% Markers | GSVA | 0.54093693 | 0.01447612 | 0.85793409 | 0 | 0.92353555 | 0 |
| SetSize50 | 60% Sparsity | 15% Markers | ssGSEA | 0.52394471 | 0.01951834 | 0.85793409 | 0 | 0.92353555 | 0 |
| SetSize50 | 60% Sparsity | 15% Markers | GSDensity | 0.60956443 | 0.0864791 | 0.85793409 | 0 | 0.92353555 | 0 |
| SetSize50 | 60% Sparsity | 15% Markers | GESSO | 0.95160419 | 0.01432455 | 0.94821379 | 0.00918014 | 0.97032076 | 0.00516724 |
| SetSize50 | 60% Sparsity | 30% Markers | AUCell | 0.5537586 | 0.03943426 | 0.85798948 | 0.00011676 | 0.92356308 | 5.805E-05 |
| SetSize50 | 60% Sparsity | 30% Markers | GSVA | 0.60171953 | 0.01819617 | 0.85793409 | 0 | 0.92353555 | 0 |
| SetSize50 | 60% Sparsity | 30% Markers | ssGSEA | 0.57615509 | 0.02087904 | 0.85793409 | 0 | 0.92353555 | 0 |
| SetSize50 | 60% Sparsity | 30% Markers | GSDensity | 0.71960348 | 0.05777316 | 0.87560233 | 0.01688995 | 0.93213648 | 0.00840125 |
| SetSize50 | 60% Sparsity | 30% Markers | GESSO | 0.99034603 | 0.00281148 | 0.97762393 | 0.00391161 | 0.98703497 | 0.00225569 |
| SetSize50 | 60% Sparsity | 50% Markers | AUCell | 0.66622905 | 0.05131726 | 0.86089726 | 0.00436903 | 0.9247724 | 0.00187708 |
| SetSize50 | 60% Sparsity | 50% Markers | GSVA | 0.79100545 | 0.02322894 | 0.86568818 | 0.00521534 | 0.92649722 | 0.00240079 |
| SetSize50 | 60% Sparsity | 50% Markers | ssGSEA | 0.71261878 | 0.0411807 | 0.8607311 | 0.00353309 | 0.9245976 | 0.00150222 |
| SetSize50 | 60% Sparsity | 50% Markers | GSDensity | 0.86059144 | 0.03526499 | 0.91772362 | 0.01587443 | 0.95394004 | 0.00857321 |
| SetSize50 | 60% Sparsity | 50% Markers | GESSO | 0.99878706 | 0.00093672 | 0.99285516 | 0.00337104 | 0.99584347 | 0.00195784 |
| SetSize50 | 60% Sparsity | 70% Markers | AUCell | 0.78337376 | 0.04358507 | 0.87607311 | 0.00614299 | 0.93163708 | 0.00282499 |
| SetSize50 | 60% Sparsity | 70% Markers | GSVA | 0.92436182 | 0.01874927 | 0.91575741 | 0.00996253 | 0.95197092 | 0.00541043 |
| SetSize50 | 60% Sparsity | 70% Markers | ssGSEA | 0.85844266 | 0.01336411 | 0.89003046 | 0.00478138 | 0.93839166 | 0.00249976 |
| SetSize50 | 60% Sparsity | 70% Markers | GSDensity | 0.92974088 | 0.01533765 | 0.94450291 | 0.00418809 | 0.96853532 | 0.00232701 |
| SetSize50 | 60% Sparsity | 70% Markers | GESSO | 0.99984226 | 0.00018874 | 0.99831072 | 0.00035632 | 0.99901585 | 0.00020749 |
| SetSize50 | 70% Sparsity | 15% Markers | AUCell | 0.52849763 | 0.02364367 | 0.85793409 | 0 | 0.92353555 | 0 |
| SetSize50 | 70% Sparsity | 15% Markers | GSVA | 0.52689712 | 0.01061453 | 0.85793409 | 0 | 0.92353555 | 0 |
| SetSize50 | 70% Sparsity | 15% Markers | ssGSEA | 0.52042694 | 0.01193584 | 0.85793409 | 0 | 0.92353555 | 0 |
| SetSize50 | 70% Sparsity | 15% Markers | GSDensity | 0.59643353 | 0.10632245 | 0.85823872 | 0.00096331 | 0.92368494 | 0.00048048 |
| SetSize50 | 70% Sparsity | 15% Markers | GESSO | 0.91678427 | 0.02307459 | 0.93137635 | 0.01077925 | 0.96095858 | 0.00592461 |
| SetSize50 | 70% Sparsity | 30% Markers | AUCell | 0.55157226 | 0.01980632 | 0.85793409 | 0 | 0.92353555 | 0 |
| SetSize50 | 70% Sparsity | 30% Markers | GSVA | 0.56020334 | 0.01619421 | 0.85793409 | 0 | 0.92353555 | 0 |
| SetSize50 | 70% Sparsity | 30% Markers | ssGSEA | 0.54487584 | 0.01474235 | 0.85793409 | 0 | 0.92353555 | 0 |
| SetSize50 | 70% Sparsity | 30% Markers | GSDensity | 0.65124409 | 0.05503957 | 0.86865134 | 0.01114861 | 0.92879516 | 0.00553131 |
| SetSize50 | 70% Sparsity | 30% Markers | GESSO | 0.9724096 | 0.00770767 | 0.9611188 | 0.00628688 | 0.97759188 | 0.00357788 |
| SetSize50 | 70% Sparsity | 50% Markers | AUCell | 0.68213807 | 0.04012221 | 0.86269731 | 0.00573634 | 0.92560451 | 0.00272681 |
| SetSize50 | 70% Sparsity | 50% Markers | GSVA | 0.65691033 | 0.02920759 | 0.85834949 | 0.00113245 | 0.92371544 | 0.00050149 |
| SetSize50 | 70% Sparsity | 50% Markers | ssGSEA | 0.64212181 | 0.03005064 | 0.8579064 | 0.00024248 | 0.92347588 | 0.00022636 |
| SetSize50 | 70% Sparsity | 50% Markers | GSDensity | 0.81068173 | 0.02677216 | 0.90747715 | 0.01241256 | 0.94859273 | 0.0066561 |
| SetSize50 | 70% Sparsity | 50% Markers | GESSO | 0.99424926 | 0.00268213 | 0.98255331 | 0.00449961 | 0.98987419 | 0.0026035 |
| SetSize50 | 70% Sparsity | 70% Markers | AUCell | 0.79432606 | 0.0430523 | 0.87978399 | 0.01438532 | 0.93357638 | 0.00733804 |
| SetSize50 | 70% Sparsity | 70% Markers | GSVA | 0.691647 | 0.01998905 | 0.85804486 | 0.00080686 | 0.92350403 | 0.00041081 |
| SetSize50 | 70% Sparsity | 70% Markers | ssGSEA | 0.74420113 | 0.01592641 | 0.86363888 | 0.00272965 | 0.92578496 | 0.00135334 |
| SetSize50 | 70% Sparsity | 70% Markers | GSDensity | 0.90454981 | 0.01526205 | 0.93533647 | 0.00491288 | 0.96354916 | 0.00270144 |
| SetSize50 | 70% Sparsity | 70% Markers | GESSO | 0.99833521 | 0.00078859 | 0.99221822 | 0.00147956 | 0.99547284 | 0.00086054 |
| SetSize50 | 80% Sparsity | 15% Markers | AUCell | 0.52886 | 0.02240826 | 0.85793409 | 0 | 0.92353555 | 0 |
| SetSize50 | 80% Sparsity | 15% Markers | GSVA | 0.51531536 | 0.00922263 | 0.85793409 | 0 | 0.92353555 | 0 |
| SetSize50 | 80% Sparsity | 15% Markers | ssGSEA | 0.51174813 | 0.0089018 | 0.85793409 | 0 | 0.92353555 | 0 |
| SetSize50 | 80% Sparsity | 15% Markers | GSDensity | 0.58983706 | 0.08109937 | 0.85793409 | 0 | 0.92353555 | 0 |
| SetSize50 | 80% Sparsity | 15% Markers | GESSO | 0.84638634 | 0.01702704 | 0.90440321 | 0.00885777 | 0.94635906 | 0.0047851 |
| SetSize50 | 80% Sparsity | 30% Markers | AUCell | 0.54943103 | 0.03958487 | 0.8579064 | 8.7573E-05 | 0.9235195 | 5.0747E-05 |
| SetSize50 | 80% Sparsity | 30% Markers | GSVA | 0.52516508 | 0.0131713 | 0.85793409 | 0 | 0.92353555 | 0 |
| SetSize50 | 80% Sparsity | 30% Markers | ssGSEA | 0.52908322 | 0.01005676 | 0.85793409 | 0 | 0.92353555 | 0 |
| SetSize50 | 80% Sparsity | 30% Markers | GSDensity | 0.63248175 | 0.06572462 | 0.86671282 | 0.00994219 | 0.92784128 | 0.00488662 |
| SetSize50 | 80% Sparsity | 30% Markers | GESSO | 0.93071642 | 0.01047124 | 0.93575187 | 0.00581337 | 0.96333231 | 0.00324192 |
| SetSize50 | 80% Sparsity | 50% Markers | AUCell | 0.66886154 | 0.05633261 | 0.85995569 | 0.00271352 | 0.92421293 | 0.00116886 |
| SetSize50 | 80% Sparsity | 50% Markers | GSVA | 0.55259792 | 0.02478478 | 0.85793409 | 0 | 0.92353555 | 0 |
| SetSize50 | 80% Sparsity | 50% Markers | ssGSEA | 0.57233322 | 0.02739052 | 0.85793409 | 0 | 0.92353555 | 0 |
| SetSize50 | 80% Sparsity | 50% Markers | GSDensity | 0.80798497 | 0.04330873 | 0.90636943 | 0.01639205 | 0.94803283 | 0.00861838 |
| SetSize50 | 80% Sparsity | 50% Markers | GESSO | 0.97446557 | 0.00829776 | 0.9611465 | 0.00612465 | 0.97759489 | 0.00347994 |
| SetSize50 | 80% Sparsity | 70% Markers | AUCell | 0.80546249 | 0.03865455 | 0.88188867 | 0.01216557 | 0.93458632 | 0.00621299 |
| SetSize50 | 80% Sparsity | 70% Markers | GSVA | 0.64168155 | 0.05247174 | 0.85785101 | 0.00037039 | 0.9234691 | 0.00021858 |
| SetSize50 | 80% Sparsity | 70% Markers | ssGSEA | 0.65345283 | 0.01883005 | 0.85810025 | 0.0006151 | 0.92356797 | 0.00034026 |
| SetSize50 | 80% Sparsity | 70% Markers | GSDensity | 0.90724557 | 0.01805289 | 0.93666574 | 0.00555867 | 0.96424724 | 0.00303613 |
| SetSize50 | 80% Sparsity | 70% Markers | GESSO | 0.99097374 | 0.00381077 | 0.97809471 | 0.00262866 | 0.98730181 | 0.00150723 |
| SetSize50 | 90% Sparsity | 15% Markers | AUCell | 0.52993697 | 0.02398634 | 0.85793409 | 0 | 0.92353555 | 0 |
| SetSize50 | 90% Sparsity | 15% Markers | GSVA | 0.5116036 | 0.0085492 | 0.85793409 | 0 | 0.92353555 | 0 |
| SetSize50 | 90% Sparsity | 15% Markers | ssGSEA | 0.5138019 | 0.0077224 | 0.85793409 | 0 | 0.92353555 | 0 |
| SetSize50 | 90% Sparsity | 15% Markers | GSDensity | 0.60448639 | 0.09787931 | 0.85793409 | 0 | 0.92353555 | 0 |
| SetSize50 | 90% Sparsity | 15% Markers | GESSO | 0.67626596 | 0.05405606 | 0.87341457 | 0.01200495 | 0.9310346 | 0.00586971 |
| SetSize50 | 90% Sparsity | 30% Markers | AUCell | 0.55890712 | 0.0227273 | 0.85793409 | 0 | 0.92353555 | 0 |
| SetSize50 | 90% Sparsity | 30% Markers | GSVA | 0.51276942 | 0.0078798 | 0.85793409 | 0 | 0.92353555 | 0 |
| SetSize50 | 90% Sparsity | 30% Markers | ssGSEA | 0.51679528 | 0.01061338 | 0.85793409 | 0 | 0.92353555 | 0 |
| SetSize50 | 90% Sparsity | 30% Markers | GSDensity | 0.67391155 | 0.04728835 | 0.86754362 | 0.00730368 | 0.92810371 | 0.00352994 |
| SetSize50 | 90% Sparsity | 30% Markers | GESSO | 0.8237392 | 0.01648058 | 0.89972307 | 0.00573731 | 0.94398742 | 0.0030004 |
| SetSize50 | 90% Sparsity | 50% Markers | AUCell | 0.68840559 | 0.03593629 | 0.86217114 | 0.0045468 | 0.92529314 | 0.00204842 |
| SetSize50 | 90% Sparsity | 50% Markers | GSVA | 0.56625673 | 0.02786755 | 0.85793409 | 0 | 0.92353555 | 0 |
| SetSize50 | 90% Sparsity | 50% Markers | ssGSEA | 0.54008849 | 0.01674926 | 0.85793409 | 0 | 0.92353555 | 0 |
| SetSize50 | 90% Sparsity | 50% Markers | GSDensity | 0.82387298 | 0.03372421 | 0.90584326 | 0.00880973 | 0.94748672 | 0.00462037 |
| SetSize50 | 90% Sparsity | 50% Markers | GESSO | 0.90206176 | 0.01498888 | 0.92298532 | 0.00498896 | 0.95632333 | 0.00271253 |
| SetSize50 | 90% Sparsity | 70% Markers | AUCell | 0.77609434 | 0.02147656 | 0.87499308 | 0.00465816 | 0.93110201 | 0.00231252 |
| SetSize50 | 90% Sparsity | 70% Markers | GSVA | 0.60193497 | 0.04139023 | 0.85793409 | 0 | 0.92353555 | 0 |
| SetSize50 | 90% Sparsity | 70% Markers | ssGSEA | 0.56421542 | 0.00786776 | 0.85793409 | 0 | 0.92353555 | 0 |
| SetSize50 | 90% Sparsity | 70% Markers | GSDensity | 0.89760067 | 0.01612188 | 0.93342564 | 0.00462143 | 0.96235019 | 0.00252065 |
| SetSize50 | 90% Sparsity | 70% Markers | GESSO | 0.93915505 | 0.00961522 | 0.93896428 | 0.00343465 | 0.96511407 | 0.00192585 |
| SetSize100 | 60% Sparsity | 15% Markers | AUCell | 0.52920705 | 0.02512072 | 0.85793409 | 0 | 0.92353555 | 0 |
| SetSize100 | 60% Sparsity | 15% Markers | GSVA | 0.5627187 | 0.0165552 | 0.85793409 | 0 | 0.92353555 | 0 |
| SetSize100 | 60% Sparsity | 15% Markers | ssGSEA | 0.52485827 | 0.01851532 | 0.85793409 | 0 | 0.92353555 | 0 |
| SetSize100 | 60% Sparsity | 15% Markers | GSDensity | 0.59071696 | 0.05194241 | 0.85840487 | 0.00148875 | 0.92376573 | 0.00072788 |
| SetSize100 | 60% Sparsity | 15% Markers | GESSO | 0.99164121 | 0.00391919 | 0.97997785 | 0.00566647 | 0.98839203 | 0.0032705 |
| SetSize100 | 60% Sparsity | 30% Markers | AUCell | 0.57575453 | 0.03890195 | 0.85810025 | 0.00052544 | 0.9236137 | 0.00024714 |
| SetSize100 | 60% Sparsity | 30% Markers | GSVA | 0.63199272 | 0.0189623 | 0.8579064 | 8.7573E-05 | 0.9235195 | 5.0747E-05 |
| SetSize100 | 60% Sparsity | 30% Markers | ssGSEA | 0.59674644 | 0.03455383 | 0.85796178 | 8.7573E-05 | 0.92354932 | 4.3537E-05 |
| SetSize100 | 60% Sparsity | 30% Markers | GSDensity | 0.71185359 | 0.05946317 | 0.88532263 | 0.01267545 | 0.93718773 | 0.00641483 |
| SetSize100 | 60% Sparsity | 30% Markers | GESSO | 0.99943182 | 0.00055907 | 0.99601218 | 0.00288151 | 0.99767868 | 0.00167559 |
| SetSize100 | 60% Sparsity | 50% Markers | AUCell | 0.7204013 | 0.04767582 | 0.86399889 | 0.00520299 | 0.92603517 | 0.0023822 |
| SetSize100 | 60% Sparsity | 50% Markers | GSVA | 0.86022366 | 0.01882149 | 0.88211022 | 0.00642212 | 0.93405878 | 0.0033028 |
| SetSize100 | 60% Sparsity | 50% Markers | ssGSEA | 0.78389598 | 0.0228555 | 0.86779286 | 0.00418402 | 0.92749966 | 0.00201604 |
| SetSize100 | 60% Sparsity | 50% Markers | GSDensity | 0.85096264 | 0.01786099 | 0.93151482 | 0.00699985 | 0.96152411 | 0.00379858 |
| SetSize100 | 60% Sparsity | 50% Markers | GESSO | 0.9999956 | 6.9119E-06 | 0.99969538 | 0.00024248 | 0.99982248 | 0.0001413 |
| SetSize100 | 60% Sparsity | 70% Markers | AUCell | 0.87133342 | 0.05213029 | 0.90193852 | 0.01789846 | 0.94496839 | 0.00930116 |
| SetSize100 | 60% Sparsity | 70% Markers | GSVA | 0.97111392 | 0.01401941 | 0.95117696 | 0.01455761 | 0.97184631 | 0.00828857 |
| SetSize100 | 60% Sparsity | 70% Markers | ssGSEA | 0.9269272 | 0.01788618 | 0.91872058 | 0.00962091 | 0.95366684 | 0.00524179 |
| SetSize100 | 60% Sparsity | 70% Markers | GSDensity | 0.91000941 | 0.0166014 | 0.95361396 | 0.00437527 | 0.97363247 | 0.00242301 |
| SetSize100 | 60% Sparsity | 70% Markers | GESSO | 0.99999994 | 1.9898E-07 | 0.99997231 | 8.7573E-05 | 0.99998386 | 5.1029E-05 |
| SetSize100 | 70% Sparsity | 15% Markers | AUCell | 0.53323527 | 0.02106413 | 0.85793409 | 0 | 0.92353555 | 0 |
| SetSize100 | 70% Sparsity | 15% Markers | GSVA | 0.54000839 | 0.01501907 | 0.85793409 | 0 | 0.92353555 | 0 |
| SetSize100 | 70% Sparsity | 15% Markers | ssGSEA | 0.51972083 | 0.01383239 | 0.85793409 | 0 | 0.92353555 | 0 |
| SetSize100 | 70% Sparsity | 15% Markers | GSDensity | 0.59155558 | 0.08693243 | 0.85793409 | 0 | 0.92353555 | 0 |
| SetSize100 | 70% Sparsity | 15% Markers | GESSO | 0.97398819 | 0.00974436 | 0.96283578 | 0.00808632 | 0.97858785 | 0.00459941 |
| SetSize100 | 70% Sparsity | 30% Markers | AUCell | 0.58727815 | 0.0370435 | 0.8579064 | 0.00020434 | 0.92351722 | 0.00011481 |
| SetSize100 | 70% Sparsity | 30% Markers | GSVA | 0.57560112 | 0.01822823 | 0.85793409 | 0 | 0.92353555 | 0 |
| SetSize100 | 70% Sparsity | 30% Markers | ssGSEA | 0.55829857 | 0.02178213 | 0.85793409 | 0 | 0.92353555 | 0 |
| SetSize100 | 70% Sparsity | 30% Markers | GSDensity | 0.6693231 | 0.05930401 | 0.88321795 | 0.01233999 | 0.93617052 | 0.00619765 |
| SetSize100 | 70% Sparsity | 30% Markers | GESSO | 0.99601686 | 0.00204163 | 0.98731653 | 0.00385677 | 0.99262853 | 0.00222987 |
| SetSize100 | 70% Sparsity | 50% Markers | AUCell | 0.72708633 | 0.03684742 | 0.86311271 | 0.0029656 | 0.92552281 | 0.00125893 |
| SetSize100 | 70% Sparsity | 50% Markers | GSVA | 0.68977974 | 0.02278349 | 0.85884796 | 0.00152829 | 0.92390231 | 0.00072929 |
| SetSize100 | 70% Sparsity | 50% Markers | ssGSEA | 0.69793308 | 0.01346779 | 0.85995569 | 0.00186937 | 0.92437083 | 0.00093039 |
| SetSize100 | 70% Sparsity | 50% Markers | GSDensity | 0.80118891 | 0.01764065 | 0.92251454 | 0.00602894 | 0.95672225 | 0.00325361 |
| SetSize100 | 70% Sparsity | 50% Markers | GESSO | 0.99979563 | 0.00027648 | 0.99742454 | 0.00114592 | 0.99849974 | 0.00066716 |
| SetSize100 | 70% Sparsity | 70% Markers | AUCell | 0.85659937 | 0.02447274 | 0.89349211 | 0.00880799 | 0.94029985 | 0.00462902 |
| SetSize100 | 70% Sparsity | 70% Markers | GSVA | 0.71077995 | 0.02775596 | 0.85840487 | 0.00097736 | 0.92355212 | 0.00062099 |
| SetSize100 | 70% Sparsity | 70% Markers | ssGSEA | 0.81877625 | 0.01619217 | 0.87590695 | 0.00531159 | 0.93129988 | 0.00267748 |
| SetSize100 | 70% Sparsity | 70% Markers | GSDensity | 0.87885299 | 0.01239117 | 0.94093049 | 0.00494538 | 0.9666583 | 0.0027105 |
| SetSize100 | 70% Sparsity | 70% Markers | GESSO | 0.99996596 | 5.9717E-05 | 0.9991969 | 0.00046063 | 0.99953199 | 0.00026839 |
| SetSize100 | 80% Sparsity | 15% Markers | AUCell | 0.53945956 | 0.02828054 | 0.85793409 | 0 | 0.92353555 | 0 |
| SetSize100 | 80% Sparsity | 15% Markers | GSVA | 0.52232447 | 0.0094656 | 0.85793409 | 0 | 0.92353555 | 0 |
| SetSize100 | 80% Sparsity | 15% Markers | ssGSEA | 0.51451776 | 0.00936902 | 0.85793409 | 0 | 0.92353555 | 0 |
| SetSize100 | 80% Sparsity | 15% Markers | GSDensity | 0.60113385 | 0.08523474 | 0.85793409 | 0 | 0.92353555 | 0 |
| SetSize100 | 80% Sparsity | 15% Markers | GESSO | 0.93416975 | 0.01506159 | 0.93868734 | 0.00978596 | 0.96501351 | 0.00547342 |
| SetSize100 | 80% Sparsity | 30% Markers | AUCell | 0.58360544 | 0.03567093 | 0.85796178 | 0.0001572 | 0.92354018 | 0.00010009 |
| SetSize100 | 80% Sparsity | 30% Markers | GSVA | 0.53800773 | 0.01025329 | 0.85793409 | 0 | 0.92353555 | 0 |
| SetSize100 | 80% Sparsity | 30% Markers | ssGSEA | 0.54763276 | 0.0160125 | 0.85793409 | 0 | 0.92353555 | 0 |
| SetSize100 | 80% Sparsity | 30% Markers | GSDensity | 0.66240032 | 0.0489277 | 0.87939629 | 0.01144371 | 0.93427179 | 0.00575549 |
| SetSize100 | 80% Sparsity | 30% Markers | GESSO | 0.98488605 | 0.00845323 | 0.97056217 | 0.0084534 | 0.98298295 | 0.00481223 |
| SetSize100 | 80% Sparsity | 50% Markers | AUCell | 0.74436576 | 0.03883367 | 0.86743284 | 0.00561814 | 0.92757164 | 0.00266735 |
| SetSize100 | 80% Sparsity | 50% Markers | GSVA | 0.57253671 | 0.02004234 | 0.85793409 | 0 | 0.92353555 | 0 |
| SetSize100 | 80% Sparsity | 50% Markers | ssGSEA | 0.60586922 | 0.01424394 | 0.85793409 | 0 | 0.92353555 | 0 |
| SetSize100 | 80% Sparsity | 50% Markers | GSDensity | 0.79828481 | 0.02207701 | 0.91941291 | 0.00690789 | 0.95503774 | 0.00369917 |
| SetSize100 | 80% Sparsity | 50% Markers | GESSO | 0.99691022 | 0.00077075 | 0.9886458 | 0.00131846 | 0.99339924 | 0.0007643 |
| SetSize100 | 80% Sparsity | 70% Markers | AUCell | 0.8756093 | 0.03569757 | 0.90182775 | 0.01517954 | 0.94477461 | 0.00804014 |
| SetSize100 | 80% Sparsity | 70% Markers | GSVA | 0.66162487 | 0.03542301 | 0.85782332 | 0.00060108 | 0.92343934 | 0.00033864 |
| SetSize100 | 80% Sparsity | 70% Markers | ssGSEA | 0.69191392 | 0.01376346 | 0.85846026 | 0.00119968 | 0.92359877 | 0.0006633 |
| SetSize100 | 80% Sparsity | 70% Markers | GSDensity | 0.88329262 | 0.01410619 | 0.94275824 | 0.00442715 | 0.96762012 | 0.00241907 |
| SetSize100 | 80% Sparsity | 70% Markers | GESSO | 0.99952437 | 0.00024332 | 0.99534755 | 0.00125759 | 0.99729092 | 0.00073163 |
| SetSize100 | 90% Sparsity | 15% Markers | AUCell | 0.52863484 | 0.02846263 | 0.85793409 | 0 | 0.92353555 | 0 |
| SetSize100 | 90% Sparsity | 15% Markers | GSVA | 0.50870234 | 0.00786813 | 0.85793409 | 0 | 0.92353555 | 0 |
| SetSize100 | 90% Sparsity | 15% Markers | ssGSEA | 0.50848557 | 0.01013186 | 0.85793409 | 0 | 0.92353555 | 0 |
| SetSize100 | 90% Sparsity | 15% Markers | GSDensity | 0.6011375 | 0.07427579 | 0.85793409 | 0 | 0.92353555 | 0 |
| SetSize100 | 90% Sparsity | 15% Markers | GESSO | 0.81819051 | 0.02930829 | 0.89819994 | 0.01034554 | 0.94322636 | 0.00542426 |
| SetSize100 | 90% Sparsity | 30% Markers | AUCell | 0.57029295 | 0.02983767 | 0.85793409 | 0 | 0.92353555 | 0 |
| SetSize100 | 90% Sparsity | 30% Markers | GSVA | 0.51021724 | 0.00722175 | 0.85793409 | 0 | 0.92353555 | 0 |
| SetSize100 | 90% Sparsity | 30% Markers | ssGSEA | 0.51902642 | 0.00838132 | 0.85793409 | 0 | 0.92353555 | 0 |
| SetSize100 | 90% Sparsity | 30% Markers | GSDensity | 0.65125944 | 0.04396299 | 0.87302686 | 0.00843341 | 0.93103129 | 0.00418169 |
| SetSize100 | 90% Sparsity | 30% Markers | GESSO | 0.91838267 | 0.0161645 | 0.92880089 | 0.00658123 | 0.9595021 | 0.00359757 |
| SetSize100 | 90% Sparsity | 50% Markers | AUCell | 0.73929807 | 0.0186904 | 0.86386043 | 0.00324061 | 0.92577754 | 0.00155642 |
| SetSize100 | 90% Sparsity | 50% Markers | GSVA | 0.54036157 | 0.01779562 | 0.85793409 | 0 | 0.92353555 | 0 |
| SetSize100 | 90% Sparsity | 50% Markers | ssGSEA | 0.55629338 | 0.00797801 | 0.85793409 | 0 | 0.92353555 | 0 |
| SetSize100 | 90% Sparsity | 50% Markers | GSDensity | 0.83629613 | 0.02985346 | 0.92846857 | 0.00623906 | 0.95981236 | 0.00333499 |
| SetSize100 | 90% Sparsity | 50% Markers | GESSO | 0.97114525 | 0.0070807 | 0.95865411 | 0.00484441 | 0.97618715 | 0.00274726 |
| SetSize100 | 90% Sparsity | 70% Markers | AUCell | 0.86228284 | 0.03378316 | 0.89346441 | 0.01249644 | 0.94024494 | 0.00656183 |
| SetSize100 | 90% Sparsity | 70% Markers | GSVA | 0.62654772 | 0.05504655 | 0.8579064 | 8.7573E-05 | 0.92351722 | 5.7973E-05 |
| SetSize100 | 90% Sparsity | 70% Markers | ssGSEA | 0.57996651 | 0.02200376 | 0.85793409 | 0 | 0.92353555 | 0 |
| SetSize100 | 90% Sparsity | 70% Markers | GSDensity | 0.8994259 | 0.02309378 | 0.94264747 | 0.00758908 | 0.96745229 | 0.0042029 |
| SetSize100 | 90% Sparsity | 70% Markers | GESSO | 0.98583057 | 0.00343493 | 0.97250069 | 0.00313326 | 0.98408775 | 0.00179865 |
| SetSize200 | 60% Sparsity | 15% Markers | AUCell | 0.55579749 | 0.02958108 | 0.85793409 | 0 | 0.92353555 | 0 |
| SetSize200 | 60% Sparsity | 15% Markers | GSVA | 0.60296104 | 0.01692986 | 0.85793409 | 0 | 0.92353555 | 0 |
| SetSize200 | 60% Sparsity | 15% Markers | ssGSEA | 0.54518233 | 0.02094914 | 0.85793409 | 0 | 0.92353555 | 0 |
| SetSize200 | 60% Sparsity | 15% Markers | GSDensity | 0.56426702 | 0.03303155 | 0.85793409 | 0 | 0.92353555 | 0 |
| SetSize200 | 60% Sparsity | 15% Markers | GESSO | 0.99962473 | 0.00033648 | 0.99526447 | 0.00217057 | 0.9972426 | 0.00126308 |
| SetSize200 | 60% Sparsity | 30% Markers | AUCell | 0.59977546 | 0.02790126 | 0.85796178 | 8.7573E-05 | 0.92354248 | 2.1929E-05 |
| SetSize200 | 60% Sparsity | 30% Markers | GSVA | 0.67904981 | 0.01409716 | 0.8578787 | 0.00025448 | 0.92345097 | 0.00012893 |
| SetSize200 | 60% Sparsity | 30% Markers | ssGSEA | 0.63220238 | 0.02426675 | 0.85793409 | 0.00013055 | 0.92352416 | 6.1448E-05 |
| SetSize200 | 60% Sparsity | 30% Markers | GSDensity | 0.71221979 | 0.03117354 | 0.89133204 | 0.01051789 | 0.94038198 | 0.00541469 |
| SetSize200 | 60% Sparsity | 30% Markers | GESSO | 0.99999975 | 5.306E-07 | 0.99991692 | 0.00018691 | 0.99995158 | 0.00010893 |
| SetSize200 | 60% Sparsity | 50% Markers | AUCell | 0.77856175 | 0.03728813 | 0.87183606 | 0.00596929 | 0.92956273 | 0.00279249 |
| SetSize200 | 60% Sparsity | 50% Markers | GSVA | 0.93425482 | 0.0139864 | 0.91808363 | 0.01231977 | 0.95320542 | 0.00683969 |
| SetSize200 | 60% Sparsity | 50% Markers | ssGSEA | 0.86211987 | 0.02157355 | 0.88756577 | 0.00832434 | 0.93707959 | 0.00434376 |
| SetSize200 | 60% Sparsity | 50% Markers | GSDensity | 0.83230903 | 0.00747153 | 0.93710883 | 0.00398211 | 0.96462026 | 0.00217484 |
| SetSize200 | 60% Sparsity | 50% Markers | GESSO | 1 | 0 | 1 | 0 | 1 | 0 |
| SetSize200 | 60% Sparsity | 70% Markers | AUCell | 0.92971269 | 0.02371261 | 0.92439767 | 0.0115915 | 0.95694737 | 0.00637268 |
| SetSize200 | 60% Sparsity | 70% Markers | GSVA | 0.9952326 | 0.0017503 | 0.9793686 | 0.00471823 | 0.98800796 | 0.00273629 |
| SetSize200 | 60% Sparsity | 70% Markers | ssGSEA | 0.97745669 | 0.00464285 | 0.9554694 | 0.00634973 | 0.97425603 | 0.0036449 |
| SetSize200 | 60% Sparsity | 70% Markers | GSDensity | 0.90122924 | 0.01316857 | 0.95580172 | 0.00249478 | 0.97487188 | 0.00138194 |
| SetSize200 | 60% Sparsity | 70% Markers | GESSO | 1 | 0 | 1 | 0 | 1 | 0 |
| SetSize200 | 70% Sparsity | 15% Markers | AUCell | 0.54764484 | 0.0222016 | 0.85793409 | 0 | 0.92353555 | 0 |
| SetSize200 | 70% Sparsity | 15% Markers | GSVA | 0.56168936 | 0.01988896 | 0.85793409 | 0 | 0.92353555 | 0 |
| SetSize200 | 70% Sparsity | 15% Markers | ssGSEA | 0.52996909 | 0.01930657 | 0.85793409 | 0 | 0.92353555 | 0 |
| SetSize200 | 70% Sparsity | 15% Markers | GSDensity | 0.59451051 | 0.05904757 | 0.85793409 | 0 | 0.92353555 | 0 |
| SetSize200 | 70% Sparsity | 15% Markers | GESSO | 0.99655151 | 0.0015044 | 0.98720576 | 0.0024797 | 0.99256523 | 0.00143825 |
| SetSize200 | 70% Sparsity | 30% Markers | AUCell | 0.61088758 | 0.02609205 | 0.85807256 | 0.00043787 | 0.92359074 | 0.00022474 |
| SetSize200 | 70% Sparsity | 30% Markers | GSVA | 0.61670442 | 0.01749585 | 0.85793409 | 0 | 0.92353555 | 0 |
| SetSize200 | 70% Sparsity | 30% Markers | ssGSEA | 0.60319083 | 0.01820347 | 0.85793409 | 0 | 0.92353555 | 0 |
| SetSize200 | 70% Sparsity | 30% Markers | GSDensity | 0.66637779 | 0.0250261 | 0.8866242 | 0.00893383 | 0.93800556 | 0.00457046 |
| SetSize200 | 70% Sparsity | 30% Markers | GESSO | 0.99989681 | 0.00022926 | 0.99889227 | 0.00079409 | 0.99935464 | 0.00046261 |
| SetSize200 | 70% Sparsity | 50% Markers | AUCell | 0.79013644 | 0.02517054 | 0.87283301 | 0.00493633 | 0.92996657 | 0.00233198 |
| SetSize200 | 70% Sparsity | 50% Markers | GSVA | 0.69876799 | 0.02571537 | 0.85801717 | 0.00089546 | 0.92343959 | 0.00050057 |
| SetSize200 | 70% Sparsity | 50% Markers | ssGSEA | 0.75370937 | 0.01842303 | 0.86078649 | 0.00194094 | 0.92422729 | 0.00099234 |
| SetSize200 | 70% Sparsity | 50% Markers | GSDensity | 0.78620855 | 0.01323321 | 0.92747161 | 0.00313435 | 0.9594351 | 0.00168154 |
| SetSize200 | 70% Sparsity | 50% Markers | GESSO | 0.99999994 | 1.9898E-07 | 0.99994461 | 0.00017515 | 0.99996772 | 0.00010207 |
| SetSize200 | 70% Sparsity | 70% Markers | AUCell | 0.94131094 | 0.01608288 | 0.92927167 | 0.01087636 | 0.95957447 | 0.00605355 |
| SetSize200 | 70% Sparsity | 70% Markers | GSVA | 0.75850904 | 0.03071365 | 0.85937413 | 0.00127776 | 0.92349102 | 0.00067542 |
| SetSize200 | 70% Sparsity | 70% Markers | ssGSEA | 0.90243684 | 0.01294991 | 0.90504016 | 0.00542587 | 0.94621187 | 0.00294627 |
| SetSize200 | 70% Sparsity | 70% Markers | GSDensity | 0.87093037 | 0.01246747 | 0.94699529 | 0.00286668 | 0.97002124 | 0.00158873 |
| SetSize200 | 70% Sparsity | 70% Markers | GESSO | 1 | 0 | 1 | 0 | 1 | 0 |
| SetSize200 | 80% Sparsity | 15% Markers | AUCell | 0.55322421 | 0.03201037 | 0.85793409 | 0 | 0.92353555 | 0 |
| SetSize200 | 80% Sparsity | 15% Markers | GSVA | 0.51940194 | 0.01579495 | 0.85793409 | 0 | 0.92353555 | 0 |
| SetSize200 | 80% Sparsity | 15% Markers | ssGSEA | 0.51196364 | 0.00812417 | 0.85793409 | 0 | 0.92353555 | 0 |
| SetSize200 | 80% Sparsity | 15% Markers | GSDensity | 0.61537274 | 0.07679663 | 0.85793409 | 0 | 0.92353555 | 0 |
| SetSize200 | 80% Sparsity | 15% Markers | GESSO | 0.98263389 | 0.00665939 | 0.96931598 | 0.00558582 | 0.98226507 | 0.00319912 |
| SetSize200 | 80% Sparsity | 30% Markers | AUCell | 0.59954382 | 0.04152672 | 0.8579064 | 0.00020434 | 0.92349888 | 0.00016045 |
| SetSize200 | 80% Sparsity | 30% Markers | GSVA | 0.52655439 | 0.01472716 | 0.85793409 | 0 | 0.92353555 | 0 |
| SetSize200 | 80% Sparsity | 30% Markers | ssGSEA | 0.54769876 | 0.01087104 | 0.85793409 | 0 | 0.92353555 | 0 |
| SetSize200 | 80% Sparsity | 30% Markers | GSDensity | 0.65357169 | 0.0420494 | 0.88127942 | 0.01118905 | 0.93529406 | 0.00567637 |
| SetSize200 | 80% Sparsity | 30% Markers | GESSO | 0.99888333 | 0.00031248 | 0.99285516 | 0.00100952 | 0.99584309 | 0.00058672 |
| SetSize200 | 80% Sparsity | 50% Markers | AUCell | 0.8031845 | 0.02862184 | 0.87623927 | 0.00647813 | 0.93160963 | 0.00314132 |
| SetSize200 | 80% Sparsity | 50% Markers | GSVA | 0.59940501 | 0.01568459 | 0.85793409 | 0 | 0.92353555 | 0 |
| SetSize200 | 80% Sparsity | 50% Markers | ssGSEA | 0.65170109 | 0.01456066 | 0.85782332 | 0.00049194 | 0.92344852 | 0.00027098 |
| SetSize200 | 80% Sparsity | 50% Markers | GSDensity | 0.78813609 | 0.01428177 | 0.92639158 | 0.00441511 | 0.95884523 | 0.00237744 |
| SetSize200 | 80% Sparsity | 50% Markers | GESSO | 0.9999698 | 2.2274E-05 | 0.99864304 | 0.00065858 | 0.99920939 | 0.00038369 |
| SetSize200 | 80% Sparsity | 70% Markers | AUCell | 0.94427663 | 0.01963077 | 0.93137635 | 0.01241486 | 0.96078408 | 0.00689565 |
| SetSize200 | 80% Sparsity | 70% Markers | GSVA | 0.68854452 | 0.0311977 | 0.85757408 | 0.00045317 | 0.92320972 | 0.00034775 |
| SetSize200 | 80% Sparsity | 70% Markers | ssGSEA | 0.76534984 | 0.01681601 | 0.86322348 | 0.00126199 | 0.92529245 | 0.00061738 |
| SetSize200 | 80% Sparsity | 70% Markers | GSDensity | 0.87661838 | 0.01164277 | 0.94862919 | 0.00213814 | 0.97090846 | 0.00116897 |
| SetSize200 | 80% Sparsity | 70% Markers | GESSO | 0.99999975 | 4.3995E-07 | 0.99986153 | 0.00023535 | 0.99991931 | 0.00013716 |
| SetSize200 | 90% Sparsity | 15% Markers | AUCell | 0.55894405 | 0.02043189 | 0.85793409 | 0 | 0.92353555 | 0 |
| SetSize200 | 90% Sparsity | 15% Markers | GSVA | 0.51361112 | 0.00817161 | 0.85793409 | 0 | 0.92353555 | 0 |
| SetSize200 | 90% Sparsity | 15% Markers | ssGSEA | 0.5080511 | 0.00832111 | 0.85793409 | 0 | 0.92353555 | 0 |
| SetSize200 | 90% Sparsity | 15% Markers | GSDensity | 0.64221204 | 0.05245734 | 0.85793409 | 0 | 0.92353555 | 0 |
| SetSize200 | 90% Sparsity | 15% Markers | GESSO | 0.9135918 | 0.01318429 | 0.92727776 | 0.00632337 | 0.95865641 | 0.003506 |
| SetSize200 | 90% Sparsity | 30% Markers | AUCell | 0.59572802 | 0.02188495 | 0.85793409 | 0.00013055 | 0.92353327 | 7.0439E-05 |
| SetSize200 | 90% Sparsity | 30% Markers | GSVA | 0.51175839 | 0.00787585 | 0.85793409 | 0 | 0.92353555 | 0 |
| SetSize200 | 90% Sparsity | 30% Markers | ssGSEA | 0.52079654 | 0.01272339 | 0.85793409 | 0 | 0.92353555 | 0 |
| SetSize200 | 90% Sparsity | 30% Markers | GSDensity | 0.64982961 | 0.03725336 | 0.87776239 | 0.01141535 | 0.93349814 | 0.00580439 |
| SetSize200 | 90% Sparsity | 30% Markers | GESSO | 0.97680513 | 0.00615067 | 0.96532816 | 0.00445163 | 0.97998558 | 0.00253925 |
| SetSize200 | 90% Sparsity | 50% Markers | AUCell | 0.79428141 | 0.02510703 | 0.87308225 | 0.00622265 | 0.93004969 | 0.00301888 |
| SetSize200 | 90% Sparsity | 50% Markers | GSVA | 0.54843287 | 0.03083663 | 0.85793409 | 0 | 0.92353555 | 0 |
| SetSize200 | 90% Sparsity | 50% Markers | ssGSEA | 0.56363019 | 0.01270355 | 0.85793409 | 0 | 0.92353555 | 0 |
| SetSize200 | 90% Sparsity | 50% Markers | GSDensity | 0.81048233 | 0.01698829 | 0.92863473 | 0.00590939 | 0.95997505 | 0.0032075 |
| SetSize200 | 90% Sparsity | 50% Markers | GESSO | 0.99612251 | 0.00107733 | 0.98532263 | 0.00286014 | 0.99147335 | 0.0016585 |
| SetSize200 | 90% Sparsity | 70% Markers | AUCell | 0.93613106 | 0.01350128 | 0.92669621 | 0.00704959 | 0.95812718 | 0.00390369 |
| SetSize200 | 90% Sparsity | 70% Markers | GSVA | 0.65744283 | 0.04466107 | 0.85754639 | 0.00041693 | 0.92329249 | 0.00025224 |
| SetSize200 | 90% Sparsity | 70% Markers | ssGSEA | 0.6117207 | 0.01374363 | 0.85793409 | 0.00013055 | 0.92353327 | 7.0439E-05 |
| SetSize200 | 90% Sparsity | 70% Markers | GSDensity | 0.90536157 | 0.01194141 | 0.94846303 | 0.00367959 | 0.97068916 | 0.00204411 |
| SetSize200 | 90% Sparsity | 70% Markers | GESSO | 0.99892391 | 0.0008351 | 0.99329826 | 0.00198757 | 0.9960994 | 0.00115586 |

**Supplementary Table 4. Dataset-specific preprocessing steps and parameter settings for GESSO computation.** The 1^st^ column indicates the platform of dataset; the 2^nd^ column indicates the specific name of the dataset we used in this work. The 3^rd^ column describes the preprocessing context for each dataset. The 4^th^ column shows the final number of spatial locations after filtering out low quality spots. The 5^th^ column shows the number of neighborhoods being used to build a spatial graph for each dataset. The 6^th^ column shows the spatial smoothing strength parameter for spatial regularization. The 7^th^ column shows the number of low-resolution partitions for high dimensional datasets.

| **Dataset Name** | **Specific Name** | **Preprocessing** | **# Spatial Locations (post-filter)** | **GESSO k** | **GESSO beta** | **GESSO low-res # partitions** |
| --- | --- | --- | --- | --- | --- | --- |
| 10x Visium human tumor datasets | BRCA | normalize log1p | N/A | 6 | 0.33 | N/A |
|  | CRC | normalize log1p | N/A | 6 | 0.33 | N/A |
|  | LUSC | normalize log1p | N/A | 6 | 0.33 | N/A |
|  | OVCA | normalize log1p | N/A | 6 | 0.33 | N/A |
|  | CESC | normalize log1p | N/A | 6 | 0.33 | N/A |
|  | PRAD | normalize log1p | N/A | 6 | 0.33 | N/A |
| Stereo-seq mouse dataset | E11.5 | none | N/A | 20 | 0.33 | 10 |
|  | E12.5 | none | N/A | 20 | 0.33 | 10 |
|  | E13.5 | none | N/A | 20 | 0.33 | 10 |
| Visium HD human colorectal cancer dataset | Normal | filter low-expression spots, normalize log1p | 319782 | 20 | 0.33 | 50 |
|  | Tumor | filter low-expression spots, normalize log1p | 460894 | 20 | 0.33 | 50 |
| Xenium Prime human lymph node dataset | N/A | filter low-expression spots, normalize log1p | 681601 | 20 | 0.33 | 200 |
| CosMx human non-small cell lung cancer dataset | section 10 | normalize log1p | N/A | 20 | 0.33 | 10 |
|  | section 16 | normalize log1p | N/A | 20 | 0.33 | 10 |
|  | section 22 | normalize log1p | N/A | 20 | 0.33 | 10 |
|  | section 28 | normalize log1p | N/A | 20 | 0.33 | 10 |

### Supplementary Notes

#### Supplementary Note 1: Methodological details for the GESSO method

##### 1.1 Key notation

$\boldsymbol{X}\in\mathbb{R}^{G\times N}$: Gene expression matrix with $G$ genes (rows) and $N$ spatial locations (columns).

$\boldsymbol{F}\in\mathbb{R}^{N\times2}$: Spatial location matrix with $N$ spatial locations (rows) and associated $x$ and $y$ coordinates (columns).

$l$: A predefined gene set consisting of $G_{l}$ genes.

$\boldsymbol{X}_{l}\in\mathbb{R}^{G_{l}\times N}$: Row-submatrix of $\boldsymbol{X}$ consisting of the $G_{l}$ genes in gene set $l$.

$\boldsymbol{L}\mathbb{\in R}^{N\times N}$: Graph Laplacian matrix constructed via *k*-nearest neighbors.

$\boldsymbol{q}\in\mathbb{R}^{N}$: Gene set activity score (GAS) vector (one GAS per spatial location).

$\boldsymbol{u}\in\mathbb{R}^{G_{l}}$: Metagene vector for gene set $l$ (one weight per gene). The metagene weight corresponding to a specific gene in $l$ can be interpreted as that gene’s contribution to the GESSO-computed GAS vector $\boldsymbol{q}$.

All other notation is explained when introduced. Bolded and capitalized letters denote matrices, bolded and lower-case letters denote vectors, and non-bolded letters denote scalars.

##### 1.2 GESSO optimization framework

For an arbitrary gene set $l$with $G_{l}$ genes, we extract a row-submatrix of $\boldsymbol{X}$, $\boldsymbol{X}_{l}\in\mathbb{R}^{G_{l}\times N}$ that contains only the $G_{l}$ genes. Then, we employ graph-Laplacian-regularized rank-one matrix decomposition of $\boldsymbol{X}_{l}$ into (1) the first left singular vector $\boldsymbol{u}\in\mathbb{R}^{G_{l}}$, and (2) the first right singular vector $\boldsymbol{q}\in\mathbb{R}^{N}$ with respect to a spatial smoothing constraint generated from the location matrix $\boldsymbol{F}$. We refer to the $G_{l}$-vector $\boldsymbol{u}$ as the “metagene” corresponding to the $N$-vector of gene set activity scores $\boldsymbol{q}$. The detailed derivation is in the following.

Using the location matrix $\boldsymbol{F}$, we construct a graph Laplacian$\boldsymbol{L}$via $k$-nearest neighbors. Specifically, we define an adjacency matrix $\boldsymbol{A}$ where $\boldsymbol{A}_{ij}=1$ if locations $i$ and $j$ are neighbors, and $0$ otherwise. The degree matrix $\boldsymbol{D}$ is diagonal, with $\boldsymbol{D}_{ii}$ as the sum of row $i$ in $\boldsymbol{A}$. The Laplacian is then given by $\boldsymbol{L}=\boldsymbol{D}-\boldsymbol{A}$.

We obtain the gene expression submatrix $\boldsymbol{X}_{l}$ from $\boldsymbol{X}$ and standard scale $\boldsymbol{X}_{l}$ along its rows. Given $\boldsymbol{X}_{l}$and the graph Laplacian $\boldsymbol{L}$*,* we proceed with rank-one approximation by solving the following optimization problem:

$$\begin{aligned} \boldsymbol{u}^{\star},\boldsymbol{q}^{\star}=\text{argmin}_{\boldsymbol{u},\boldsymbol{q}}\left\{ \left| |\boldsymbol{X}_{l}\boldsymbol{-}\boldsymbol{uq}^{\top} \right|\left. \right|_{F}^{2}+\alpha\text{Tr}\left( \boldsymbol{q}^{\top}\boldsymbol{Lq} \right) \right\},\text{ such that }\boldsymbol{q}^{\top}\boldsymbol{q}=1.\#\left( 1 \right) \end{aligned}$$

Jiang et al. (2013) derived the closed-form solution for this optimization problem^1^. Let $\lambda_{n}$ be the largest eigenvalue of ${\boldsymbol{X}_{l}}^{\boldsymbol{\top}}\boldsymbol{X}_{l}$, $\xi_{n}$ be the largest eigenvalue of $\mathbf{L}$, and define $\alpha=\frac{\beta\lambda_{n}}{\left( 1-\beta\right)\xi_{n}}.$ We introduce $\boldsymbol{G}_{\beta}=\left( 1-\beta\right)\left( I-\frac{{\boldsymbol{X}_{l}}^{\top}\boldsymbol{X}_{l}}{\lambda_{n}} \right)+\frac{\beta}{\xi_{n}}\boldsymbol{L}$, and $\boldsymbol{v}_{1}$ is the eigenvector corresponding to the smallest eigenvalue of $\mathbf{G}_{\beta}$. The solution is:

$$\begin{aligned} \boldsymbol{q}^{\star}=\boldsymbol{v}_{1},\boldsymbol{u}^{\star}=\boldsymbol{X}_{l} \boldsymbol{q}^{\star}.\#\left( 2 \right) \end{aligned}$$

For completeness, we present a detailed derivation of Equation 2, the closed-form solution to Equation 1.

First, we fix $\boldsymbol{q}$ and solve for optimal $\boldsymbol{u}$. Taking the derivative of the objective with respect to $\mathbf{u}$ and setting it to zero yields:

$$\frac{\partial}{\partial\boldsymbol{u}}\left. ||\boldsymbol{X}_{l}-\boldsymbol{u}\boldsymbol{q}^{\top} \right.{||}_{F}^{2}=-2\left( \boldsymbol{X}_{l}-\boldsymbol{u}\boldsymbol{q}^{\top} \right)\boldsymbol{q}=0$$

$$\Rightarrow-2\boldsymbol{X}^{'}\boldsymbol{q}+2\boldsymbol{u}\boldsymbol{q}^{\top}\boldsymbol{q}=0$$

$$\Rightarrow\boldsymbol{u}\boldsymbol{q}^{\top}\boldsymbol{q}=\boldsymbol{X}_{l}\boldsymbol{q}$$

$$\begin{aligned} \Rightarrow\boldsymbol{u}^{\star}=\boldsymbol{X}_{l}\boldsymbol{q}.\#\left( 3 \right) \end{aligned}$$

Then, we substitute this optimal $\mathbf{u}^{\boldsymbol{\star}}$ (Equation 3) back into the objective function and simplify, noting that $\mathbf{q}^{\top}\mathbf{q}=1$:

$$\left. ||\boldsymbol{X}_{l} -\boldsymbol{X}_{l} \boldsymbol{q}\boldsymbol{q}^{\top} \right.{||}_{F}^{2}+\alpha\text{Tr}\left( \boldsymbol{q}^{\top}\boldsymbol{Lq} \right)=\text{Tr}\left[ \left( \boldsymbol{X}_{l} -\boldsymbol{X}_{l} \boldsymbol{q}\boldsymbol{q}^{\top} \right)^{\top}\left( \boldsymbol{X}_{l} -\boldsymbol{X}_{l} \boldsymbol{q}\boldsymbol{q}^{\top} \right) \right]+\alpha\text{Tr}\left( \boldsymbol{q}^{\top}\boldsymbol{Lq} \right)$$

$$\text{=}\text{Tr}\left( {\boldsymbol{X}_{l}}^{\top}\boldsymbol{X}_{l} \right)-\text{Tr}\left( {\boldsymbol{X}_{l}}^{\top}\boldsymbol{X}_{l} \boldsymbol{q}\boldsymbol{q}^{\top} \right)-\text{Tr}\left( \boldsymbol{q}\boldsymbol{q}^{\boldsymbol{\top}}{\boldsymbol{X}_{l}}^{\top}\boldsymbol{X}_{l} \right)+\text{Tr}\left( \boldsymbol{q}\boldsymbol{q}^{\boldsymbol{\top}}{\boldsymbol{X}_{l}}^{\top}\boldsymbol{X}_{l}\boldsymbol{q}\boldsymbol{q}^{\top} \right)+\alpha\text{Tr}\left( \boldsymbol{q}^{\top}\boldsymbol{Lq} \right)$$

$$\begin{aligned} =\text{Tr}\left( {\boldsymbol{X}_{l}}^{\top}\boldsymbol{X}_{l} \right)-\text{Tr}\left( \boldsymbol{q}^{\boldsymbol{\top}}{\boldsymbol{X}_{l}}^{\top}\boldsymbol{X}_{l}\boldsymbol{q} \right)+\alpha\text{Tr}\left( \boldsymbol{q}^{\top}\boldsymbol{Lq} \right).\#\left( 4 \right) \end{aligned}$$

The first term of Equation 4 does not involve $\mathbf{q}$, so we only need to consider the two remaining terms. Our optimization problem becomes:

$$\begin{aligned} \boldsymbol{q}^{\star}=\text{argmin}_{\boldsymbol{q}}\left\{ \text{Tr}\left[ \boldsymbol{q}^{\top}\left( -{\boldsymbol{X}_{l}}^{\top}\boldsymbol{X}_{l}+\alpha\boldsymbol{L} \right)\boldsymbol{q} \right] \right\},\text{ such that }\boldsymbol{q}^{\top}\boldsymbol{q}=1\mathbf{.}\#\left( 5 \right) \end{aligned}$$

Let $\boldsymbol{G}_{\alpha}=-{\boldsymbol{X}_{l}}^{\top}\boldsymbol{X}_{l}+\alpha\boldsymbol{L}$. We define the Lagrangian as $L\left( \boldsymbol{q},\lambda\right)=\text{Tr}\left( \boldsymbol{q}^{\boldsymbol{\top}}\boldsymbol{G}_{\alpha}\boldsymbol{q} \right)-\lambda\left( \boldsymbol{q}^{\boldsymbol{\top}}\boldsymbol{q}-1 \right)$*,* where $\lambda$ is the Lagrange multiplier. Taking the derivative with respect to $\mathbf{q}$ and setting the result to $0$, we find:

$$\frac{\partial L}{\partial\boldsymbol{q}}=2\boldsymbol{G}_{\alpha}\boldsymbol{q}-2\lambda\boldsymbol{q}=0$$

$$\Rightarrow\boldsymbol{G}_{\alpha}\boldsymbol{q}=\lambda\boldsymbol{q}.$$

This is an eigenvalue equation; $\boldsymbol{q}$ must therefore be an eigenvector of $\boldsymbol{G}_{\alpha}$, where $\lambda$ is its corresponding eigenvalue. Returning to our objective, we find $\text{Tr}\left( \boldsymbol{q}^{\top}\boldsymbol{G}_{\alpha}\boldsymbol{q} \right)=\text{Tr}\left( \boldsymbol{q}^{\top}\lambda\boldsymbol{q} \right)=\lambda$. Naturally, to minimize the objective, $\lambda$ must be the smallest eigenvalue of $\mathbf{G}_{\alpha}$. Let $\mathbf{v}_{1}$ denote the eigenvector corresponding to the smallest eigenvalue. We arrive at our closed-form solution to Equation 1:

$$\boldsymbol{q}^{\star}=\boldsymbol{v}_{1},\boldsymbol{u}^{\star}=\boldsymbol{X}_{l}\boldsymbol{q}^{\star}.$$

We follow previous study^1^ to reparametrize Equation 1 to allow for a more interpretable regularization term scaling constant. We adopt this reparameterization.

Let $\lambda_{n}$ denote the largest eigenvalue of ${\boldsymbol{X}_{l}}^{\top}\boldsymbol{X}_{l}$. Let $\xi_{n}$ denote the largest eigenvalue of $\boldsymbol{L}$. Let $\alpha=\frac{\beta\lambda_{n}}{\left( 1-\beta\right)\xi_{n}}$. Substituting this value in for $\alpha$ in Equation 5, we obtain the following:

$$\begin{aligned} \boldsymbol{q}^{\star}=\text{argmin}_{\boldsymbol{q}}\left\{ \text{Tr}\left[ \boldsymbol{q}^{\top}\left( -{\boldsymbol{X}_{l}}^{\top}\boldsymbol{X}_{l}+\left( \frac{\beta\lambda_{n}}{\left( 1-\beta\right)\xi_{n}} \right)\boldsymbol{L} \right)\boldsymbol{q} \right] \right\},\text{ such that }\boldsymbol{q}^{\top}\boldsymbol{q}=1\boldsymbol{.}\#\left( 6 \right) \end{aligned}$$

Following Jiang et al. (2013) ^1^, Equation 6 is equivalent to:

$$\begin{aligned} \boldsymbol{q}^{\star}=\text{argmin}_{\boldsymbol{q}}\left\{ \text{Tr}\left[ \boldsymbol{q}^{\top}\left( \left( 1-\beta\right)\left( \boldsymbol{I}-\frac{{\boldsymbol{X}_{l}}^{\top}\boldsymbol{X}_{l}}{\lambda_{n}} \right)+\frac{\beta}{\xi_{n}}\boldsymbol{L} \right)\boldsymbol{q} \right] \right\},\text{ such that }\boldsymbol{q}^{\top}\boldsymbol{q}=1\boldsymbol{.}\#\left( 7 \right) \end{aligned}$$

To verify, let

$$\boldsymbol{G}_{\boldsymbol{1}}=-{\boldsymbol{X}_{l}}^{\top}\boldsymbol{X}_{l}+\frac{\beta\lambda_{n}}{\left( 1-\beta\right)\xi_{n}}\boldsymbol{L},$$

$$\boldsymbol{G}_{\beta}=\left( 1-\beta\right)\left( I-\frac{{\boldsymbol{X}_{l}}^{\top}\boldsymbol{X}_{l}}{\lambda_{n}} \right)+\frac{\beta}{\xi_{n}}\boldsymbol{L}.$$

A straightforward calculation shows that

$$\boldsymbol{G}_{\beta}=\left( 1-\beta\right)\boldsymbol{I}+\frac{1-\beta}{\lambda_{n}}\boldsymbol{G}_{1}.$$

We find that $\boldsymbol{G}_{1}$ and $\boldsymbol{G}_{\beta}$ differ only by a positive scalar multiple and a shift by the identity matrix. Since multiplying by a positive scalar and adding a constant multiple of the identity does not change the eigenvectors, both matrices produce the same minimizer $\boldsymbol{q}^{\star}$ when we solve $\min_{|\boldsymbol{q}|=1} \boldsymbol{q}^{\top}\boldsymbol{G}_{1}\boldsymbol{q}$ *or* $\min_{|\boldsymbol{q}|=1} \boldsymbol{q}^{\top}\boldsymbol{G}_{\beta}\boldsymbol{q}$. Therefore, Equation 6 and Equation 7 are equivalent.

Our solution to the reparametrized optimization problem as defined in Equation 7 is identical to Equation 2 (our solution to Equation 1) but with $\boldsymbol{v}_{1}$ representing the eigenvector corresponding to the smallest eigenvalue of $\boldsymbol{G}_{\beta}$. Note that $\boldsymbol{G}_{\beta}$ is positive semidefinite.

We interpret the $G_{l}$-vector $\boldsymbol{u}^{\star}$ as the metagene (one importance weight for each gene in the pathway) and the $N$-vector $\mathbf{q}^{\star}$ as the gene set activity scores (one score for each spatial location). To enhance metagene interpretability, we normalize $\boldsymbol{u}^{\star}$ by scaling it by $1/\boldsymbol{||u}^{\star}||$*.* We scale $\boldsymbol{q}^{\star}$ by $\boldsymbol{||u}^{\star}||$ accordingly. One can therefore interpret the metagene $\boldsymbol{u}^{\star}$ as a single overall gene (that is, $\boldsymbol{u}^{\star}$ has magnitude $1$) representative of all genes in the pathway.

##### 1.3 Gene set activity score sign determination

Matrix decomposition methods, including the graph-regularized principal component analysis method employed by GESSO, require a heuristic to deterministically assign signs. Concretely, if $\boldsymbol{u}^{\star}$ and $\boldsymbol{q}^{\star}$ minimize $\left| |\boldsymbol{X}_{l}\boldsymbol{-}\boldsymbol{uq}^{\top} \right|\left. \right|_{\boldsymbol{F}}^{\boldsymbol{2}}+\alpha\text{Tr}\left( \boldsymbol{q}^{\top}\boldsymbol{Lq} \right),$then so do $\boldsymbol{-u}^{\star} a$nd $\boldsymbol{-}\boldsymbol{q}^{\star}$.

We adopt a sign assignment heuristic that aligns the direction of $\boldsymbol{q}^{\star}$ with a gene set activity proxy. We define the proxy as the mean gene expression across all genes in the pathway at each spatial location. Formally, let

$$\boldsymbol{p}=\frac{1}{G_{l}}\sum_{g=1}^{G_{l}} {\boldsymbol{(X}_{l}\boldsymbol{)}}_{g}.$$

${\boldsymbol{(X}_{l}\boldsymbol{)}}_{g}$ denotes the $g$-th row (an $N$-vector) of matrix $\boldsymbol{X}_{l}$. The $N$-vector $\boldsymbol{p}$ serves as a coarse summary of gene set activity. To resolve the sign ambiguity, we compute the Pearson correlation between the raw activity scores $\boldsymbol{q}^{\star}$ and the proxy vector $\boldsymbol{p}$. If this correlation is negative or undefined, we flip the sign of both $\boldsymbol{u}^{\star}$ and $\boldsymbol{q}^{\star}$ to ensure that the direction of the gene set activity scores is positively aligned with the overall expression level of the pathway. This step ensures consistent biological interpretation of gene set activity across datasets, improving the robustness and comparability of downstream analyses.

##### 1.4 Low-resolution analysis of partitioned datasets

The GESSO method exhibits at least quadratic scaling in both time and space complexity with respect to $N$, the number of spatial locations. For large spatial transcriptomics datasets (such as those exceeding $20,000$ spots) this scaling behavior imposes significant computational demands, often rendering analysis infeasible on standard hardware with less than $100$ GB of random-access memory (RAM). To address this limitation, we introduce a low-resolution approximation strategy that enables efficient gene set activity inference on large datasets while preserving the spatial structure of the original sample.

Our approach begins by partitioning the full spatial domain into $p$ non-overlapping subsets, each of which retains the spatial characteristics of the tissue. For each subset, GESSO is run independently to compute gene set activity scores, effectively treating each subset as a down sampled, spatially coherent region. After scoring, all computed activity profiles are aggregated to recover pathway scores for all $N$ spatial locations.

To ensure that each partition captures representative spatial variation and avoids spatial bias, we implement a stratified $k$-means partitioning scheme. This method first clusters the tissue into $k$ spatial regions using standard $k$-means on the spatial coordinates. Then, within each cluster, we randomly redistribute the spatial locations across all $p$ partitions, ensuring that each partition receives a balanced sample from every spatial cluster. This stratification guarantees that each subset approximates the full spatial diversity of the tissue while reducing the number of locations processed per GESSO call. In this work, we set the number of $k$-means clusters, $k$, equal to the number of partitions, $p$.

This strategy reduces the memory and runtime burden of GESSO by limiting the number of locations analyzed per run, while preserving pathway resolution across the full tissue. In practice, we find that appropriate values of $p$ (e.g., $p=10$ for Stereo-seq, $p=50$ for Visium HD, and $p=200$ for Xenium Prime) substantially reduce computational overhead. A post-hoc merging step reconstructs the full-resolution spatial map of gene set activity from the independently processed subsets.

We evaluated the accuracy of the low-resolution GESSO approximation by comparing its gene set activity scores to those computed using the full-resolution method on 10x Visium human tumor datasets. We shall also refer to the full-resolution method as the “standard” method. Across experiments, the low-resolution approach produced pathway scores that closely matched those from the standard method, demonstrating that computational efficiency can be gained without sacrificing accuracy. Moreover, we found that the use of stratified $k$-means for partitioning outperformed random partitioning, yielding more faithful approximations of full-resolution scores. Detailed benchmarking results are provided in Supplementary Note 2.

##### 1.5 Permutation-based hypothesis testing for elevated pathway activity detection

Consider an arbitrary pathway composed of $G_{l}$ genes, selected from a total of $G$ genes in the full gene expression dataset $\boldsymbol{X}$. The gene set activity score for this gene set is computed and denoted by $\boldsymbol{q}$, an $N$-dimensional vector. To evaluate the significance of $\boldsymbol{q}$, we perform a permutation-based hypothesis test: $G_{l}$ genes are randomly selected from the full set $n_{\text{control}}$ times, yielding $n_{\text{control}}$ control GASs. These are aggregated into an $N\times n_{\text{control}}$ matrix $\boldsymbol{P}$, where each column represents a control score vector.

We treat $\boldsymbol{P}$ as a sample from the null distribution of GASs in $N$-dimensional space. To estimate the significance of $\boldsymbol{q}$, we compare its values to the corresponding rows of $\boldsymbol{P}$. For each spot $i$, let $\boldsymbol{P}_{i}$denote the $i$-th row of $\boldsymbol{P}$, containing control scores at that spot. Let $c$ be the number of values in $\boldsymbol{P}_{i}$ that exceed$\boldsymbol{q}_{i}$*.* We estimate the p-value for spot $i$ as $\frac{c}{n_{\text{control}}}$, representing the probability under the null that the observed score $\boldsymbol{q}_{i}$is not greater than expected by chance.

Repeating this procedure across all $N$ spots yield a vector of p-values, one for each spot. Application of a multiple testing correction, such as the Bonferroni method, is left to the user’s discretion.

#### Supplementary Note 2: Experimental validation of the low-resolution GESSO method

To enable GESSO to be scalable to large datasets, we developed a low-resolution computation approach (see Supplementary Note 1). Briefly, we partition the spatial locations into subsets of approximately equal size and apply the standard GESSO procedure to each subset independently, yielding a GAS for every spatial location. In this Supplementary Note, we experimentally assess the accuracy of the low-resolution method relative to the standard GESSO approach. We also evaluate the performance of the proposed stratified k-means algorithm for generating spatially balanced partitions, comparing it to random partitioning. For both partitioning strategies (stratified k-means and random), we generated low-resolution GAS matrices using 5, 10, and 20 partitioned subsets for the 10x Visium human tumor datasets. “Partitioned subsets” refers to the $p$ parameter described in Supplementary Note 1; we also referred to $p$ as the “number of partitions” elsewhere. To conserve computational resources, we limited the experiment to the first 500 of the 8,729 total pathways, selected based on alphabetical order. We then quantified the root mean squared error (RMSE) and Pearson correlation between each flattened low-resolution GAS matrix and the flattened full-resolution GAS matrix produced by the standard GESSO method.

Our findings (Supplementary Figure S41) demonstrated that the low-resolution method consistently produced GAS matrices that closely matched the full-resolution results, with Pearson correlation coefficients exceeding 0.94 across all datasets for the $p=5$ setting. These results suggest that the low-resolution partitioning method offers a substantial gain in computational scalability while preserving high accuracy in GAS estimation relative to the standard GESSO method. As the number of partitions increased, the accuracy of the low-resolution GASs declined as expected, though Pearson correlation coefficients remained above 0.81 across all datasets, indicating a reasonable trade-off between resolution and computational efficiency. We also found that the stratified k-means partitioning approach yielded more accurate GASs ($p=20$) compared to random partitioning, achieving a greater Pearson correlation and lower RMSE on five of the six datasets.

#### Supplementary Note 3: Description on compared gene set scoring methods

We compared GESSO with four widely used gene set level activity scoring methods such as AUCell, GSVA, and ssGSEA and GSDensity. Below, we describe each method briefly:

- **ssGSEA (single-sample Gene Set Enrichment Analysis)**^2,3^: For a given sample, genes are ranked by expression, and an enrichment score is computed by walking down the ranked list and increasing a running-sum statistic when a gene in the set is encountered (and decreasing when it is not). The resulting score reflects how strongly the genes in the pathway are coordinately up- or down-regulated in that individual sample. This method, derived from the original GSEA algorithm, allows pathway analysis in settings where each sample must be evaluated independently (such as individual patients or single cells).
- **GSVA (Gene Set Variation Analysis)**^4^: GSVA is a non-parametric, unsupervised method that transforms the gene expression matrix into pathway space by estimating the enrichment of gene sets in each sample’s expression profile. It calculates a Kolmogorov-Smirnov-like statistic over the ranked expression values of a sample to assess the relative enrichment of a gene set, thereby providing a continuous gene set activity score. By evaluating the variation of gene set enrichment across samples, GSVA captures pathway-level differences in a sample population without requiring a phenotype label for each sample.
- **AUCell**^5^: AUCell determines whether a gene set is active in each individual cell (primarily designed for single-cell RNA-seq data) by computing the Area Under the Curve (AUC) of the recovery curve of gene-set members in the ranked expression list of that cell. In practice, genes are ranked by expression within a cell, and the fraction of the gene set present toward the top of this ranking is measured as an AUC score. A high AUC indicates that many genes of the set are among the top-expressed genes in the cell, signifying an active pathway. This method is robust to normalization and has been used to identify cell states or cell-specific pathway activities in single-cell analyses.
- **GSDensity**^6^: GSDensity is a recent pathway-centric method designed to analyze both single-cell and spatial transcriptomics data using graph-based modeling. It constructs a nearest-neighbor graph that includes both cells and genes embedded in a shared low-dimensional space, then applies a density-based scoring approach to assess gene set activity without requiring prior cell clustering. By evaluating the coordinated expression of pathway genes across this graph, GSDensity can detect subtle subpopulations of cells and uncover cell-pathway associations that may be missed by other methods. In spatial transcriptomics, GSDensity identifies spatially relevant pathways by comparing kernel-density estimates of gene set activity across physical tissue coordinates. While the initial scoring is spatially agnostic, the final spatial enrichment analysis incorporates physical distances between spots to highlight pathways localized to specific regions or tissue structures.

#### Supplementary Note 4: Data preprocessing, gene set selection, and dataset details

##### Data preprocessing

For all datasets, spatial locations with fewer than 50 detected transcripts were removed. Raw count matrices were then normalized by total counts per location, scaled to the median total count across all locations, and log1p transformed. Datasets that were already normalized in their original publications (e.g., Stereo-seq mouse embryo) were used as provided. More details on the preprocessing for each dataset are summarized in Supplementary Table 4.

##### Gene set selection

All mouse and human gene sets analyzed in this study were collected from MsigDB database^2,7^ using the R package *msigdbr()*. For the 10x Visium human tumor datasets, we retained pathways with at least 10 overlapping genes across all six tumor datasets, yielding a total of 8,729 pathways. For the Stereo-seq mouse embryo data, we selected 313 gene sets by identifying those whose names contained annotated organ terms. For the Visium HD human CRC, and CosMx human NSCLC datasets where cell type labels were provided by their original studies, and 10x 5K human lymph node where we manually annotated GCs, we first performed fast gene set enrichment analysis^8^ (fGSEA) following <https://crazyhottommy.github.io/scRNA-seq-workshop-Fall-2019/scRNAseq_workshop_3.html> to identify cell-type-associated gene sets. We then selected the top 20 gene sets per cell type based on adjusted p-values. Additionally, for the lymph node and NSCLC datasets, we manually included gene sets containing “lymphnode” or “EMT” (Epithelia-Mesenchymal Transition) in their names. This procedure resulted in 97 gene sets for the CRC data, 150 for the lymph node data, and 151 for the NSCLC data.

##### Spatial transcriptomics dataset details and availability

###### 10x Visium human tumor datasets

We obtained spatial transcriptomics data for six cancer types—breast ductal carcinoma (BRCA), colorectal cancer (CRC), lung squamous cell carcinoma (LUSC), ovarian carcinoma (OVCA), cervical cancer (CESC) and prostate adenocarcinoma (PRAD)—from the 10x Genomics platform (<https://www.10xgenomics.com/datasets>). All tumor tissue samples were Formalin-Fixed Paraffin-Embedded (FFPE) samples. The number of spatial locations measured varied across cancer types: 2,518 in BRCA, 2,660 in CRC, 3,858 in LUSC, 3,455 in OVCA, 2,781 in CESC, and 4,371 in PRAD. The number of genes measured was consistent across most samples, with 17,943 genes in BRCA, CRC, OVCA, CESC, and PRAD, and 18,085 genes in LUSC. For the BRCA dataset (<https://www.10xgenomics.com/datasets/human-breast-cancer-ductal-carcinoma-in-situ-invasive-carcinoma-ffpe-1-standard-1-3-0>), an expert-annotated H&E image was provided. These annotations were used to evaluate the biological relevance of the GASs by testing their ability to recover known tissue regions.

###### Stereo-seq mouse embryo dataset

We downloaded mouse embryo SRT data^9^ from STOmics (<https://db.cngb.org/stomics/mosta/download/>). The data originated from three mouse embryonic development stages: E11.5, E12.5, and E13.5. The number of measured spatial locations varies across stages, with 30,124 in E11.5, 51,365 in E12.5, and 77,369 in E13.5. Each stage included expression measurements for 23,761 genes and provided anatomical organ region annotations.

###### Visium HD human colorectal cancer dataset

We obtained human colorectal cancer spatial transcriptomics data from 10x Genomics Platform^33^ (<https://www.10xgenomics.com/platforms/visium/product-family/dataset-human-crc>) . There were two datasets, a tumor tissue dataset (54,1968 locations) and an adjacent normal tissue (43,5773 locations) dataset, with expression data for 18,035 genes. For the tumor data, cell types label are provided in (<https://github.com/10XGenomics/HumanColonCancer_VisiumHD/tree/main/MetaData>).

###### Xenium Prime human lymph node dataset

We acquired human lymph node spatial transcriptomics data from 10x Genomics (<https://www.10xgenomics.com/datasets/preview-data-xenium-prime-gene-expression>). The dataset contained the expression quantification of 11,094 genes across 708,983 cells. We manually annotated germinal centers based on morphological features and staining patterns from protein-stained (Xenium Morphology) tissue images.

###### CosMx human non-small cell lung cancer dataset

We obtained 3D human non-small cell lung cancer data^10^ from Zenodo (<https://zenodo.org/records/15240431>). The sample comprises 55,339 cells and 960 measured genes. This dataset also included niche-level labels and tumor pseudo time information.
